# SPP1+ Microglia Are Associated with Neuroimmune Rewiring and Glutamatergic Neuronal Injury in ART-Suppressed People Living with HIV

**DOI:** 10.64898/2026.09.04.749241

**Authors:** Ciniso S. Shabangu, Hongjie Chen, Ashokkumar Manickam, Nikesh Katuwal, Jacob B. Lovins, Edward P. Browne, Sara Gianella, Antoine Chaillon, David M. Margolis, Yuyang Tang, Guochun Jiang

## Abstract

Antiretroviral therapy (ART) effectively suppresses systemic HIV replication but does not eradicate viral reservoirs in the brain, where their identity and contribution to neurological injury remain poorly defined. Using bulk, single-cell, and single-nucleus transcriptomics of postmortem human brain tissue, we identify an activated SPP1⁺ microglial population that expands 5.8-fold during ART and serves as the primary central nervous system (CNS) reservoir, preferentially harboring HIV transcripts. These reservoir microglia adopt a distinct, immune-evasive reprogramming state marked by chronic type I interferon signaling and inflammasome activation, which we recapitulate in primary human microglia via prolonged interferon-β exposure. We show that viral transcription in ART-suppressed brains is dominated by nef, which may sustain the viral reservoir by disrupting host HLA-A and HLA-E presentation machinery. This persistent, immune-evasive state is associated with TREM2-C1QC complement-mediated synaptic pruning and severe DNA damage response dysregulation, culminating in a profound loss of VGLUT1⁺ and GAD67⁺ glutamatergic neurons that persists despite viral suppression. Our findings establish SPP1⁺ microglia as an active, pathogenic CNS reservoir, identifying the SPP1, MHC-I, and complement pathways as therapeutic targets to eliminate viral persistence and reverse HIV-associated neurocognitive dysfunction.

## Introduction

Brain microglia (MG) are long-lived, self-renewing myeloid cells that serve as primary immune sentinels of the central nervous system (CNS), making them well suited to sustain chronic viral infection. Unlike rapidly turning over peripheral immune cells, microglia persist for decades within an immune-privileged environment partially shielded from systemic antiviral immune responses and antiretroviral therapy (ART) penetration. Numerous studies have detected HIV DNA and RNA in the brains of ART-suppressed people with HIV (PWH) (*1–4*). While viral infection occurs in astrocytes, pericytes, and CNS T cells (*5–7*), we and others have demonstrated that replication-competent HIV persists primarily within myeloid cells despite effective ART (*4, 8, 9*). Over 95% of these infected myeloid cells express TMEM119, establishing microglia as a stable, epigenetically controlled viral reservoir in the brains of PWH on durable ART (*4, 10*).

Notably, the number of MG isolated from PWH on ART was higher than in controls without HIV (*4*), indicating ongoing immune activation despite virologic suppression. This is supported by chronic neuroinflammation, marked by persistent type I interferon (IFN-I) signaling, one of the hallmarks of HIV-associated neurocognitive disorders (HAND) that affects 30%–50% of individuals on effective ART (*11, 12*). These observations suggest a pathological paradox where infected microglia simultaneously propagate inflammatory signals that drive neurodegeneration while successfully evading the immune responses required to clear the reservoir (*3, 13–15*). However, the precise molecular mechanisms by which persistent microglial infection orchestrates this dual phenotype of chronic neuroinflammation and immunological stealth remain poorly understood.

Here, we resolve these mechanisms using rapid research autopsies of brain tissues and freshly isolated microglia from ART-suppressed PWH, viremic individuals, and uninfected controls. Integrating bulk, single-cell, and single-nucleus transcriptomics, CellChat-based cell-cell communication modeling, and structural host-pathogen protein interaction analysis, we identify a distinct SPP1⁺ activated microglial subset that serves as the primary CNS viral reservoir. This population undergoes profound transcriptional reprogramming, acting as a “broadcasting hub” that drives toxic TREM2-C1QC-C3 complement opsonization and a subsequent neuronal DNA damage response (DDR) inversion. Concurrently, these cells insulate themselves from clearance via a precise Nef-mediated trafficking arrest of HLA-A and HLA-E. Together, our findings map a self-perpetuating cycle of neuroinflammation and lineage-specific neuronal decay that persists despite systemic viral suppression.

## Results

### Predominant SPP1 Signaling in The Brain MG in ART-Suppressed PWH 1

We began our investigation by analyzing postmortem brain tissues from uninfected, viremic, and ART-suppressed individuals to characterize microglial populations and identify condition-specific signaling pathways. Expression of HIV in brain MG from ART-suppressed patients (n=5) was confirmed by total HIV DNA quantification (**Fig. 1A and *Supplementary Table 1***), establishing the presence of persistent viral genomes in the CNS despite suppressive therapy.

**Figure 1.**
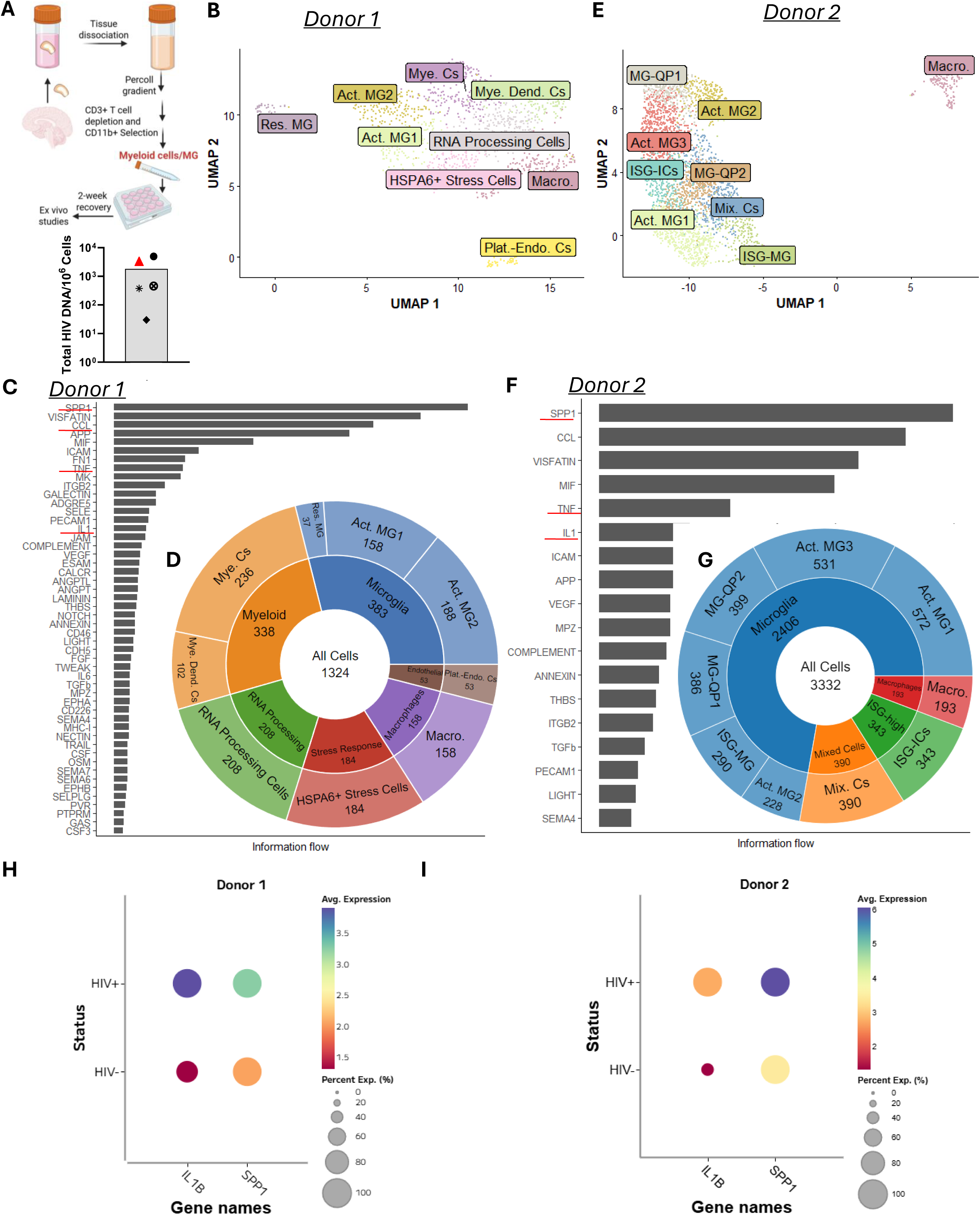
Predominant SPP1 signaling in the brains of ART-suppressed PWH. **(A)** Upper panel: Isolation of brain MG from PWH. Lower panel: quantification of total proviral HIV DNA in the isolated human brain MG on suppressive ART (n=5). **(B, E)** UMAP projections of single-nucleus transcriptome from donor 1 and donor 2, showing annotated brain cell populations including activated MG (Act.MG1/2/3), resident MG (Res.MG), interferon-stimulated MG (ISG-MG, ISG-ICs), astrocytes (A1 reactive, A2 reactive, quiescent), macrophages (Macro.), myeloid cells (Mye.Cs), myeloid dendritic cells (Mye.Dend.Cs), mixed immune cells (Mix.Cs), and platelet–endothelial cells (Plat.– Endo.Cs). **(C, F)** CellChat communication networks illustrating aberrant SPP1 signaling in donor 1 and donor 2 brains. **(D, G)** Distribution of cell types in donor 1 **(D)** and donor 2 **(G)** in ART-suppressed HIV⁺ brain tissues. (**H, I**) Expression of SPP1 and IL-1β in HIV⁺ and HIV⁻ CNS cells Donor 1 and 2. SPP1 and IL-1β are upregulated in HIV⁺ compared to HIV⁻ cells.

Single-cell RNA sequencing (scRNA-seq) revealed diverse MG subsets defined by canonical markers including HEXB and P2RY12 (**Supplementary Fig. 1A**), with MG subpopulations (**Fig. 1B, D**). To understand intercellular communication networks within the ART-suppressed CNS microenvironment, we applied CellChat analysis, which predicts ligand-receptor interactions based on expression patterns across cell types. SPP1 (osteopontin) signaling dominated the predicted communication landscape (**Fig. 1C**), with significant enrichment in SPP1-CD44 interactions with MG subsets (**Supplementary Fig. 1B**). Elevated SPP1 expression concentrated in activated microglia subsets (Act. MG1: log normalization (lognorm): 0.92, Act. MG2: lognorm: 0.11) in donor 1 (**Fig. 1D, Supplementary Fig. 1C**). Within the activated MG population, HIV^+^ MG exhibited markedly higher SPP1 expression (lognorm: 3.26) compared to HIV^-^ bystander cells (lognorm: 2.06) (**Fig. 1H**), suggesting that SPP1 expression is associated with HIV persistence.

To validate these findings in an independent dataset, we analyzed ART-suppressed donor 2, previously documented to harbor latently infected, replication-competent HIV in brain MG (*4*) (**Fig. 1E, G**). Enriched CD11b^+^ MG profiling provided higher resolution of microglial subsets, identifying activated MG1 and MG3 (marker genes: HEXB, APOE), interferon-stimulated MG (ISG-MG: HEXB, APOE, ISG15), and macrophages (LYZ, CD68) (**Supplementary Fig. 2A**). CellChat confirmed high SPP1 signaling (**Fig. 1F**) across MG subsets. Elevated expression of SPP1 was observed in activated MG subsets (Act. MG1: lognorm: 0.59, Act. MG2: lognorm: 0.74, ISG-MG: lognorm: 0.75) (**Supplementary Fig. 2C**). Critically, as in donor 1, HIV^+^ activated MG similarly showed elevated SPP1 expression (lognorm: 6.07) compared to HIV^-^ bystander cells (lognorm: 3.47) (**Fig. 1I**), replicating the pattern observed in donor 1. These convergent findings across independent donors and experimental platforms establish SPP1 as a defining feature of HIV^+^ activated MG in the ART-suppressed brain.

Beyond SPP1, both donors showed enrichment of CCL chemokines, TNF, and IL-1 pathways (**Fig. 1C and F**), indicating IL-1β neuroinflammatory signaling within the MG compartment despite suppressive ART. IL-1β was consistently upregulated in HIV^+^ activated MG (D1:(HIV^+^/HIV^-^)3.91/1.32; D2:2.77/1.34) (**Fig. 1H and I**).

SPP1 pathways are involved in negative regulation of intrinsic apoptosis, impaired DNA damage response, and disrupted cell survival signaling (**Supplementary Fig. 2D**), while CD44 was linked to axon guidance, neuronal projection, and synaptic regeneration. This dichotomy suggests that SPP1 expression in HIV^+^ MG promotes a survival-adapted, stress-resistant phenotype that may facilitate long-term viral persistence. Together, these results establish that SPP1 and IL-1β expression predominate in brain MG under ART, specifically within activated MG subsets, and that SPP1 expression is enriched in HIV^+^ cells.

We next sought to determine whether this transcriptional signature reflects a stable, condition-specific feature of the CNS microenvironment. We expanded our analysis to single nuclei (snRNA-seq), which included viremic (n=3) and uninfected (n=3) donors alongside ART-suppressed (n=4) individuals (**Supplementary Table 1**), providing a multi-platform framework that minimizes technical bias and captures cross-condition interactions.

### Aberrant SPP1 Signaling Marks Incomplete CNS Communication Recovery in PWH brains on ART

To assess cross-platform reproducibility within this expanded cohort, Donor 1 was included and re-analyzed using snRNA-seq, enabling direct cross-methodology validation within the same ART-suppressed individual. Multimodal integration using the Harmony algorithm (*17*) across a 2,000 highly variable gene space (27 PCA) revealed a highly consistent, batch-corrected cellular landscape of brain-resident populations across clinical conditions (**Fig. 2A-B; Supplementary Fig. 3A**). This integrated dataset allowed us to examine how HIV infection and ART suppression reshape the CNS.

**Figure 2.**
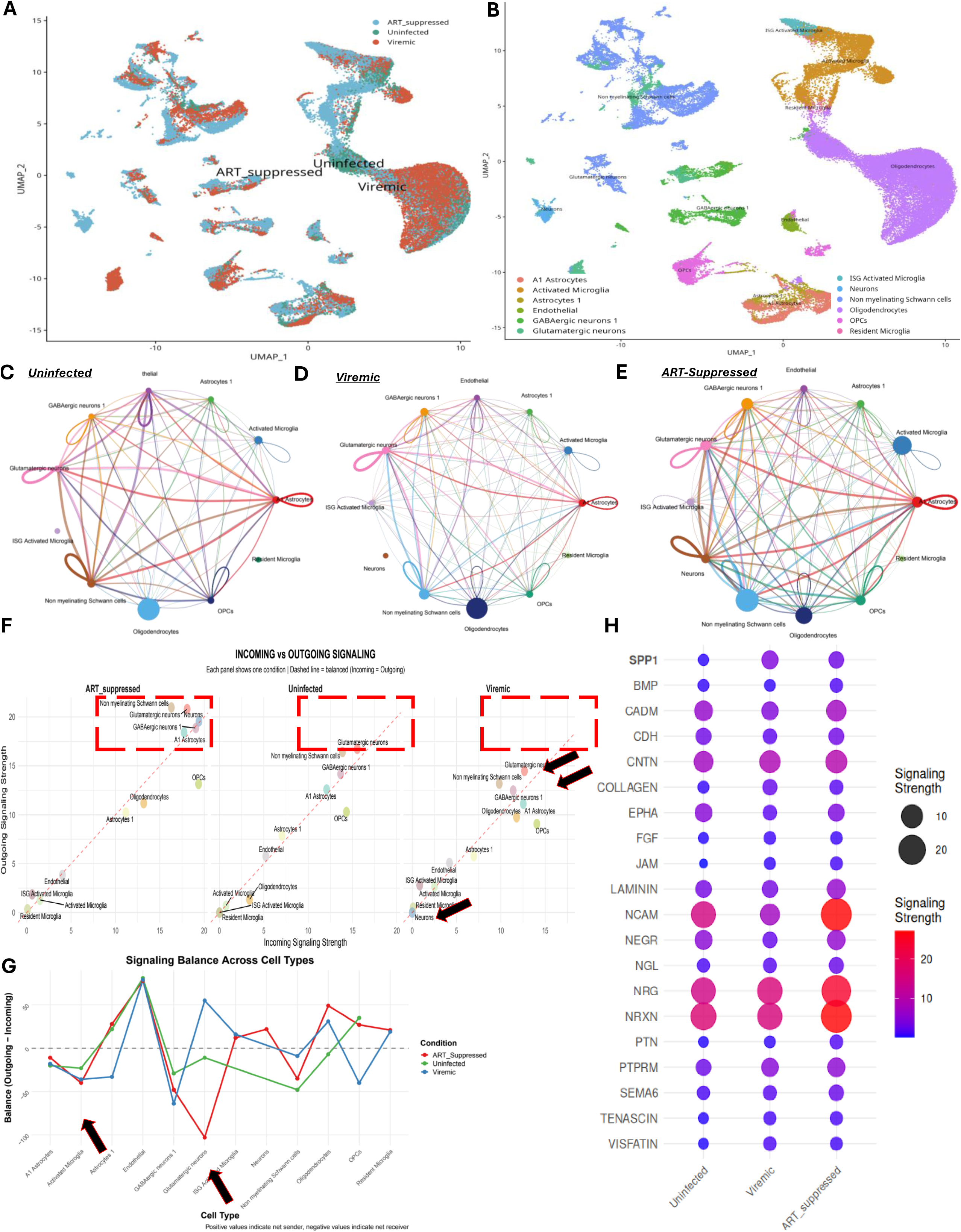
HIV reshapes conserved but modulated neuroimmune communications. **(A)** UMAP projection of integrated snRNA-seq datasets from uninfected, viremic, and ART-suppressed human brains showing donor distribution across infection states. **(B)** UMAP projection annotated by major brain cell populations, including oligodendrocytes, astrocytes, neurons, endothelial cells, oligodendrocyte precursor cells, and microglial subsets **(C–E)** Plots depicting condition-specific signaling networks for uninfected **(C**), viremic **(D)**, and ART-suppressed (**E**) brains. Edge thickness denotes communication strength, and node size reflects total outgoing signaling probability. **(F)** Scatter plots of outgoing versus incoming communication strength for each brain cell population across uninfected, viremic, and ART-suppressed conditions. Each point represents the mean signaling probability for a given cell type; dashed lines indicate equal outgoing and incoming strength. **(G)** Signal balance (outgoing-incoming) across cell types per condition. **(H)** Top 20 enriched predicted cell signaling pathways.

Microglial subset distribution analysis revealed striking differences across conditions: Activated Microglia expanded from 738 cells in Uninfected donors to 2,605 in Viremic (3.5-fold increase) and further to 4,330 in ART_Suppressed (5.8-fold increase). ISG-activated microglia showed even more dramatic expansion, increasing from 4 cells in Uninfected to 374 in Viremic (93-fold) and 539 in ART_Suppressed (135-fold). Resident microglia also expanded, though more modestly, from 7 to 65 (9-fold) to 82 (12-fold) (**Supplementary Fig. 3B**). Consistent with our previous report (*4*), these data demonstrate that microglial activation and expansion persist despite ART suppression, suggesting that viral exposure triggers long-lasting cellular reprogramming that is not reversed by therapy.

Chi-square analysis verified that while global immune cell compositions fluctuated across clinical conditions (χ² = 10,581, p < 2.2e-16, Cramér’s V = 0.29), the microglial subset internal architecture remained highly stable (χ² = 159.3, p = 2.1e-33, Cramér’s V = 0.095). The dissociation between global immune remodeling (Cramér’s V = 0.29, large effect) and microglial subset stability (Cramér’s V = 0.095, small effect) is critical for interpreting our findings. It indicates that the dramatic 5.8-fold expansion of activated microglia and 135-fold expansion of ISG-activated microglia are not simply passive consequences of global immune shifts (**Supplementary Fig. 3B**). Rather, they represent active, cell-intrinsic reprogramming driven by HIV exposure, a process that persists despite ART and fundamentally alters microglial function. This stable subset composition with dramatic functional changes suggests that microglia are not merely responding to the altered CNS environment but are themselves drivers of the pathological state.

Donor-resolved verification demonstrated the presence of this specialized SPP1⁺ microglial subset (average expression (lognorm) range 2.90-3.60, **Supplementary Fig. 3C, E**) with the activated MG across all five ART-suppressed donors, spanning a tight frequency range of 51.9%–98.8% (**Supplementary Fig. 3D, F**), while false-positive validation tracking confirmed zero HIV transcript reads within the activated microglial compartment of uninfected controls (**Supplementary Fig. 3G**).

We sampled 10,000 cells per condition and performed CellChat analysis (**Fig. 2C–E**) which revealed increasing information flow from uninfected (4,162 interactions) to viremic (4,574 interactions) to ART-suppressed (7,070 interactions) conditions (**Supplementary Table 2**). This represents a 70% increase in total communication events in ART-suppressed compared to uninfected, indicating that ART does not normalize intercellular communication but rather establishes a new, hyper-connected pathological state.

ISG-activated MG emerged as novel communicators under inflammatory conditions. The uninfected group showed no predicted interactions with ISG-activated MG, while such interactions were observed in both viremic and ART-suppressed conditions (**Fig. 2C-E**). ISG-activated MG showed 53 outgoing and 37 incoming interactions in viremic donors and 45 outgoing and 33 incoming interactions in ART-suppressed donors (**Supplementary Table 3**). This indicates that IFN-stimulated MG become communication-competent during viremia and remain active under ART suppression, representing a persistent pathological feature of the ART-suppressed CNS. Neuronal communication was disrupted during viremia but partially recovered on ART (**Fig. 2A– F**), suggesting that ART partially restores neuronal connectivity while failing to normalize microglial signaling.

Signaling balance revealed cell-type-specific communication roles that were fundamentally altered by HIV infection and ART. Activated MG consistently functioned as net receivers across all conditions (balance: Uninfected = -23, Viremic = -36, ART = - 40) (**Supplementary Table 3**), indicating that these cells are primarily recipients rather than senders of communication signals. In striking contrast, ISG-activated MG showed a net sender role in both viremic (balance = +16) and ART-suppressed (balance = +12), suggesting that IFN-stimulated MG shift from receiving to sending signals under inflammatory conditions. Resident MG also showed net sender roles in viremic (+19) and ART-suppressed (+21), though with fewer total interactions. Balance (outgoing minus incoming) signals did not return to baseline in glutamatergic neurons and activated MG, where the signals moved further from baseline in both ART-suppressed and viremic states (**Fig. 2F**), supporting persistent dysregulation of communication in these CNS cell types despite ART.

Pathway-specific communication revealed SPP1 signaling dominated during viremia (4.18), which persisted under ART (2.91), compared to uninfected (0.46) (**Fig. 2H; Supplementary Table 4**). Neuronal signaling pathways (NCAM, NRXN, NRG) showed the highest communication and were markedly enhanced under ART suppression compared to uninfected (NCAM – 1.85-fold increase, NRXN – 1.63-fold increase, NRG – 1.68-fold increase). In contrast, NCAM, CADM, EPHA, and NEGR were slightly suppressed in the viremic state, showing that active viral replication suppresses certain neuronal communication pathways while ART selectively enhances others. LAMININ signaling strength was highest in the ART-suppressed state (5.91 vs 3.42 in uninfected, 1.73-fold increase). Collectively, these results demonstrate that while inflammatory SPP1 signaling is profoundly disrupted during the viremic state and persists even under ART, neuronal signaling pathways are either conserved or enhanced under ART, with only a subset transiently suppressed during viremia. The persistent elevation of SPP1 signaling under ART coincides with ‘viral memory’ mechanism that is not resolved by viral suppression.

### Activated and ISG-Activated Microglia Exhibit Divergent Signaling Profiles

To characterize the functional differences between microglial subsets and their contribution to persistent CNS inflammation before segregating MG by SPP1, we performed differential expression and pathway enrichment analysis on Activated Microglia and ISG-activated microglia across conditions, as this would aid in distinguishing MG signatures and SPP1-driven signatures.

Activated Microglia showed a robust inflammatory signature in response to HIV. Comparing Viremic vs Uninfected, Activated Microglia upregulated 160 genes, including the iron transporter SLC11A1 (log2FC: 1.54), interferon-stimulated IFI44L (log2FC: 1.91), scavenger receptor MSR1 (log2FC: 1.76), hypoxia response HIF1A (log2FC: 1.46), and interferon-stimulated EPSTI1 (log2FC: 1.81) (**Supplementary Fig. 4A**). Concurrently, 514 genes were downregulated, including homeostatic markers P2RY12 (log2FC: -1.98) and CX3CR1 (log2FC: -2.28***), indicating loss of surveillance function. Pathway enrichment revealed activation of interferon response, inflammation, phagocytosis, and HIF-1 signaling, while synaptic signaling and neuronal development pathways were suppressed (**Supplementary Fig.4E-F**).

In ART Suppressed vs Uninfected, activated microglia showed a distinct metabolic signature with 50 upregulated genes, including SLC11A1 (log2FC: 1.98), cholesterol efflux transporter ABCA1 (log2FC: 1.57), MSR1 (log2FC: 1.47), HIF1A (log2FC: 1.34), and thymidine phosphorylase TYMP (log2FC: 1.43) (**Supplementary Fig. 4B**). However, 1,014 genes were downregulated, including stress chaperone HSP90AA1 (log2FC: - 3.90), myelin protein PLP1 (log2FC: -3.65), and heat shock protein HSPA1A (log2FC: - 2.25). KEGG pathway analysis revealed enrichment of Ferroptosis, Hepatitis C, and Measles pathways in Viremic Up, while Viremic Down showed synaptic signaling, cell adhesion, and MAPK/PI3K-Akt pathway suppression (**Supplementary Fig. 4G-H**). ART Suppressed showed persistent metabolic reprogramming with lipid/cholesterol metabolism and ABC transporter enrichment, but also widespread synaptic and signaling disruption (**Supplementary Fig. 4E-F**).

ISG-Activated Microglia exhibited a more restricted but highly specific transcriptional response. In ART_Suppressed vs Uninfected, ISG-MG showed only 1 upregulated gene MSR1 (log2FC: 4.00) but 17 downregulated genes, including centrosomal protein RTTN (log2FC: -4.00), synaptic plasticity gene SYNDIG1 (log2FC: -4.00**), stress chaperone HSP90AA1 (log2FC: -3.65), and DNA damage repair kinase ATM (log2FC: - 2.77) (**Supplementary Fig. 4C, 4D**). GO enrichment revealed regulation of lipid storage, cholesterol storage, and phagocytosis/engulfment (**Supplementary Fig. 4E**). KEGG pathway analysis showed Ferroptosis enrichment (**Supplementary Fig. 4F**).

MSR1 (Macrophage Scavenger Receptor 1) was the most dramatically upregulated gene across all conditions, with its highest expression observed in ISG-MG (log2FC: 4.00). SLC11A1 showed consistent upregulation across Viremic (1.54), ART (1.98), and ISG_ART (2.98) conditions (**Supplementary Fig. 4D**). MSR1 may serve as a key marker of persistent microglial activation and a potential therapeutic target for reducing neuroinflammation in ART-suppressed PWH. In contrast, homeostatic markers P2RY12 and CX3CR1 showed consistent downregulation, with the most profound suppression observed in ISG-MG (P2RY12: -3.91). ISG-MG-specific genes including RTTN (-4.00), SYNDIG1 (-4.00), ATM (-2.77), SORL1 (-3.41), and C3 (-3.25*) showed minimal expression changes in Activated MG but profound downregulation in ISG-MG (**Supplementary Fig. 4D**).

These divergent profiles-Activated Microglia showing broad inflammatory and metabolic reprogramming, while ISG-MG showing a focused stress response with profound synaptic gene suppression-distinguish how microglia subsets play distinct but complementary roles in CNS pathology. Activated Microglia drive persistent inflammation, while ISG-MG may contribute to synaptic dysfunction through loss of homeostatic support. The distinct transcriptional profiles of Activated and ISG-activated microglia and particularly the persistent upregulation of SPP1 signaling despite ART (**Fig. 2H**), prompted us to investigate whether SPP1^+^ cells represent a unique functional subset.

### SPP1-Driven MG-to-Neuron Signaling Persists Despite ART Suppression

Given that SPP1 was predominantly expressed by activated and ISG-activated MG subsets across both viremic and ART-suppressed states (**Fig. 3A**) and progressively increased from uninfected to viremic to ART (**Fig. 3B**), we hypothesized that this progressive elevation represents a form of ‘viral memory’, a persistent inflammatory signature that ART cannot erase.

**Figure 3.**
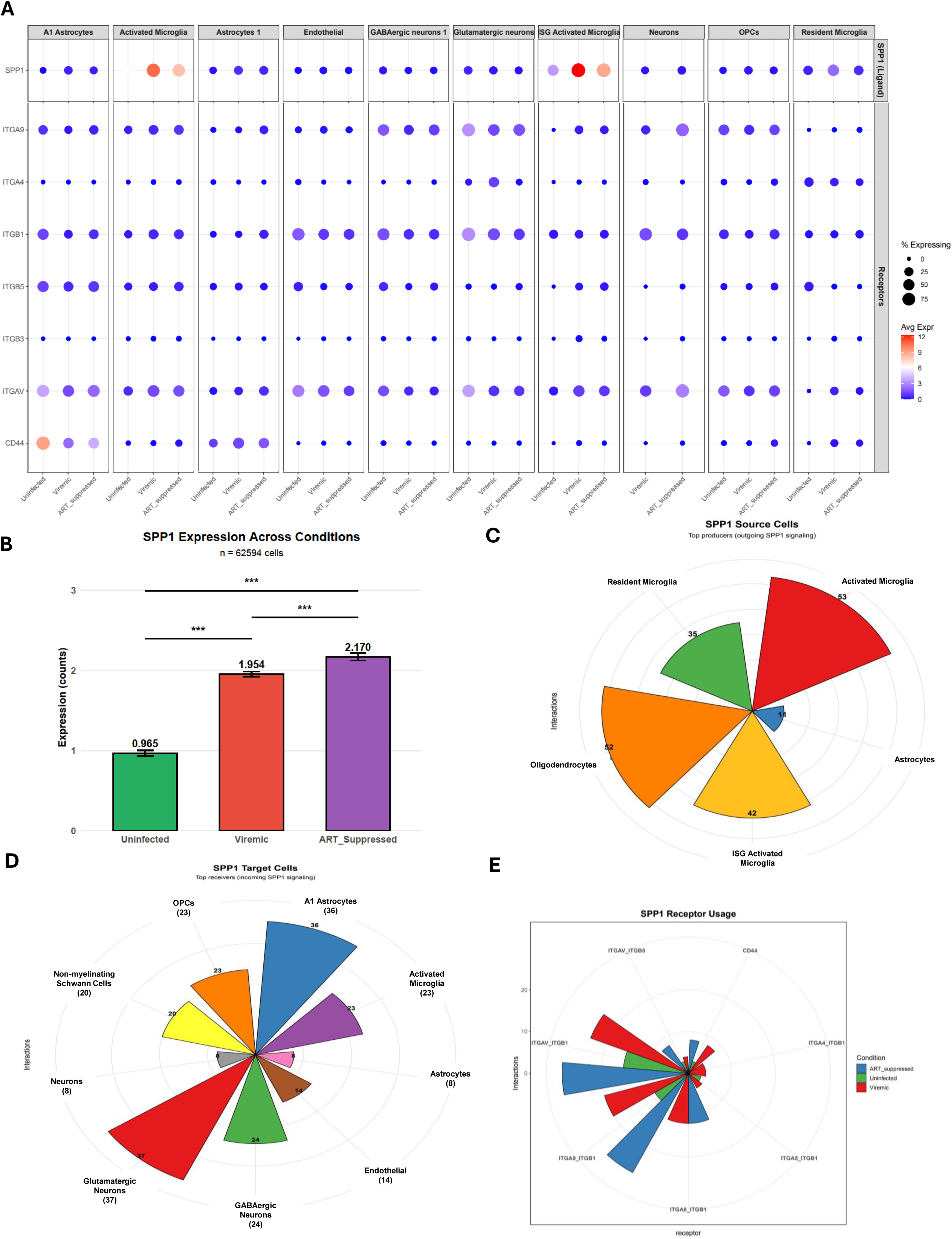
SPP1 Expression and Receptor Usage Across Cell Types and Conditions. **(A)** Dot plot showing SPP1 and its receptor expression (ITGA9, ITGA4, ITGB1, ITGB5, ITGB3, ITGAV, CD44) across cell types (A1 astrocytes, activated MG, astrocytes 1, endothelial, GABAergic neurons, glutamatergic neurons, ISG-activated MG, neurons, OPCs, and resident MG) in ART-suppressed donors. Dot size represents the percentage of cells expressing each receptor, and color intensity represents average expression level. **(B)** Bar graph showing SPP1 expression across conditions (uninfected: 0.965, viremic: 1.954, ART-suppressed: 2.170), demonstrating progressive increase in SPP1 expression from uninfected to viremic to ART-suppressed states. **(C)** Top source cells sending SPP1 signals, including activated microglia and oligodendrocytes, based on number of interactions. **(D)** Top target cells receiving SPP1 signals, including OPCs and A1 astrocytes, based on number of interactions. **(E)** Pie chart illustrating SPP1 receptor usage, highlighting ITGAV_ITGB5 and CD44 as dominant SPP1 receptors.

MG subsets were the top senders of SPP1 signals (**Supplementary Table 5**). In ART-suppressed brains, SPP1 source cell type predicted interactions (PI) included activated MG (PI=53), ISG-activated MG (PI=42), resident MG (PI=35), oligodendrocytes (PI=52), and astrocytes (PI=11) (**Fig. 3C**). Activated MG are the dominant source of SPP1 signaling in the ART-suppressed brain, with ISG-activated MG also contributing substantially. The primary receivers of SPP1 signals were glutamatergic neurons (PI=37) and A1 astrocytes (PI=36) (**Fig. 3D**), demonstrating an SPP1-mediated communication axis from microglia to neurons and astrocytes in the ART-suppressed CNS.

Distinct SPP1 receptor usage across conditions provided further insight into the nature of this signaling. Integrin receptors were dominant in ART-suppressed and viremic conditions (**Fig. 3E**). Specifically, ITGA4_ITGB1 was present only in viremic donors, while ITGA5_ITGB1 usage was higher in viremic compared to uninfected and was not observed in ART-suppressed donors (**Fig. 3E**). This receptor switching suggests that the functional consequences of SPP1 signaling differ between viremic and ART conditions, potentially contributing to distinct pathological outcomes. SPP1 microglia target A1 astrocytes via CD44 in A1 astrocytes (**Fig. 3A, D, and E**), supporting that SPP1-expressing microglia induce neuropathogenesis via CD44 in neurotoxic astrocytes in viremic and ART-suppressed brain.

The progressive increase in SPP1-expression from uninfected to viremic to ART-suppressed, coupled with distinct receptor usage patterns under ART versus viremia, indicates that ART does not normalize SPP1-driven communication. Instead, a persistent and distinct SPP1^+^ microglia signaling network remains, potentially contributing to ongoing neuroinflammation and incomplete neuronal recovery despite viral suppression.

### SPP1^+^ Activated MG Subset Marks a Persistent Reservoir and Drives Neuronal Communication

To determine whether SPP1^+^ microglia represent a distinct functional subset that could explain this persistent signaling, we stratified activated MG by SPP1 expression across all three conditions. SPP1 was upregulated in activated MG and ISG-activated MG in both viremic and ART-suppressed donors, with a similar trend observed in ART-suppressed donors (**Fig. 4A**). This was further validated in brain tissue in vivo by CD68^+^/SPP1^+^ immunohistochemistry (**Fig. 4B**), where SPP1 expression during ART was >2-fold, compared with uninfected controls.

**Figure 4.**
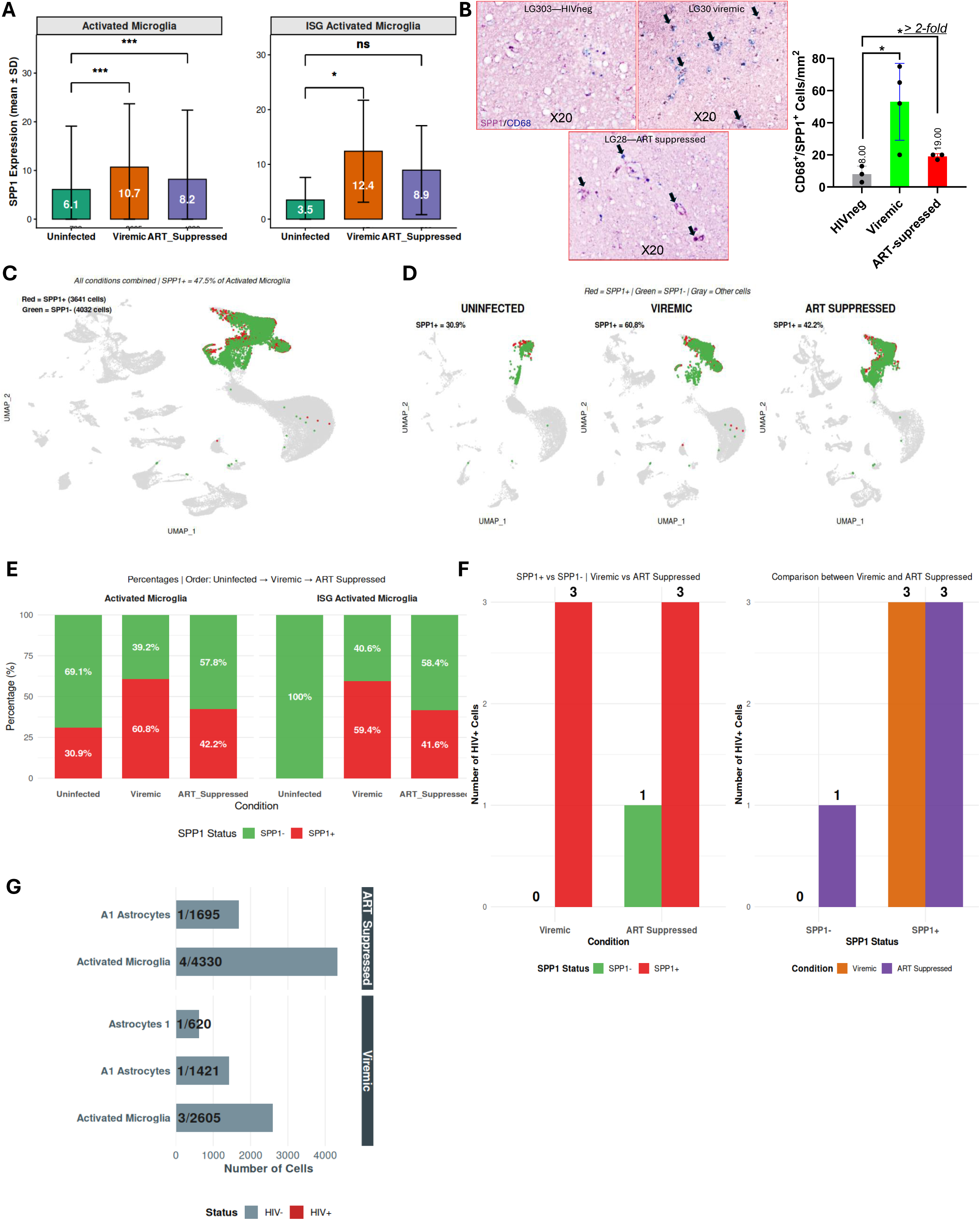
SPP1⁺ MG persist across infection states and drive immune activation and adaptive remodeling in ART-suppressed brains. **(A)** Expression of SPP1 across conditions in activated MG and ISG-activated MG subsets, demonstrating elevation in viremic and ART-suppressed donors. **(B)** Immunohistochemical analysis of the frontal cortex tissues shows induced expression of SPP1 protein CD68^+^ myeloid cells in viremic donors, which persists (> 2-fold) on ART. Images are displayed at 20× magnification. Purple: SPP1; Blue: CD68. **(C)** UMAP projection of all brain cells highlighting SPP1⁺ MG (red) vs SPP1-MG (green). **(D)** UMAP projections across uninfected, viremic, and ART-suppressed donors showing progressive expansion of SPP1⁺ MG. **(E)** SPP1⁺ microglial abundance by condition, demonstrating marked enrichment in ART-suppressed tissue. **(F)** Fraction of HIV⁺ cells among SPP1⁺ and SPP1⁻ MG subsets, showing the highest shift in viremic and ART-suppressed activated MG. HIV+ RNA frequency is markedly higher in SPP1⁺ activated MG (6/3299) compared with SPP1⁻ (1/3178). Comparative fraction of HIV⁺ cells across conditions demonstrates that SPP1⁺ MG harbor the majority of detectable HIV⁺ nuclei under both viremic and ART-suppressed states, confirming preferential viral persistence within the SPP1⁺ subset. **(G)** Relative abundance of HIV⁺ cells within each annotated cell population under viremic and ART-suppressed conditions. The gray bars represent total cell counts per cell type. HIV^+^ cell/ total cells detected per population (n). HIV⁺ cells were primarily enriched within activated MG under both conditions. Minimal infection was detected in astrocytes, further supporting MG as the dominant HIV reservoir in the CNS.

Activated MG (excluding ISG-activated MG) were stratified by SPP1 expression (SPP1^+^ vs SPP1^-^ activated MG, **Fig. 4C**) per condition (**Fig. 4D**). SPP1^+^ MG were defined as cells with detectable SPP1 expression (normalized expression > 0), while SPP1-neg MG lacked SPP1 expression (normalized expression = 0). Both viremic and ART-suppressed donors exhibited a clear shift in the ratio of SPP1^-^ to SPP1^+^ MG (**Fig. 4E**) that did not return to the uninfected ratio even under ART. Interestingly, ISG-activated MG showed SPP1^+^ MG expansion during viremia and ART suppression (**Fig. 4E**), further supporting homeostatic dysregulation (**Supplementary Fig.4**), where SPP1^+^ MG become the dominant phenotype under inflammatory conditions and persist despite viral suppression.

We further examined whether SPP1^+^ MG harbor residual HIV and whether this population contributes to the ongoing neuroinflammatory state observed in ART-suppressed PWH. Critically, SPP1^+^ MG remained dominant during ART and contained the highest number of HIV RNA^+^ cells (6 of 7) (**Fig. 4F**): 3 HIV^+^ cells in viremic SPP1^+^ activated MG, 3 HIV^+^ cells in ART SPP1^+^ activated MG, and 1 HIV^+^ cell in ART SPP1-activated MG. HIV^+^ cells were in activated MG and persisted in ART (**Fig. 4G**), supporting that SPP1^+^ activated MG is a key MG reservoir of HIV persistence (*4*). This finding directly links the persistent SPP1 signaling we observed to the cellular reservoir, establishing SPP1^+^ MG as both the source of inflammatory signals and the MG subset of viral persistence.

### HIV RNA^+^/SPP1^+^ Activated MG Subset Are Reprogrammed Toward Immune-Evasive, Pro-inflammatory State with Synapse Destabilization in ART-suppressed individuals

Having determined that SPP1^+^ MG tend to harbor residual HIV RNA, we profiled the transcriptomes of HIV RNA^+^ cells. In donor 1, HIV transcripts were detected in 1 of 1,324 MG (∼0.1%), localized in the activated MG cluster. Donor 2 showed a similar frequency in the activated MG cluster (2 of 3,332 MG; ∼0.1%). These values align with reports of <0.1% reservoir frequency (*18*), confirming that rare HIV RNA^+^ MG persist in the brain despite ART. We then compared HIV^+^ MG with nearby HIV^-^ MG from the same tissue using K-Nearest Neighbors (KNN) clustering (*19*) (***Supplementary* Fig. 5A-C**) to identify cell-intrinsic transcriptional changes driven by residual viral transcription. Across three independent groups, Group 1: HIV^+^ MG expressed tolerance/stress regulators (e.g., GDF15, CD300LB), whereas HIV^-^ MG upregulated immune surveillance genes (ETS2, ATP6V1B2) (***Supplementary* Fig. 5D**). Group 2: HIV^+^ MG upregulated immune-evasive transcripts (PRDM1, TNFAIP3), while HIV^-^ MG were enriched for antiviral mediators (CTSB, GRB2) (***Supplementary* Fig. 5E**). Group 3: HIV^+^ MG expressed innate activation and chemokine genes (CD14, TLR2), contrasting with HIV^-^ MG enrichment for antigen presentation (HLA-DRB1, SQSTM1) (***Supplementary* Fig. 5F**).

Differential expression (adj. p < 0.05, logFC ≥ 0.5) identified 533, 144, and 169 DEGs in HIV^+^ MG versus 83, 164, and 65 DEGs in HIV^-^ MG, with minimal overlap (2–4 genes) across groups (***Supplementary* Fig. 5G-I**). This divergence demonstrates that, although rare, HIV^+^ MG are individually transcriptionally distinct while showing an immune-evasive phenotype. Thus, combining all groups, we identified 824 HIV^+^-specific and 303 HIV^-^-specific genes, with filtering (logFC ≥ 0.5) yielding 295 high-confidence HIV^+^ genes and 10 HIV^-^ genes. HIV^+^ MG were reprogrammed toward immune-driven inflammation (**Supplementary Table 6**), characterized by suppressed apoptosis and enrichment of cancer-associated transcriptional networks (***Supplementary* Fig. 5J**). Consistent with the SPP1^+^ MG signature (***Supplementary* Fig. 2D**), apoptotic programs were inhibited. HIV^-^ MG, in contrast, were enriched for apoptosis, viral suppression, and CD44-mediated adhesion modules, consistent with a turnover-competent, homeostatic phenotype (***Supplementary* Figs. 5J and 2D**).

Innate immune cascades were enriched in HIV^+^ MG, including MyD88-dependent endosomal signaling, TLR4 activation, iNOS, and SLC15A4: TASL–IRF5 pathways, all converging on type I interferon (IFN-I) responses (*20*) and neuronal signaling via glutamate receptor genes (DLG4, GNB5, HOMER3) (***Supplementary Figs. 5K-L***), highlighting how HIV^+^ cells exert pressure on glutamatergic neurons. The upregulation of DLG4 and GNB5 in MG supports phagocytosis of synapses, while increased HOMER3 supports active interaction with synapses. Together, these findings demonstrate how HIV may cause MG-mediated synapse injury that is partially reversed by ART. Interestingly, MHC-I and metabolic pathways were highly activated in HIV^+^ MG (***Supplementary Fig. 5M***). HIV^+^ MG showed enrichment of CoA biosynthesis and amyloid precursor protein (APP) pathways (***Supplementary Fig. 5N***), consistent with previous reports of β-amyloid accumulation in HAND brains (*21*)(*22*). COASY and PPCDC (CoA pathway), as well as NCSTN and AGO2 (APP pathway), were upregulated (***Supplementary Fig. 6A***), linking HIV persistence to amyloidogenic processing and neurodegeneration (*23*). The stem-cell-associated regulator BCL7B (***Supplementary Fig. 6A***) was upregulated (***Supplementary Fig. 6A***), supporting long-term survival of the HIV+ MG reservoir. Multifunctional regulators included CHUK, IKBKG, and LY96 (***Supplementary Fig. 6B***), encoding IKK and TLR/MyD88 components contributing to NF-κB–driven inflammation (*24*). This transcriptional divergence reveals a striking dichotomy, where HIV^+^ cells abandon homeostatic functions and adopt immune-evasive, pro-inflammatory programs while HIV^-^ cells retain homeostasis.

HIV^+^ MG expressed pro-inflammatory ligand, including CCL3, CCL4, and CXCL8 (***Supplementary Fig. 6C-E***). HIV^+^ MG predicted ligands showed strong interactions with IFNGR1 (**Supplementary Fig. 6E**), confirming HIV^+^ cells send IFN signals to neighboring HIV^-^ cells. HIV^-^ MG were enriched in lipid degradation, autophagy, and phagosome formation (**Supplementary Fig. 6F**), with CTSH and DHRS7 supporting redox balance (**Supplementary Fig. 6G-H**). IFNGR1 was suppressed in HIV^+^ MG but elevated in HIV^-^ MG, illustrating how HIV^+^ MG function as senders rather than receivers of IFN signaling (**Supplementary Fig. 6I-J**). Thus, HIV^+^ MG function primarily as broadcasting hubs while insulating themselves from antiviral feedback.

Together, these findings demonstrate how HIV⁺ MG abandon homeostatic degradation but instead adopt anabolic and stress-adapted programs. This reprogramming links the persistence of SPP1⁺ MG HIV reservoirs to neuroinflammation, synapse destabilization, and neurodegeneration in ART-suppressed individuals (*12*).

### HIV^+^ MG Show Persistent IFN-I/MHC-I Immune-Evasive Signature in ART-Suppressed PWH

Both scRNA-seq (***Supplementary Fig. 5***) and snRNA-seq (**Fig. 5**) confirmed the localization of HIV RNA⁺ in the activated MG subset during ART suppression. We next expanded and focused on the HIV⁺ activated MG subset to confirm the reproducibility and biological consistency of this phenotype. Integration of snRNA-seq datasets from HIV⁺ and HIV⁻ MG within ART-suppressed donors (p = 3.83 × 10⁻¹¹; **Fig. 5A & B and *Supplementary Table 1***) using KNN clustering identified 410 HIV⁻-specific, 232 HIV⁺-specific, and 162 shared genes (**Fig. 5C**). HIV⁺ MG were enriched for MHC-I and IFN pathways, both of which are regulated by IFN-I signaling (*26, 27*) (**Fig. 5C-D**). Gene set analysis further revealed activation of neuroinflammatory and neurodegenerative modules, such as multiple sclerosis and pathogen-induced cytokine storm signaling (**Fig. 5D–E).** These findings confirm that HIV⁺ activated MG sustain an IFN-I-dependent neuroinflammatory program under ART. By contrast, HIV⁻ MG (**Fig. 5F–G**) showed growth-factor and receptor-mediated pathways, including serotonin, estrogen, and insulin cascades, reflecting a stable, reparative MG state. Thus, IFN-I/MHC-I-mediated neuroinflammation may be central to the neurotoxic landscape of HIV⁺ MG, consistent with scRNA-seq contrasts between HIV⁺ and HIV⁻ MG (***Supplementary Fig. 5***) and our previous observations in the brains of PWH with HAND (*12*).

**Figure 5.**
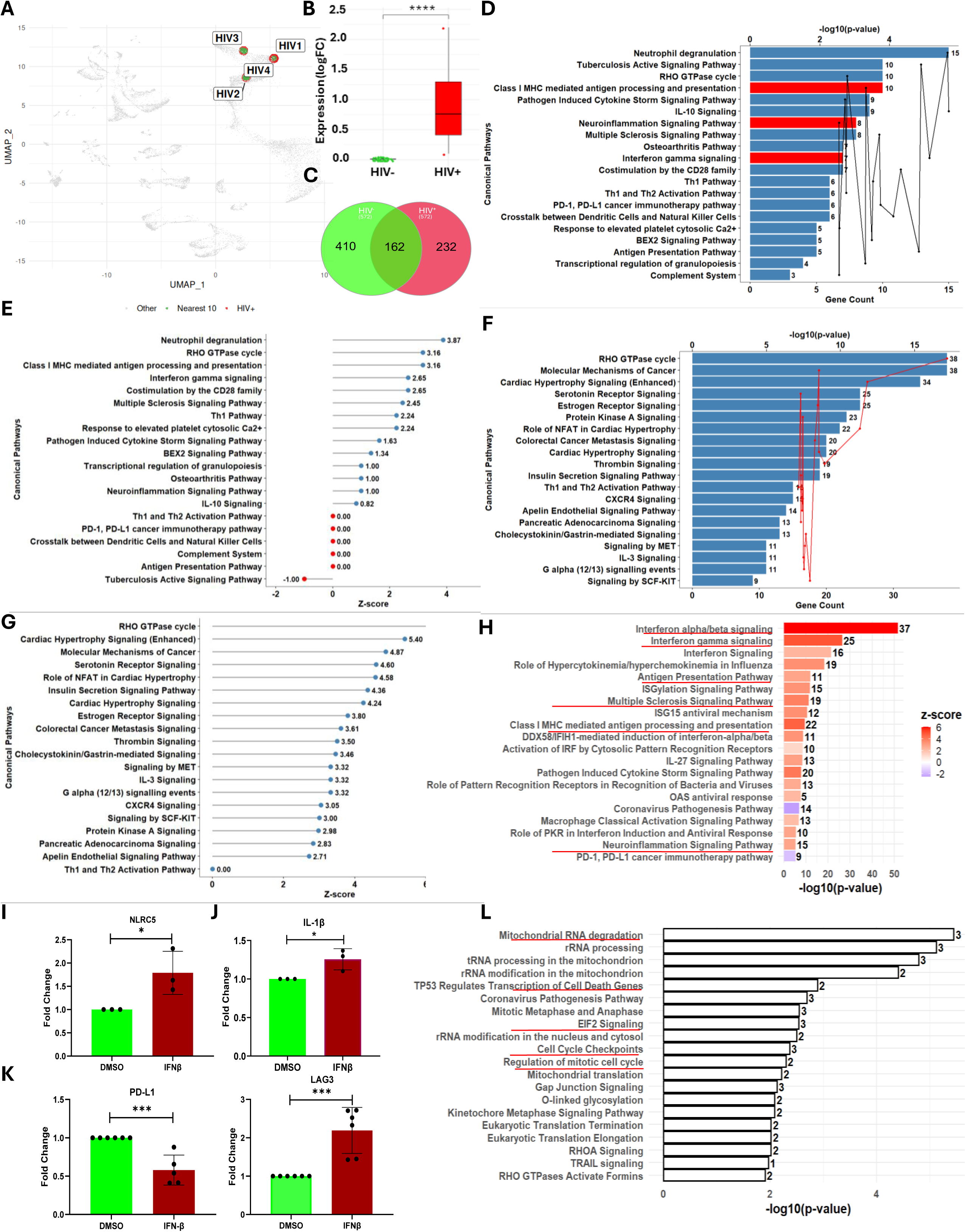
HIV⁺ MG exhibits distinct transcriptomic signatures marked by inflammation activation and MHC-I signaling activation. **(A)** UMAP projection of single-nucleus transcriptomes showing localization of HIV⁺ nuclei (red) within microglial clusters. Inserts highlight individual HIV⁺ cells identified by viral transcript expression with neighboring cells without HIV RNA (HIV^-^). **(B)** Normalized HIV transcript expression (log₂ FC) between HIV⁻ and HIV⁺ nuclei, showing significantly higher viral RNA levels in HIV⁺ cells (**** p < 0.0001). **(C)** A Venn diagram illustrates the overlap of DEGs between HIV⁻ (572 genes) and HIV⁺ (394 genes) nuclei, with 162 shared DEGs representing core transcriptional alterations. **(D, F)** Canonical pathway enrichment of HIV⁺ (D) and HIV^-^(F) DEGs. **(E, G)** Z-scores depicting pathway activation directionality in HIV⁺ (E) and HIV^-^(G) DEGs; positive values denote activation, while negative values denote suppression. Gene count and p-value ranked the most significant canonical pathways. **(H)** Canonical pathway enrichment of upregulated genes showing strong activation of IFN-I signaling, antigen presentation, TLR/MyD88 cascades, and immune synapse pathways. Bar colors denote activation z-scores (red = activated, blue = suppressed). **(I-K)** Expression of inflammasome (NLRC5, IL1B) and immune checkpoint genes (PD-L1, LAG3) after IFN stimulation in the brain MG. **(L)** Enrichment of downregulated genes highlighting suppression of mitochondrial RNA processing, translation, and metabolic pathways, consistent with disrupted oxidative and biosynthetic function. Red lines: Highlighted signaling discussed in the text.

To further determine whether the IFN-I/MHC-I-linked immune-evasive phenotype observed in ART-suppressed HIV⁺ MG also occurs during active viremia, we integrated transcriptomes from HIV⁺ and HIV⁻ MG populations from viremic donors (***Supplementary Fig. 7A***). HIV expression and transcriptome were confirmed (p = 2.32 × 10⁻⁸; ***Supplementary Fig. 7B-C***). In viremic donors, HIV⁺ MG (***Supplementary Fig. 6D***) and HIV⁻ MG (***Supplementary Fig. 7E***) enriched pathways (***Supplementary Fig.7D***) that lacked the defining transcriptional hallmarks of the ART-suppressed state, such as the robust IFN-I/MHC-I amplification.

Together, these findings reveal that long-term ART reshapes MG immunity from an antiviral state during active viremia into a chronic, IFN-I-driven, MHC-I-amplified, immune-exhausted phenotype.

### IFN Signaling Directly Amplifies MHC-I Activation and Neuroinflammation While Suppressing PD-L1 Immune Checkpoint in MG

The strong signal sent from SPP1 in HIV⁺ MG to IFNGR1 in HIV⁻ MG (***Supplementary Fig. 6E***) in scRNA-seq and interferon-associated (MHC-I and IFN) signaling (**Fig. 5D**) in snRNA-seq suggested an IFN-driven feedback loop in these cells, as we recently reported (*12, 26*). To test whether IFN directly drives the neuroinflammatory transcriptional landscape, purified MG from people without HIV were treated with IFNβ for five days to mimic chronic exposure to IFN-I and analyzed by bulk RNA-seq.

IFN-I exposure induced 377 DEGs, including 329 upregulated and 48 downregulated (***Supplementary Fig. 8A-B and Supplementary Table 7***), demonstrating robust activation of canonical IFN-I pathways. Upregulated transcripts reflected broad immune activation, dominated by interferon-stimulated genes (ISGs) and MHC-I antigen processing modules, including RIG-I, MDA5, IRFs, and TLR3 (**Fig. 5H**). Inflammasome signaling was also activated (***Supplementary Table 7***), with increased NLRP3 and IL-1β expression confirmed by RT-qPCR (**Fig. 5I-J**). Cytokine, chemokine networks and neurological disease pathways were amplified, showing converging pressure on the CNS (**Fig. 5H**). PD-L1 checkpoint signaling was suppressed (z-score = –1.63), consistent with the inhibition of PD-L1 expression by IFN-I, indicating immune exhaustion in MG (**Fig. 5H and K**). Of note, the immune checkpoint LAG3 gene was upregulated (**Fig. 5K**), a known immune regulator in MG that plays a role in neurodegeneration (*28*).

Downregulated pathways primarily involved mitochondrial function and protein synthesis, including EIF2 signaling, ribosomal activity, and translational elongation (**Fig. 5L**), indicating bioenergetic stress. The concurrent regulation of TRAIL and p53-dependent cell death pathways (*29, 30*), reflects a balance between stress-induced apoptosis and survival.

Collectively, these findings suggest that IFN-I signaling in MG promotes immune and inflammasome activation, and mitochondrial exhaustion. This state mirrors the transcriptional landscape of HIV⁺ MG in vivo (sc and snRNA-seq), where viral RNA induces IFN signaling sustaining inflammation while exhausting cellular energy reserves.

### IFN-I Signaling Broadly Activates MHC-I within SPP1^+^/HIV^+^ MG

From our findings, persistent IFN-I signaling in HIV⁺ MG led to widespread induction of MHC-I (***Supplementary Fig. 5M* and Fig. 5D**). Classical HLA-A (logFC = 1.68), HLA-B (2.10), HLA-C (1.79), and non-classical HLA-E (1.17), HLA-F (2.22), and HLA-G (logFC = 1.8) were all markedly upregulated, with B2M (1.29) for complex stabilization and TAP1 (1.04) for peptide translocation. Antigen-processing components, including PSMB9 (1.56) and CTSS (2.35), were similarly elevated, reinforcing the expansion of IFN-driven antigen presentation.

Post-translational regulators were elevated, including ISGylation enzymes (HERC5, HERC6, UBA7, UBE2L6) and E3 ligases (FBXO6, TRIM21, TRIM69) (**Supplementary Table 7**), suggesting tighter control of antigen loading and turnover. MHC-I was activated in ART-suppressed HIV⁺ MG, which was regulated through perturbation of the ubiquitin–proteasome system (**Supplementary Fig. 9A**). Key regulators, including FBXW5, SIAH2, and ANAPC10, were upregulated in MG (**Supplementary Fig. 9B**). Regulators critical for proteasomal degradation and the APC/C complex, which control antigen degradation (**Supplementary Fig. 9A**; p = 0.00194) (*31*). IKBKG was associated with LY96 and TLR4 regulation, suggesting a feed-forward NF-κB-dependent inflammatory loop (*32*) that may contribute to ER stress (*33*). Importantly, SPP1 signaling was associated with IKBKG regulation (p = 0.00182) during MHC-I activation (**Supplementary Fig. 9C**), suggesting a mechanistic link between SPP1-driven inflammation and MHC-I dysfunction via the TLR–NF-κB axis (*34*).

Expanding on MHC-I signaling using snRNA-seq, network analysis identified a tightly coordinated MHC-I module in HIV⁺ MG (p = 4.61 × 10⁻⁵; **Supplementary Fig. 9D**). This cluster encompassed genes regulating antigen degradation (BTBD1, CD14), peptide transport (TAP2), ER trimming (ERAP1, RCHY1), MHC-I loading (HLA-E, TRIM36), vesicular trafficking (SNAP23, MRC2), and surface presentation (FCGR1A). Collectively, the results support convergence of IFN and MHC-I pathways in HIV⁺ MG.

Expression validation showed upregulation of TAP2, SNAP23, RCHY1, MRC2, HLA-E, FCGR1A, and ERAP1 in HIV⁺ MG (**Fig. 6A**). In vitro, chronic IFNβ treatment of MG recapitulated this pattern, inducing TAP2 and SEC23A (**Fig. 6B**). The presence of a canonical IFN-I-stimulated response element in the TAP2 promoter (*35*) suggests its direct regulation by IFN-I.

**Figure 6.**
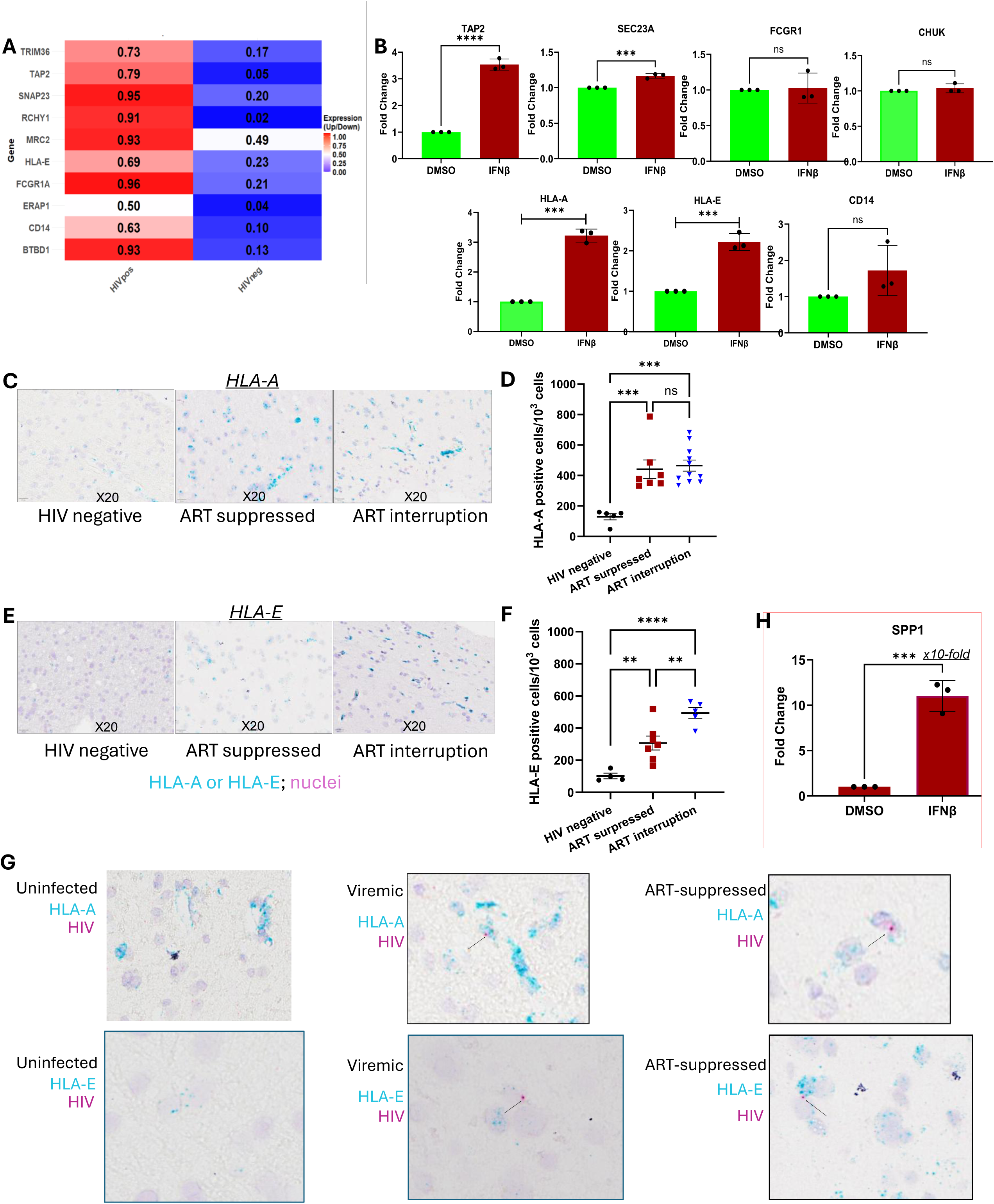
SPP1⁺ MG are Associated with Persistent IFN/MHC-I Activation in the CNS Despite ART Suppression. **(A)** Expression of MHC-I-associated genes (TAP2, SEC23A, FCGR1A, CHUK, HLA-A, HLA-E, CD14) in HIV⁺ versus HIV⁻ MG from ART-suppressed donors. **(B)** qPCR validation showing expression levels of MHC-I-associated genes (TAP2, SEC23A, FCGR1A, CHUK, HLA-A, HLA-E, CD14) in IFN-stimulated MG. **(C-D)** Immunohistochemical detection and quantification of HLA-A in uninfected, ART-interrupted, and ART-suppressed frontal cortex brain tissues, showing high HLA-A expression in viremic donors that persists under ART suppression. **(E-F)** Immunohistochemical detection and quantification of HLA-E in uninfected, ART-interrupted, and ART-suppressed frontal cortex brain tissues, demonstrating persistent HLA-E expression despite ART. **(G)** HIV and HLA-A or HLA-E co-staining within the same frontal cortex brain tissues (20X), indicating that HIV-infected cells express elevated MHC-I molecules under ART suppression. **(H)** qPCR validation showing expression of SPP1 in IFN-stimulated MG.

Interestingly, SEC23A induction suggests a potential novel ISG role that warrants further investigation. Consistent with these transcriptional profiles, brain tissues from ART-suppressed donors displayed increased MHC-I gene expression (HLA-A and HLA-E) compared with HIV^-^ controls (***Supplementary Fig. 10A-B***). HIV⁺ MG confirmed this pattern, with elevated HLA-A and HLA-E in ART-suppressed donors (***Supplementary Fig. 10C***) and no expression in HIV⁺ MG from viremic donors. Importantly, direct IFN-I stimulation of MG upregulated HLA-A and HLA-E (**Fig. 6B**), highlighting the role of IFN signaling in MHC-I activation.

Notably, IFNβ did not induce FCGR1A, CHUK, CD14, or SNAP23 in MG (**Fig. 6B**), suggesting that HIV may not directly stimulate their expression. Despite that, MHC-I and inflammatory signaling activation (***Supplementary Fig. 10D***, p = 2.46e-11) were associated with pressure on neuronal genes (SNAP23) (**Fig. 6A**). The findings were corroborated by IHC analysis, showing significantly elevated HLA-A and HLA-E expression in brain tissues from ART-suppressed PWH (**Fig. 6C-F**), with some HLA-A and E co-localizing with or nearby HIV⁺ cells in the brain (**Fig. 6G).** Furthermore, SPP1 was directly stimulated 10-fold by IFNβ in brain MG, compared with untreated controls (**Fig. 6H**). Together, these observations suggest that the upregulated MHC-I signaling is IFN-dependent and attributed to SPP1-mediated inflammatory stimuli in activated MG on ART.

### Viral Sequence Recovery Identifies Nef-Mediated Disruption of MHC-I during Antigen Presentation in Brains from PWH on ART

Viral sequence recovery in MG from PWH on ART revealed HIV transcription of the 3’ LTR and HIV antisense transcript ASP among donors 1-5 on ART (*36*), with dominant expression from the 3’ Nef ORF tail (***Supplementary Fig. 11A***) and silencing of other genes. This pattern differs from HIV transcripts recovered in MG from ART-interrupted PWH, where *gag* and *integrase/vif* were also detectable in addition to *nef* RNA. Consistent with persistent *Nef* transcripts and protein expression in PWH with HAND (*37–40*), *Nef* protein was detected by IHC in both ART-suppressed and ART-interrupted brains, but no in uninfected donors (***Supplementary Fig. 11*B-C**). Since *nef* expression requires less sustained transcriptional elongation, driven by basal LTR activity for short elongation bursts, and is less dependent on *Tat* than structural genes like *gag* or *pol*, these data strongly support a latent HIV infection in brain MG during ART.

HIV *Nef* is known to suppress MHC-I antigen processing and presentation through *Nef*-mediated MHC-I endocytosis (*41*). Furthermore, *Nef* directly binds the HLA-A cytoplasmic tail to disrupt MHC-I trafficking (*42*) and downregulates HLA-E surface expression (*43*). Structural modeling predicted a stable *Nef*–HLA-A interaction, with *Nef* binding HLA-A via residues Glu11–Ala34, Cys14–Cys31, and Thr33–Glu11, forming atomic contacts at 2.6–3.9 Å (***Supplementary Fig. 11*D**). In addition, parallel interactions with HLA-E were detected, including Met45–Cys31 (3.63 Å), Gln43–Thr33 (3.84 Å), and Ala29–Val34 (3.50 Å) (***Supplementary Fig. 11*E**). These conserved molecular contacts provide structural modeling evidence that Nef may not only impact MHC-I expression but also directly disrupt antigen presentation, contributing to HIV persistence in the CNS of PWH on ART. These findings suggest that persistent expression of HIV Nef may impair effective MHC-I antigen presentation in the CNS.

### Inflamed SPP1^+^ MG and TREM2–C1QC/ “Eat Me” Signaling on ART

Next, we investigated whether HIV⁺ activated MG were associated with neuronal injury under ART. Both HIV⁺ and HIV⁻ MG interacted with synapses and glutamatergic neurons (***Supplementary Fig. 12A-B***). HIV⁺ MG appeared to adopt pro-active and remodeling functions related to neuronal injury, differentiation, migration, and transport, targeting the post-synapse. In contrast, HIV⁻ MG exhibited a profile focused on homeostasis at mature synaptic sites (post-synapse, axon) and immune-neuronal communication (immunological synapse) (***Supplementary Fig. 12C***), supporting a balanced regulatory role.

Furthermore, HIV⁺ MG were enriched for synaptic vesicle fusion and synapse pruning pathways (**Supplementary Fig. 13A**), marked by enrichment of TREM2 and *C*1QC, the core components of the “Eat-Me” phagocytic signaling (*45*). *SNAP23* and STX11, critical components of the SNARE complex, were also highly induced following HIV infection and persisted during ART. These genes were enriched in HIV⁺ MG, implicating excessive presynaptic elimination (**Supplementary Fig. 13A**). Top DEGs (IL1B, C1QC, TREM2, APOE) confirmed that SPP1⁺ MG are associated with neuroinflammation and synaptic pruning (**Supplementary Fig. 13B**), processes critical for cognitive decline (*46, 47*).

Of note, HLA-A, HLA-E, TAP2, CTSH, and synapse-pruning genes (C1QC, TREM2) were upregulated in SPP1⁺ MG from PWH on ART compared with SPP1⁻ MG. IFN-I regulators (IRF5), APP-associated genes (NCSTN, AGO2, APP), and coenzyme A metabolism genes (PPCDC, COASY) were upregulated in SPP1⁺ MG, along with DAM marker ITGAM and the immune checkpoint gene CD274 (***Supplementary Fig. 13C-I***). Furthermore, IL-1β was upregulated and remained high across conditions (viremic and ART-suppressed) (***Supplementary Fig. 13J***), supporting persistent inflammation in SPP1⁺ MG despite ART. Top pathways in this subset included cell adhesion, DAM activation, inflammation, and phagocytosis (***Supplementary Fig. 14A***), dominated by SPP1 and TREM2 signaling, reflecting a positive feedback loop linking HIV infection, SPP1 expression, and IFN activation.

To assess complement activation,the expression of classical complement molecules was analyzed (***Supplementary Fig.14B***). SPP1⁺ MG exhibited higher complement activation across all conditions compared to SPP1⁻ MG, with especially in ART-suppressed donors. C3 was elevated in SPP1⁺ MG (5.22) compared to SPP1⁻ MG (1.62) (***Supplementary Fig. 14B***). Complement regulators CD59 and CFH remained at baseline levels, suggesting that complement activation is not counterbalanced. Correlation analysis of C3, SKAP1, PTPRM, IL1B, and THEMIS revealed positive correlations between C3 and PTPRM (0.16) and between C3 and IL1B (0.11), and negative correlations between C3 and SKAP1 (-0.05) and between C3 and THEMIS (-0.06) (***Supplementary Fig. 14C***). The positive correlation between C3 and IL1B suggests that complement activation and inflammasome signaling are co-regulated in SPP1⁺ MG, linking the “Eat-Me” phagocytic axis with pro-inflammatory cytokine production.

Pathway enrichment revealed that SPP1⁺ MG gained inflammatory, immune regulatory, cell adhesion, and neuronal interaction functions, while homeostatic functions were lost, consistent with a shift from a resting surveillance state toward an activated, inflammatory phenotype (***Supplementary Fig. 14D***).

### SPP1⁺ MG Associated with Aberrant DNA Damage Response (DDR) Of Neuronal Injury Under ART Suppression

To assess the impact on neuronal health, we analyzed genes involved in core neuronal functions (**Fig. 7A**). A progressive decline of neuronal pathways was observed in ART-suppressed SPP1⁺ MG-targeted receptors, affecting synaptic transmission and plasticity, neuronal development and migration, and calcium signaling. The broad suppression of neuronal function-related pathways suggests that neuronal homeostasis is not restored despite viral suppression.

**Figure 7.**
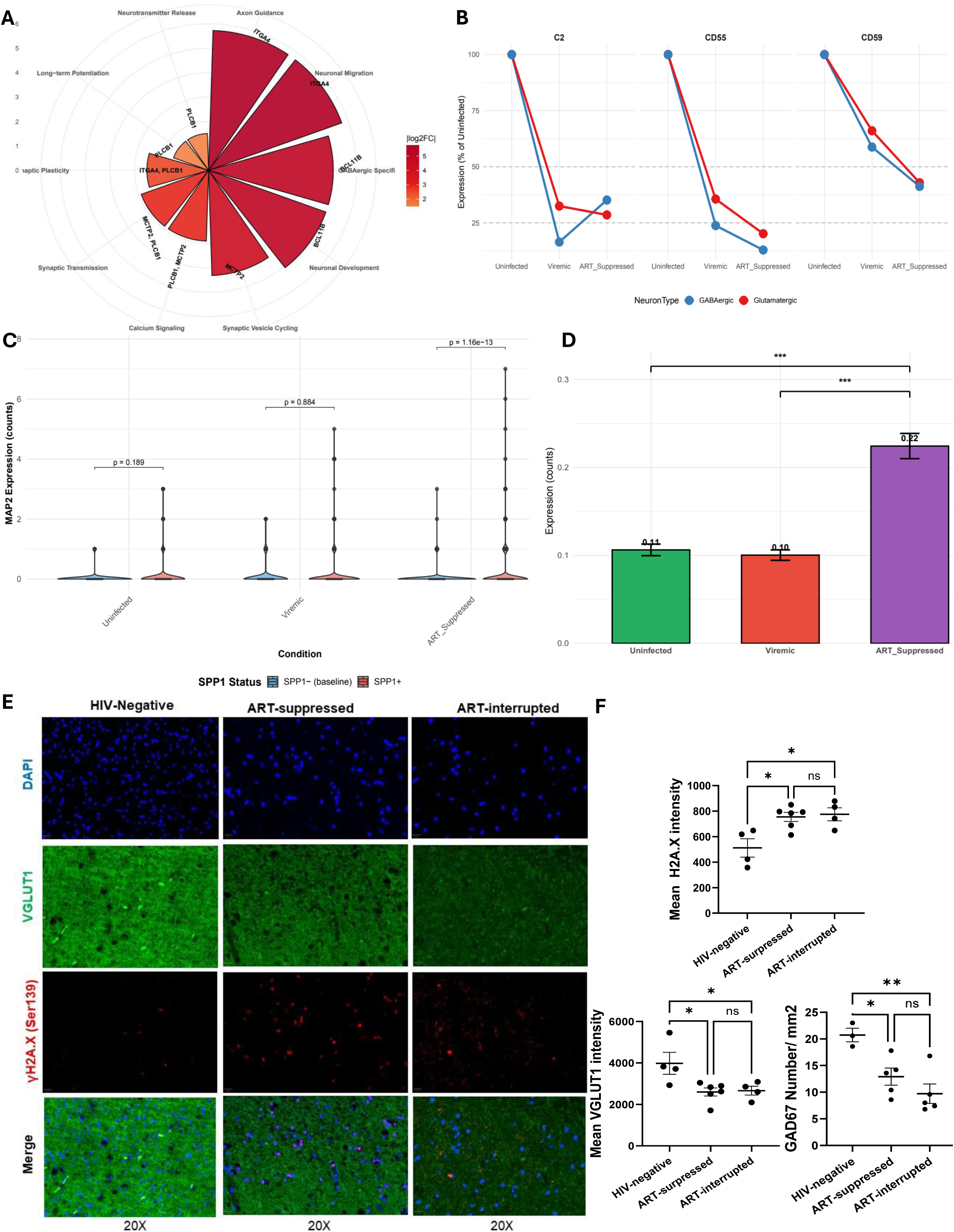
SPP1⁺ MG Drive Neuronal Dysfunction, MAP2 dysregulation, and DNA damage response to drive the loss of VGLUT+ and GAD67+ neurons on ART. **(A)** Circular plot illustrating lost neuronal functions in ART-suppressed brains, including axon guidance, long-term potentiation, neurogenesis, synaptic plasticity, and T cell receptor signaling. **(B)** Gradual decrease of complement inhibitor CD59 in neurons from uninfected to viremic to ART-suppressed states, expressed as percent of uninfected baseline. **(C)** MAP2 expression in SPP1⁺ versus SPP1⁻ MG across uninfected, viremic, and ART-suppressed conditions, with statistical comparison using SPP1⁻ as baseline. **(D)** Aberrant MAP2 expression in GABAergic and glutamatergic neurons across conditions in donor brains. (**E-F**) IHC analysis of DSB biomarker γH2AX, excitatory neuron biomarker VGLUT1, and inhibitory neuron biomarker GAD7 expression among frontal cortex brain tissues from HIV-negative, ART-suppressed, and ART-interrupted donors.

A gradual and progressive decrease in complement inhibitors (C2, CD55, CD59) was observed from uninfected to viremic to ART-suppressed states in both GABAergic and glutamatergic neurons, suggesting that neurons progressively lose their protection against complement-mediated damage. This neuroprotective decline is not reversed by ART. Quantification of neuronal abundance revealed substantial loss of both GABAergic and glutamatergic neurons under viremic conditions, with GABAergic neurons showing the most severe reduction (as low as 10-18% of uninfected levels) (**Fig. 7B**). Under ART suppression, both neuronal populations remained significantly below baseline (GABAergic: 10-45%; glutamatergic: 28-42%), indicating that viral suppression does not fully restore neuronal abundance. Together, these findings suggest that progressive loss of complement inhibitors may contribute to ongoing complement-mediated neuronal injury and depletion, even under suppressive ART.

To investigate the association between SPP1⁺ microglia and neuronal injury, we compared MAP2 expression in the presence of SPP1⁺ versus SPP1⁻ microglia (**Fig. 15C**). No significant difference was observed in uninfected (p = 0.189) or viremic (p = 0.884) donors. In contrast, under ART suppression, a highly significant increase in MAP2 expression was associated with SPP1⁺ microglia (p = 1.16e-13), suggesting that the association between SPP1⁺ microglia and altered MAP2 expression is observed only in of long-term ART. Under ART, MAP2 expression is elevated in the presence of SPP1⁺ microglia (**Fig. 7C**), but this occurs alongside complement activation, loss of neuronal complement inhibitors (CD55 loss 87%, **Fig. 7B**), and reduced MAP2 expression in glutamatergic and GABAergic neurons (**Fig. 7D**).

Therefore, MAP2 upregulation in the presence of SPP1⁺ microglia is consistent with reactive neuronal stress rather than neuroprotection, and neurons remain vulnerable to complement-mediated damage. Aberrant MAP2 expression in microglia (**Fig. 7C-D**) further supports this interpretation, consistent with our recent report in HAND brains (*12, 48–50*).

As reflected by the enrichment of SPP1 in DNA damage repair pathways (***Supplementary Fig. 2D***), these pathways were associated with IFN (**Fig. 5D**) and HIV (**Fig. 5H**)-induced neuroinflammation as well as neurodegenerative diseases such as multiple sclerosis. Enrichment of the complement system in HIV⁺ MG on ART suggested dysregulated DNA damage response (DDR) and neuronal injury pathways in the CNS. Transcriptomic data showed that DDR genes (ATR, BLM, CHEK2, PARP1, RAD51C, RAD51D) were upregulated (compared to uninfected) in activated MG but downregulated in glutamatergic and GABAergic neurons (**Supplementary Fig. 15**), demonstrating cell-specific regulation. This opposing regulation highlights a fundamental difference in cellular strategy: activated MG, oligodendrocytes, and OPCs prioritize genomic maintenance, whereas glutamatergic neurons appear to prioritize functional output over specific DNA repair pathways. This was supported by the previously observed complement system in the protein levels in brains from PWH on ART with HAND in vivo (*12*). Furthermore, γ-H2AX, a double-strand break (DSB) biomarker, was significantly increased following ART interruption and persisted despite ART, leading to loss of VGLUT1⁺ and GAD67⁺ neurons under ART suppression (**Fig. 7E-F and *Supplementary Fig. 15***).

SPP1⁺ activated MG subset sustains a pathological state characterized by chronic IFN activation and MHC-I upregulation. These inflammatory and immune-evasive MG are associated with dysregulated DDR signaling in glutamatergic and GABAergic neurons, leading to synapse dysfunction and disrupted neuronal signaling in ART-suppressed brains.

## Discussion

Our study uncovers a distinct SPP1^+^ activated MG reservoir subset in ART-suppressed PWH, characterized by heightened MHC-I–mediated immune evasion by HIV Nef. These features were tightly linked to persistent IFN signaling activation, as we recently reported in PWH with HAND (*12*). These activated MG subset cells exhibited an inflammatory signature enriched for SPP1^+^ disease-associated MG (DAM) signaling (*51*), which may be mediated *via* SPP1/ITGAV/ITGB interactions and is crucial for MG-mediated neuroinflammation even in the presence of ART. Of note, SPP1^+^ MG-mediated neuroinflammation causes dysregulated DDR signaling and TREM2/C1Q “Eat me” signaling, thereby enhancing DSBs and inducing neuronal injury on ART.

SPP1 regulates HIV replication in myeloid cells and is markedly expressed in the CNS of both humans and non-human primate models in the setting of HIV infection (*52–55*). The compositional patterns revealed by chi-square analysis provide biological insight. Viremic donors show depletion of activated microglia (residual -28.59), consistent with viral cytopathic effects. ART-suppressed donors show expansion of activated microglia (residual +36.22), driven by the emergence of an SPP1^+^ subset (51.9-98.8% of activated microglia). We propose that SPP1^+^ microglia represent a survival-adapted reservoir that resists depletion seen during viremia and persists under ART, maintaining HIV persistence and driving neuroinflammation despite viral suppression.

Consistent with these observations, enriched SPP1 expression in MG from ART-suppressed donors underscores their activated, neurodegenerative phenotype (*56*). Within the DAM network, the SPP1-driven signaling, *via* SPP1/ITGAV/ITGB, dysregulates intercellular communication and perpetuates neuroinflammation that ART fails to resolve. Interestingly, there is also unusual signaling communications SPP1/ITGAV/ITGB or NAMPT/ITGAV/ITGB between activated MG and endothelial cells, possibly involving BBB disruption. Therefore, the HIV^+^/SPP1^+^ activated MG subset may represent the true DAM in PWH, central to NeuroHIV pathogenesis. Importantly, these SPP1^+^ activated MG are closely associated with IFN activation, distinct from homeostatic resident MG, representing an immune-evasive yet proinflammatory state that may sustain neuroinflammation and neuronal injury despite virologic control (*47, 57*).

By hijacking MHC-I pathways, HIV^+^ activated MG subset maintains chronic signaling while avoiding clearance, an adaptation that ensures long-term CNS viral persistence (*44*). HLA-A loci are highly polymorphic, with expression varying by variant (*58*), though our within-donor comparison of HIV^+^ vs HIV^-^ cells controls for allele-specific effects. Unlike ART-interrupted PWH, HIV transcripts in activated MG were dominated by 3’ *nef* RNA in ART-suppressed PWH but lacked other HIV transcripts, reflecting partial, transcriptionally restricted HIV expression, providing us with evidence of ongoing viral transcription within latently infected CNS reservoirs in all these 5 donors. Structural modeling revealed close contacts between Nef residues and the MHC-I α-helical domain, consistent with Nef-mediated blockade of MHC-I trafficking (*41, 59, 60*). The Nef protein was detected in the brain of 50% of PWH with dementia (*61*), and its expression in the brain MG induces myelin impairment and HAND-like pathology *in vivo* (*62, 63*). Together, these data position Nef as an important modulator of NeuroHIV in PWH despite ART.

Emerging evidence implicates genomic instability as a hallmark of neurodegenerative disease, including Alzheimer’s and HAND (*64–66*). Our data revealed widespread dysregulated DDR-associated genes, particularly in MG, oligodendrocytes, and OPCs, which are critical for myelin integrity. While most DNA repair genes were upregulated, the DSB repair gene *BLM* was selectively downregulated in glutamatergic neurons, aligning with only partial glutamatergic neuronal recovery and the loss of VGLUT1^+^ excitatory neurons despite ART. This could be due to persistent Nef transcripts-activated IFN-I/NRLC5/IL-1β inflammasome, thereby inducing neurotoxic CXCL10/IP10 cytokine in MG (*67, 68*) to cause excitotoxicity and neuronal apoptosis *in vivo* (*69–71*). This is associated with enhanced SPP1/TREM2/C1Q “Eat me” signaling and aberrant SPP1/NRG/NRXN/NCAM communications of neurodegeneration. Considering that glutamatergic neurons make up to 80% of the brain’s neuronal population and are critical for cognition, learning, and mood, the failure to restore normal function likely contributes to the neurocognitive symptoms characteristic of HAND (*57, 72*).

We analyze ART as a binary exposure (suppressed vs. viremic) rather than performing regimen-level pharmacokinetic analysis. This approach is intentional: PWH typically changes ART regimens multiple times over their lifetime due to tolerability, adherence, or drug-drug interactions, making any single ‘last regimen’ an incomplete reflection of cumulative CNS exposure (*73*). Furthermore, definitive pharmacokinetic-pharmacodynamic analysis would require prospective sampling from a single patient from therapy initiation—impossible in postmortem studies. Thus, our central biological question is whether viral suppression normalizes MG transcriptional states, not which specific regimen achieves suppression.

We acknowledge age differences as a limitation of postmortem clinical samples. We cannot control age at the end of life; however, the SPP1^+^ signature was present across all ART-suppressed donors regardless of age. Also, postmortem interval and comorbidities are potential confounders in postmortem human brain studies; despite that, the SPP1^+^ MG signature was consistently observed across all ART-suppressed donors regardless of these variables.

Together, our study defines a unique immune reprogramming within the SPP1⁺/activated HIV⁺ MG subset in brains from ART-suppressed PWH, linking persistent IFN activation, neuroimmune evasion, and genomic injury across MG, oligodendrocytes, and glutamatergic neurons, leading to neuronal injury (***Supplementary Figure 16***). Thus, targeting aberrant SPP1 signaling in brain MG may represent a potential therapeutic avenue to limit neuroinflammation and potentially eliminate latent HIV reservoirs in the CNS of PWH on ART, as evidenced by a recent study *in vivo* (*74*).

## Data Availability Statement

All relevant data are within the manuscript and its Supporting Information files.

## Contributions

GJ conceived the study. CSS designed the custom reference genome and performed sc/snRNA-seq data analysis. YT performed the approaches to characterize brain tissue and cell samples and developed and coordinated the experiments for sc/snRNA-seq, bulk RNA-seq, immunostaining, and RNAscope. YT, AM, and EPB isolated single cells and/or single nuclei for sc/snRNA-seq analyses. HC treated the cells and prepared MG RNA for bulk RNA-seq. NK and CSS performed RT-qPCR validation in purified MG. JBL and HC performed IHC. AC and SG coordinated patient recruitment, performed rapid research autopsies, and shipped samples to UNC. DMM provided institutional support. CSS prepared the initial manuscript draft. CSS and GJ discussed, revised, and edited multiple versions, while YT, AC, SG, and DMM contributed to manuscript editing. GJ finalized the submitted version. All authors approved the final manuscript.

## Acknowledgements and Financial Disclosure Statement

We thank the participants who donated their tissues for this study. We thank Zhaojiayi Zhang and Madhura Manjunath for their work on IHC. This work was supported by R21MH128034, R01MH136852, R21AI167709, R01AI186609, and R01MH139466-01A1 (to GJ); UM1AI164567 (to DMM); the Collaborative Development Program at B-HIVE (U54AI170855 to GJ), and UNC School of Medicine supplemental funding (to YT). Additional supports were provided by the Translational Virology Core at the San Diego Center for AIDS Research (P30AI036214), the James B. Pendleton Charitable Trust, and the “Last Gift” cohort (P01AI169609).

## Materials and Methods

### Ethics Statement

All study protocols were approved by the Institutional Review Boards (IRBs) of the University of California, San Diego (IRB no. 160563), the National Disease Research Interchange (NDRI), and the National NeuroAIDS Tissue Consortium (NNTC). Participants enrolled in the Last Gift cohort were altruistic, terminally ill PWH with <6 months expected life, without CNS malignancy and immune checkpoint therapy. Written informed consent was obtained before postmortem tissue collection and autopsy procedures.

### Brain Tissue Collection and MG Isolation

Snap-frozen brain tissues (∼20 g frontal cortex, basal ganglia, and/or hippocampus) were collected from five ART-suppressed and two viremic PWH, with two HIV-negative individuals serving as uninfected controls (**Table 1**), which were submerged in ice-cold sterile DMEM supplemented with (HIV^+^ tissues) or without (HIV^-^ tissues) ART drugs (200 nM raltegravir, 100 nM nevirapine, and 25 nM darunavir) during transport to prevent ex vivo viral reactivation. Samples were transported on ice to the UNC HIV Cure Center and processed within 24 h. Some tissues were then snap-frozen and stored at −80 °C for further use, while the remainder of the tissues were immediately processed for CNS cell isolation, as we previously reported (*4*).

### γH2AX, VGLUT1, GAD67, and Nef Immunohistochemistry in Brain Tissues

This was performed as previously described (*12*). Briefly, brain tissues were fixed in 10% formaldehyde for one week, then embedded in paraffin, and sectioned into 6 μm-thick slices. Tissue sections underwent deparaffinization using xylene baths, followed by rehydration through ethanol. Subsequently, the slides were exposed to 100% ethanol, followed by protease plus at 40°C for an additional 30 minutes. A hydrophobic barrier was created using a hydrophobic barrier pen to facilitate the efficient application of reagents. The slides were blocked with a mixture of 15% goat serum and 1% FcR human block in TBST for 45 minutes. the primary γH2AX (Cell Signaling Technology, Cat No 2577L,1:1000), HIV-1 Nef (Santa Cruz, Cat No sc-65904 1:500), VGLUT1 (Abcam, Cat No ab242204, 1:5000), or GAD67 (Abcam, Cat. No ab26116, 1:2000) antibody, diluted in PBST) was incubated overnight at 4°C. After three washes with 1XTBST, a biotinylated secondary antibody diluted at 1:800 was incubated for 30 minutes. Following a single 5-minute wash, slides were incubated with peroxidase streptavidin at a 1:500 dilution in TBST for IHC. After a final 5-minute wash, slides were exposed to ImmPACT VIP Peroxidase substrate (Cat. No SK4605) at room temperature for 2 minutes to develop the staining. For immunofluorescence, an Alexa-Fluor 488-conjugated 2^nd^ Ab (1:400) was added and incubated for another 1 hour, followed by washes 3 times with 1XTBST. The reaction was stopped by rinsing with distilled water. DAPI or Methyl Green was used to detect nuclei. The image was further scanned by the VS200 Slide Scanner (Olympus) and analyzed by Qupath (v0.6.0).

### Sc/snRNA-Seq Analysis

a. **Single-Nuclei Isolation** Brain tissues were dissociated into nuclei using Dounce homogenization in nuclei lysis buffer (Sigma, NUC-201), followed by Percoll gradient purification. Filtered homogenates were overlaid on a 1.8 M sucrose cushion and centrifuged at 13,000 g for 45 min at 4 °C. The nuclear pellet was resuspended in diluted nuclear buffer with RNase inhibitor. Final single-nuclei suspensions were filtered through 40 µm strainers and used for 10x Genomics 3′ library preparation.
b. **10x Genomics 3′ Library Construction and Sequencing** Libraries were prepared as we described previously (*12*) using the Chromium Next GEM Single Cell 3′ Reagent Dual Index Kit v3.1 (10x Genomics, CG000315, Rev F), targeting recovery of 10,000 nuclei per sample. Barcoded cDNA was generated via GEM-based reverse transcription and amplified for library construction. Libraries were sequenced on an Illumina NovaSeq S2 (paired-end, 28×90 cycles; dual i7/i5 indices, 10 cycles each). Sequencing quality was assessed using FastQC, and library complexity was evaluated via Qubit dsDNA HS assay and Agilent TapeStation 4200. The data are deposited and available upon request.
c. **Custom Reference Genome Design and Alignment** A combined reference genome was created by merging GRCh38 (human) and HIV-1 sequences. Annotation files were customized to capture all HIV open reading frames, regulatory elements, and antisense transcripts. Reads were aligned using cellranger v9.0.1, using relaxed multi-mapping parameters to enhance viral read detection while preserving cellular alignment specificity. HIV reads were quantified via samtools idxstats and visualized in the Integrative Genomics Viewer (IGV).
d. **“Tissue pooling, alignment, and batch correction.** Three independent sections from the frontal cortex (Brodmann area 9) were collected per donor. Each section was aligned independently using Cell Ranger (10x Genomics). After alignment, the resulting expression matrices from the three sections were pooled by merging the batch matrix files for each donor. Libraries were prepared and sequenced across two batches. Batch effects were corrected using Harmony (*17*) integrated into the Seurat pipeline. Data from all donors were integrated using 2000 highly variable genes and 27 principal components.
e. **Sc/sn RNA-Seq Data Processing.** All datasets were processed using Cell Ranger (v9.0.1) for demultiplexing, alignment, and quantification. Expression matrices were imported into Seurat (v4.3.0) and filtered for quality control using thresholds on mitochondrial content, UMI counts, and nGene/nUMI relationships. Dimensionality reduction was performed using PCA followed by UMAP embedding (cosine metric). Louvain clustering identified transcriptionally distinct populations, which were annotated using ScType (brain-human) and manually validated using canonical marker genes. Doublets were removed using DoubletFinder, and ribosomal, mitochondrial, and cell cycle genes were regressed out. We performed a chi-square test, which revealed a highly significant association (χ² = 10,581, df = 22, p < 2.2e-16). Cramér’s V = 0.291 indicated a moderate effect size, consistent with notable differences in the proportions of several immune populations. Standardized residuals and relative proportion analyses confirmed that ART-suppressed, viremic, and uninfected samples have distinct global immune cell profiles. These results justify proceeding with downstream analyses while accounting for overall compositional differences. We also specifically focused on microglial subsets, including Activated, ISG-Activated, and Resident MG, to investigate condition-specific shifts within this compartment. A focused chi-square test indicated a significant association across conditions (χ² = 159.3, df = 4, p < 2.1e-33). Still, the effect size was smaller (Cramér’s V = 0.08), reflecting more subtle variations in microglial composition than in the global immune landscape. Standardized residuals and relative proportions indicate that, while some subsets are slightly enriched or depleted under specific conditions, the overall microglial compartment remains relatively stable, providing a precise framework for examining microglial-specific transcriptional changes.
f. **Cell–Cell Communication and Ligand–Receptor Signaling Analysis** Cell–cell communication networks were inferred using CellChat (v1.6.1). Normalized Seurat data were used to compute probabilistic ligand–receptor (LR) interactions. Directionality and pathway activation z-scores were compared using CellChat’s differential analysis framework. LR-linked pathway enrichment was mapped to GO biological processes and KEGG canonical pathways using clusterProfiler (v4.14.6), integrating ligand–receptor data with downstream gene ontology. Communication probabilities were calculated using permutations and a significance threshold of p < 0.05. Pathway-level aggregation was performed using ‘computeCommunProbPathway’. Outgoing and incoming signaling strengths were computed for each cell type, and balance scores were calculated as outgoing minus incoming signaling strength. All statistical comparisons were performed using the permutation-based framework implemented in CellChat.
g. **Pathway and Functional Enrichment Analysis** Differentially expressed genes were analyzed with Ingenuity Pathway Analysis (IPA, Qiagen), STRING (v11.5), and clusterProfiler. For each condition, top canonical pathways were ranked by –log10(p-value) and activation z-score.
h. **Structural Modeling of Nef–MHC-I Interactions** Predicted NEF–MHC-I (HLA-A) interactions were modeled using AlphaFold-Multimer (v2.3). Top PLDDT-ranked complexes were visualized in Mol Viewer*, and residue–residue contact distances (< 4 Å) were quantified. Interactions between the Nef tail and the MHC-I α3 domain were highlighted as structural mediators of immune evasion.
i. **HIV Sequence Recovery** Viral transcripts were identified by aligning reads to a curated HIV-1 BED annotation (gag, pol, env, nef, LTRs). Read coverage was visualized in IGV, confirming active transcription across HIV structural and regulatory genes in ART-suppressed donor nuclei.
j. **Ligand–Receptor–Driven Neuronal Pathway Analysis** Ligands and receptors from CellChat were mapped to Entrez IDs (org.Hs.eg.db), followed by GO-BP enrichment (terms containing “neuro,” “axon,” “synap,” “glia”). Networks were visualized using igraph (v2.1.4) and ggnetwork (v0.5.13) to depict neuronal signaling connectivity and crosstalk with microglial ligands. This analysis revealed coordinated neuroimmune modules integrating immune and synaptic communication.

### Bulk RNA-Seq and RT-qPCR Validation of Gene Expression in the Brain MG

Primary human MG were isolated from rapid research autopsies of postmortem HIV^-^ brain tissues, then treated with (n=3) or without (n=3) IFN-β (1 ng/mL) for 5 days to mimic chronic interferon exposure. Then, cells were collected for Bulk RNA-Seq analysis. RNA extraction, library preparation, sequencing, and analysis were conducted at Azenta Life Sciences (South Plainfield, NJ, USA). The data are upon request.

For gene expression validation of bulk RNA-seq data, primary human MG were isolated from postmortem brain tissues, then treated with or without IFN-β (1 ng/μL) for 5 days. Total RNA was extracted using the Qiagen RNeasy Mini Kit, and cDNA was synthesized via the High-Capacity cDNA Reverse Transcription Kit. Gene expression was quantified by qPCR (QuantStudio) using TaqMan Fast Advanced Master Mix, with β-actin as an internal control during qPCR. Differential expression (2^-ΔΔCt) validated transcriptional signatures from bulk RNA-seq analyses.

## DATA AVAILABILITY STATEMENT

The raw and processed single-nucleus RNA-seq data are available upon request.

### Statistical Analysis

All sc/snRNA-seq analyses were conducted in R (v4.3.1). Group comparisons used Wilcoxon rank-sum or Kruskal–Wallis tests with Benjamini–Hochberg correction. FDR-adjusted hypergeometric tests evaluated pathway-level differences. Interaction strengths were ranked using Cell Chat’s permutation-based differential framework. Significance was set at p < 0.05 (FDR < 0.05). For RT-qPCR data, a Student’s t-test was used to assess differences between the control and treated groups; p<0.05 was considered significant.

Single-cell RNA-seq metadata were extracted from the Seurat objects. For each analysis, contingency tables were generated to summarize cell counts per sample and per cell type. Global immune landscape to assess overall differences in cell-type composition across conditions (ART-suppressed, viremic, uninfected), Pearson’s chisquare test was performed, and standardized residuals were computed to identify cell types contributing most to observed differences. Effect sizes were quantified using Cramér’s V, computed with the cramer function from the rcompanion R package.

In the MG-focused analysis, the dataset was subset to include only Activated MG, ISG-Activated MG, and Resident MG. Contingency tables and chi-square tests were computed as above. Standardized residuals and Cramér’s V were again used to quantify condition-specific shifts. Relative proportions of MG subsets were calculated to contextualize observed differences. Multiple testing correction, p-values were adjusted for multiple comparisons using the false discovery rate (FDR) method via the p. adjust() function in R.

## Key resources table

| REAGENT or RESOURCE | SOURCE | IDENTIFIER |
| --- | --- | --- |
| **Antibodies** |  |  |
| Anti-CD68 (mouse mAb) | Santa Cruz Biotechnology | Cat# sc-20060, RRID: AB_628402 |
| Anti-SPP1 (rabbit pAb) | Abcam | Cat# ab274625, RRID: AB_2924848 |
| Anti-HLA-A (rabbit mAb) | Abcam | Cat# ab238438, RRID: AB_2924849 |
| Anti-HLA-E (mouse mAb) | Santa Cruz Biotechnology | Cat# sc-517197, RRID: AB_2924850 |
| γH2AX (rabbit mAb) | Cell Signaling Technology | Cat# 2577L, RRID: AB_2118009 |
| HIV-1 Nef (mouse mAb) | Santa Cruz Biotechnology | Cat# sc-65904, RRID: AB_1135945 |
| VGLUT1 (rabbit mAb) | Abcam | Cat# ab242204, RRID: AB_2924847 |
| GAD67 (rabbit mAb) | Abcam | Cat# ab26116, RRID: AB_449015 |
| Biotinylated secondary antibody (anti-rabbit) | Vector Laboratories | Cat# BA-1000 |
| Biotinylated secondary antibody (anti-mouse) | Vector Laboratories | Cat# BA-2000 |
| Alexa-Fluor 488-conjugated anti-rabbit IgG | Thermo Fisher Scientific | Cat# A-11008 |
| Alexa-Fluor 488-conjugated anti-mouse IgG | Thermo Fisher Scientific | Cat# A-11001 |
| Peroxidase streptavidin | Vector Laboratories | Cat# SA-5004 |
| **Bacterial and Virus Strains** |  |  |
| HIV-1 reference strain | NIH AIDS Reagent Program | Cat# 395 |
| **Biological Samples** |  |  |
| Postmortem human brain tissue (frontal cortex, BA9) | UNC HIV Cure Center / Brain Banks | N/A |
| Primary human microglia (rapid research autopsies) | UNC HIV Cure Center | N/A |
| **Chemicals, Peptides, and Recombinant Proteins** |  |  |
| DMEM (sterile, ice-cold) | Gibco | Cat# 11965-092 |
| Nuclei lysis buffer (NUC-201) | Sigma-Aldrich | Cat# NUC-201 |
| Percoll | Sigma-Aldrich | Cat# P1644 |
| Sucrose (1.8 M) | Sigma-Aldrich | Cat# S0389 |
| Formaldehyde (10%) | Thermo Fisher Scientific | Cat# 28906 |
| Xylene | Thermo Fisher Scientific | Cat# X3P-1GAL |
| Ethanol (100%) | Thermo Fisher Scientific | Cat# 2701 |
| Protease Plus | Agilent | Cat# S2022 |
| Goat serum (15%) | Jackson ImmunoResearch | Cat# 005-000-121 |
| FcR human block | Miltenyi Biotec | Cat# 130-059-901 |
| TBST (1X) | Thermo Fisher Scientific | Cat# 28360 |
| DAPI | Thermo Fisher Scientific | Cat# D1306 |
| Methyl Green | Vector Laboratories | Cat# H-3402 |
| ImmPACT VIP Peroxidase substrate | Vector Laboratories | Cat# SK4605 |
| IFN-β (human) | R&D Systems | Cat# 8499-IF |
| DMSO | Sigma-Aldrich | Cat# D2650 |
| **Critical Commercial Assays** |  |  |
| Chromium Next GEM Single Cell 3' Reagent Dual Index Kit v3.1 | 10x Genomics | Cat# 1000268, CG000315 Rev F |
| Qubit dsDNA HS Assay Kit | Thermo Fisher Scientific | Cat# Q32851 |
| Agilent TapeStation 4200 D1000 ScreenTape | Agilent | Cat# 5067-5582 |
| RNeasy Mini Kit | Qiagen | Cat# 74104 |
| High-Capacity cDNA Reverse Transcription Kit | Thermo Fisher Scientific | Cat# 4368814 |
| TaqMan Fast Advanced Master Mix | Thermo Fisher Scientific | Cat# 4444557 |
| β-actin (TaqMan probe) | Thermo Fisher Scientific | Cat# 4352935E |
| Qiagen DNA Mini Kit | Qiagen | Cat# 51304 |
| **Experimental Models: Cell Lines** |  |  |
| Primary human microglia | UNC HIV Cure Center | N/A |
| **Oligonucleotides** |  |  |
| TaqMan probes for target genes | Thermo Fisher Scientific | Available upon request |
| HIV gag primers and probes | Published | Available upon request |
| Albumin primers and probes | Published | Available upon request |
| **Software and Algorithms** |  |  |
| Cell Ranger (v9.0.1) | 10x Genomics | https://www.10xgenomics.com/support/software/cell-ranger |
| Seurat (v4.3.0) | Satija Lab | https://satijalab.org/seurat/ |
| Harmony (v1.0) | Korsunsky et al. | https://github.com/immunogenomics/harmony |
| CellChat (v1.6.1) | Jin et al. | https://github.com/sqjin/CellChat |
| clusterProfiler (v4.14.6) | Yu Lab | https://bioconductor.org/packages/clusterProfiler/ |
| ScType (brain-human) | lanevski et al. | https://github.com/lanevskiAleksandr/ScType |
| DoubletFinder | McGinnis et al. | https://github.com/chris-mcginnis-ucsf/DoubletFinder |
| Ingenuity Pathway Analysis (IPA) | Qiagen | https://digitalinsights.qiagen.com/products/ingenuity-pathway-analysis/ |
| STRING (v11.5) | STRING Consortium | https://string-db.org/ |
| AlphaFold-Multimer (v2.3) | DeepMind | https://github.com/deepmind/alphafold |
| Integrative Genomics Viewer (IGV) | Broad Institute | https://igv.org/ |
| R (v4.3.1) | R Core Team | https://www.r-project.org/ |
| Qupath (v0.6.0) | Bankhead et al. | https://qupath.github.io/ |
| VS200 Slide Scanner | Olympus | https://www.olympus-lifescience.com/ |
| Agilent TapeStation 4200 | Agilent | https://www.agilent.com/ |
| FastQC | Babraham Institute | https://www.bioinformatics.babraham.ac.uk/projects/fastqc/ |
| PyMOL (v2.5) | Schrödinger | https://pymol.org/ |
| STAR (v2.7) | Dobin et al. | https://github.com/alexdobin/STAR |
| DESeq2 (v1.38) | Love et al. | https://bioconductor.org/packages/DESeq2/ |
| **Other** |  |  |
| CD11b MicroBeads (human) | Miltenyi Biotec | Cat# 130-091-644 |
| LS Columns | Miltenyi Biotec | Cat# 130-042-401 |
| MACS MultiStand | Miltenyi Biotec | Cat# 130-042-303 |
| Countess II Automated Cell Counter | Thermo Fisher Scientific | Cat# AMQAX1000 |
| NovaSeq 6000 | Illumina | N/A |
| Chromium Controller | 10x Genomics | N/A |

