## Supplementary Figures and Tables for "SPP1+ Microglia Are Associated with Neuroimmune Rewiring and Glutamatergic Neuronal Injury in ART-Suppressed People Living with HIV" (from another version of this preprint: version 1, 2026-09-08)

**Supplementary Figure 1. Ligand–Receptor Pairs, Probabilistic and Communication Networks Across Microglial and Glial Populations in ART-Suppressed Donors (Donor 1)**

(**A**) Expression of marker genes used for cell type annotation, showing distinct expression patterns that enabled identification of microglial, astrocytic, and other glial populations in Donor 1.

(**B**) Ligand–receptor (LR) predicted interactions among annotated brain cell types in Donor 1. Each dot represents a significant LR pair identified by CellChat; dot size corresponds to the adjusted p-value, and color indicates the communication probability. Prominent predicted signaling pairs included SPP1–CD44, NAMPT (ITGA5-ITGB1), CCL3/L1-CCR1, and APP-CD74, which are enriched within the ART-suppressed microenvironment. Astrocyte- and microglia-centered hubs were observed, consistent with immune, structural, and repair signaling networks.

(**C**) Expression of SPP1 and CD44 in single-nuclei data. SPP1 expression is higher in myeloid lineage cells, including activated microglia, dendritic cells, and macrophages, while CD44 is highly expressed in resident microglia in Donor 1, suggesting distinct expression patterns between activated and resident microglial subsets.

**Supplementary Figure 2. Ligand–Receptor Pairs and Probabilistic Communication Networks Across Microglial and Glial Populations in ART-Suppressed Donors (Donor 2 Replication) and SPP1-CD44 Enrichment Analysis**

**(A)** Expression of marker genes that aided in cell type annotation, confirming the cellular landscape in Donor 2.

**(B)** Ligand–receptor (LR) predicted interactions among annotated brain cell types in Donor 2. Equivalent analysis for Donor 2 showing reproducible interaction architecture across an independently processed, purified CD11b$^+$ dataset. Each dot represents a significant LR pair; dot size corresponds to adjusted p-value, and color indicates the strength of communication probability. Prominent signaling included SPP1 (SPP1–CD44), NAMPT (ITGA5-ITGB1), CCL3/L1-CCR1, and APP-CD74, which collectively define immune–glial coordination within the ART-suppressed microenvironment. Together, both donors exhibit a consistent hierarchy of high-probability immune and neurotrophic signaling pathways, establishing microglia as central hubs orchestrating chronic neuroimmune communication in ART-suppressed HIV$^+$ brain tissue.

**(C)** Expression of SPP1 and CD44 in single-nuclei data. SPP1 expression is higher in activated microglia, while CD44 was highly expressed in macrophages, suggesting altered functions between activated microglia and macrophages.

**(D)** Gene Ontology biological-process enrichment analysis of SPP1 (red) and CD44 (blue)-associated genes. SPP1 was uniquely enriched for pathways involved in the negative regulation of intrinsic apoptosis and cell survival, whereas CD44 was associated with axon guidance, neuronal projection, and synaptic regeneration. The overlapping enrichment profile highlights a coordinated signaling role of the SPP1–CD44 axis in microglial activation and tissue remodeling.

**Supplementary Figure 3. and Marker Gene Expression**

(**A**) Expression of marker genes that aided in cell type annotation in multimodal annotation, confirming the identity of microglial, astrocytic, neuronal, and other glial populations across conditions.

(**B**) Bar plot showing the absolute number of microglial cells in each subset across Uninfected, Viremic, and ART_Suppressed conditions. Activated Microglia expand from 738 (Uninfected) to 2,605 (Viremic, 3.5-fold) to 4,330 (ART, 5.8-fold). ISG-activated microglia expand from 4 (Uninfected) to 374 (Viremic, 93-fold) to 539 (ART, 135-fold). Resident microglia expand from 7 (Uninfected) to 65 (Viremic, 9-fold) to 82 (ART, 12-fold).

(**C,E**) SPP1 expression levels in activated microglia across all 5 ART-suppressed donors. The higher expression in Donors 1-2 (E) (scRNA-seq) reflects the greater sensitivity of single-cell RNA capture (Mean expression levels ranged from 3.47 to 3.59) (C), which captures cytoplasmic RNA, compared to single-nucleus RNA-seq (Mean expression levels ranged from 2.90 to 3.10), which is restricted to nuclear RNA.

(**D,F**) SPP1+ frequency in activated microglia from Donors 1, 3-5 (snRNA-seq, main cohort, excluding ISG-activated subset): Donor_1: 91.6%, Donor_3: 61.3%, Donor_4: 51.9%, Donor_5: 70.4% (D). SPP1+ frequency in activated microglia from Donors 1-2 (scRNA-seq, preliminary): Donor 1: 98.8%, Donor 2: 89.1% (F).

(**G**) HIV detection in microglia across conditions: ART-suppressed donors showed 0.0808% HIV+ cells (4/4,951 cells), viremic donors showed 0.0986% HIV+ cells (3/3,044 cells), and uninfected donors showed 0% (0/749 cells).

SPP1+ activated microglia are present in all 5 unique ART-suppressed donors, with frequencies ranging from 51.9% to 98.8% across donors and platforms. The signature is observed across both scRNA-seq and snRNA-seq platforms, confirming that no single donor drives the result and demonstrating cross-platform robustness.

**Supplementary Figure 4. Activated and ISG-Activated Microglia Exhibit Divergent Signaling Profiles**

**(A)** Volcano plot of differentially expressed genes (DEGs) in Activated Microglia comparing Viremic vs Uninfected. Red indicates upregulated genes (n = 160); blue indicates downregulated genes (n = 514). Top upregulated genes include the iron transporter SLC11A1 (log2FC: 1.94), interferon-stimulated IFI44L (log2FC: 1.91), scavenger receptor MSR1 (log2FC: 1.76), hypoxia response HIF1A (log2FC: 1.46), and interferon-stimulated EPSTI1 (log2FC: 1.81). Top downregulated genes include homeostatic markers P2RY12 (log2FC: -1.98) and CX3CR1 (log2FC: -2.28), indicating loss of microglial surveillance function.

**(B)** Volcano plot of DEGs in Activated Microglia comparing ART_Suppressed vs Uninfected. Red indicates upregulated genes (n = 50); blue indicates downregulated genes (n = 1,014). Top upregulated genes include SLC11A1 (log2FC: 1.98), cholesterol efflux transporter ABCA1 (log2FC: 1.57), MSR1 (log2FC: 1.47), HIF1A (log2FC: 1.34), and thymidine phosphorylase TYMP (log2FC: 1.43). Top downregulated genes include stress chaperone HSP90AA1 (log2FC: -3.90), myelin protein PLP1 (log2FC: -3.65), and heat shock protein HSPA1A (log2FC: -2.25).

**(C)** Volcano plot of DEGs in ISG-Activated Microglia comparing ART_Suppressed vs Uninfected. Only 1 gene is upregulated—MSR1 (log2FC: 4.07)—while 17 genes are downregulated, including centrosomal protein RTTN (log2FC: -4.18), synaptic plasticity gene SYNDIG1 (log2FC: -4.08), stress chaperone HSP90AA1 (log2FC: -3.65), and DNA damage repair kinase ATM (log2FC: -2.77).

**(D)** Heatmap of gene signatures showing Activated Up (red), Activated Down (blue), and ISG-MG (teal) gene expression patterns across Viremic, ART, and ISG_ART conditions. Rows represent individual genes; columns represent conditions. Color scale indicates log2 fold change from -4 (blue, downregulated) to +4 (red, upregulated). MSR1 is prominently highlighted in the ISG-MG category with the strongest signal.

**(E)** GO Biological Process enrichment bar plot for Activated and ISG-Activated Microglia. Viremic Up: Interferon Response, Inflammation, Phagocytosis, HIF-1 Signaling, Iron Homeostasis. Viremic Down: Synaptic Signaling, Neuron Development, Cell Adhesion, GTPase Signaling, Axonogenesis. ART Up: Lipid Metabolism, Cholesterol Storage, ABC Transporters, Ferroptosis. ART Down: Synaptic Signaling, Neuron Development, Cell Adhesion, MAPK Signaling, PI3K-Akt Signaling. ISG-MG: Lipid Storage, Cholesterol Storage, Phagocytosis, Ferroptosis.

**(G)** KEGG Pathway enrichment dot plot. Circle size represents gene ratio; color represents comparison group. Viremic Up: Ferroptosis, Hepatitis C, Measles. Viremic Down: Synaptic Signaling, Cell Adhesion, MAPK, PI3K-Akt. ART Up: Lipid Metabolism, ABC Transporters. ART Down: Synaptic Signaling, MAPK, PI3K-Akt, FoxO. MSR1 is highlighted as the sole upregulated gene in ISG-MG (4.07-fold), with consistent upregulation in Activated MG under both Viremic (1.76-fold) and ART (1.47-fold) conditions.

**Supplementary Figure 5. Detection and transcriptional divergence of HIV$^+$ microglia in ART-suppressed human brains.**

(**A–C**) Dot plots showing exclusive expression of HIV RNA in HIV$^+$ microglia from Donor 1 (A), Donor 2 (B-C). HIV$^-$ cells display complete absence of viral transcripts, consistent with infection specificity.

(**D–F**) Top differentially expressed genes (DEGs) distinguishing HIV$^+$ and HIV$^-$ microglia across donors. HIV$^+$ cells upregulate stress-response, immune-evasive, and biosynthetic genes (GDF15, PRDM1, ERO1B, HERPUD1), whereas HIV$^-$ cells maintain surveillance and homeostatic programs (CTSB, ATP6V1B2, ID2, SQSTM1).

(**G–I**) Venn diagrams showing minimal overlap between HIV$^+$ and HIV$^-$ DEGs across donors. HIV$^+$ microglia display a larger and largely unique transcriptional signature (Donor 1: 533 vs 83 genes (G); Donor 2 cell 1: 144 vs 164 genes (H); cell 2: 169 vs 65 genes (I); shared = 2–4 genes), consistent with a distinct transcriptional state associated with persistent infection under ART.

(**J**) Fold change of synaptic genes (DLG4, GNB5, HOMER3) in activated MG and glutamatergic neurons across viremic and ART-suppressed conditions compared to uninfected baseline, demonstrating neuronal signaling alterations associated with SPP1$^+$ MG.

(**K–N**) Enrichment of diseases/functions (K), pathways (L), z-score activation (M), and biological processes (N) of HIV$^+$ microglia gene sets. HIV$^+$ microglia show enrichment of immune and metabolic processes, including antigen processing, endoplasmic reticulum (ER) stress, and glycosphingolipid biosynthesis, whereas HIV$^-$ microglia are enriched in apoptotic and cell-migration pathways (K-L). Pathway enrichment (L) and Z-scores (M) indicate enrichment of MHC-I-associated transcripts. (N) Gene Ontology (GO) biological process enrichment among HIV$^+$ microglia upregulated genes. Dominant processes include glycoprotein, carbohydrate, and lipid biosynthesis, suggesting a shift toward anabolic and survival-associated metabolism.

Together, these results reveal that HIV$^+$ microglia display distinct transcriptional profiles compared to HIV$^-$ microglia in ART-suppressed brain tissue, with signatures associated with immune adaptation and metabolic activation.

**Supplementary Figure 6. Detection and transcriptional divergence of HIV⁺ microglia in ART-suppressed human brains. HIV⁺ microglia exhibit distinct ligand–receptor signaling and metabolic profiles compared with HIV⁻ counterparts.**

(**A**) Dot plot showing normalized expression of selected genes distinguishing $HIV^+$ from $HIV^-$ microglia. $HIV^+$ cells show upregulation of genes involved in RNA processing (AGO2, NCSTN), chromatin remodeling (SMARCC1), metabolic homeostasis (ALG11, DLST, AGPAT5), and proteostatic regulation (CTSH). In contrast, $HIV^-$ microglia show minimal expression across these pathways, consistent with a transcriptionally restrained phenotype. The coordinated expression of these genes in $HIV^+$ cells is associated with a survival-driven, metabolically active microglial state in the context of viral persistence and immune-evasive adaptation.

(**B**) Circos plot mapping $HIV^+$ microglia–specific genes to enriched biological processes. Interconnections link immune activation, metabolism, and biosynthesis, featuring key hubs such as CHUK, IKBKG, LY96, and ERAP1, which are associated with inflammatory and lipid-driven transcriptional activity.

(**C**) Ligand expression profiles of $HIV^+$ microglia showing elevated secretion-associated transcripts (SPP1, TGFB1, CCL2, CCL5, CXCL8, IFNG, IL6), consistent with enhanced predicted outgoing signaling capacity. Together, these analyses demonstrate that $HIV^+$ microglia display a coordinated transcriptional profile associated with biosynthetic and immune-evasive states, coupling metabolic remodeling with sustained inflammatory potential consistent with persistent reservoirs in ART-suppressed brains.

(**D**) Receptor expression profiles of $HIV^-$ microglia showing elevated secretion-associated transcripts. $HIV^-$ microglia exhibit markedly reduced expression across these genes, reflecting a subdued transcriptional profile except for IFNGR1. The data show differential activation of cytokine and immune-regulatory modules between infected and uninfected microglia.

(**E**) Heatmap of ligand–receptor (LR) interactions expressed in $HIV^+$ microglia, revealing (scale up to 5) pro-survival and immunoregulatory ligands (SPP1, CCL3, CCL4, CXCL8, CCL2, IL10) paired with receptors (CD44, CCR5, IFNGR1) on $HIV^-$ cells. The strongest predicted interactions were between SPP1, CCL3, CCL4, and CXCL8 and the IFNGR1 receptor, suggesting IFN-associated signaling.

(**F–G**) Enrichment of pathways (F) and biological processes (G) of $HIV^-$ microglia gene sets. $HIV^-$ microglia show enrichment of metabolic maintenance and oxidoreductase-linked processes, autophagy, and degradation (F), including NADPH-dependent enzyme and apoptosis-regulating activities (G). These features are consistent with a preserved but quiescent physiological state.

(**H**) Chord diagram illustrating limited cross-pathway (PLIN2, DHRS7) connectivity in HIV⁻ microglia compared with the more complex immune–metabolic network of HIV⁺ cells. Pathways converge on energy metabolism and proteostasis, but these cells lack the immune feedback loops observed in infected cells.

(**I**) Reversal signaling (predicted HIV⁻ → HIV⁺): HIV⁻ microglia express the same ligand set (CXCL8, CCL3, CCL4), but at markedly lower magnitude (maximum scale ≈ 1), with predicted engagement of receptors (TNFRSF1A, IFNGR1) on HIV⁺ microglia. This subdued predicted communication is consistent with bystander HIV⁻ cells having reduced signaling potential.

(**J**) Expression of CellChat-predicted ligands and receptors in HIV⁺ and HIV⁻ cells in Donor 1 and Donor 2, showing consistent expression of inflammatory ligands in HIV⁺ cells while IFNGR1 was observed in HIV⁻ cells. This expression pattern is consistent with predicted IFN-related signaling between HIV⁺ and bystander cells and is associated with a chronic inflammatory state.

**Supplementary Figure 7. Viremic microglia show transcriptional enrichment without coordinated pathway activation, consistent with decoupled immune and signaling responses.**

(**A**) UMAP projection of single-nucleus transcriptomes showing localization of HIV⁺ nuclei (red) within active microglia clusters.

(**B**) Expression of HIV between HIV⁺ and HIV⁻ active microglia in viremic donors.

(**C**) Venn diagram comparing consolidated HIV⁺ (20 genes) and HIV⁻ (15 genes) microglia-specific transcriptomes in viremic donors. No genes overlapped between groups, reflecting distinct transcriptional profiles.

(**D**) Pathways enriched in genes upregulated in HIV⁺ microglia compared to HIV⁻ microglia in viremic donors. HIV⁺ microglia show enrichment of metabolic pathways, including carbohydrate metabolism (galactose catabolism, glycogen degradation), ketone body metabolism (ketolysis, ketogenesis), lipid biosynthesis (cholesterol biosynthesis, mevalonate pathway, geranylgeranyl biosynthesis), and amino acid catabolism (branched-chain amino acids, isoleucine, tryptophan, histidine). Enrichment of TCA cycle maturation and fatty acid oxidation genes is also observed. These metabolic pathways are associated with an anabolic and energy-producing transcriptional state in HIV⁺ microglia.

(**E**) Pathways enriched in genes in HIV⁻ microglia in viremic donors. HIV⁻ microglia show enrichment of signaling pathways, including nuclear receptor signaling (RAR activation,

TR/RXR activation), G protein-coupled signaling (opioid signaling, G alpha(s) signaling events), cytokine signaling (IL-10 signaling), and neuropeptide signaling (relaxin signaling, apelin signaling). Downregulation of visual phototransduction and NAD signaling is also observed. These pathways are associated with signaling and regulatory functions that are less active in $HIV^+$ microglia.

Together, these data show that in viremic donors, $HIV^+$ microglia display a metabolically enriched transcriptional profile without coordinated activation of signaling pathways. The absence of overlapping gene expression between $HIV^+$ and $HIV^-$ microglia reflects distinct transcriptional states. $HIV^+$ microglia are characterized by enrichment of metabolic pathways, consistent with an anabolic state, while $HIV^-$ microglia show enrichment of signaling pathways. This pattern is consistent with decoupled immune and signaling responses between infected and bystander microglia in the context of active viremia.

**Supplementary Figure 8. Bulk RNA-seq analysis of brain microglia treated with or without IFNβ.**

(**A**) Isolation of brain microglia from HIV-negative donors, which were further treated with 1 ng/mL IFNβ for 5 days to model chronic IFN-I activation.

(**B**) Volcano plot of differential gene expression following IFNβ treatment in microglia, compared with DMSO control. Red = upregulated genes; green = downregulated; grey = not significant.

IFNβ treatment of primary human microglia recapitulates key transcriptional features observed in ART-suppressed $HIV^+$ microglia, including upregulation of SPP1, ISGs, and MHC-I-associated genes. This supports the association between chronic IFN-I signaling and the $SPP1^+$ inflammatory microglial state identified in our human tissue data.

**Supplementary Figure 9. Mechanistic analysis of MHC-I-associated signaling in brain microglia from PWH on ART.**

(**A**) Schematic diagram illustrating predicted HIV-associated remodeling of the NF-κB/ER–Golgi–ubiquitin axis. The diagram shows how FBXW5, SIAH2, and APC/C–linked ubiquitination machinery are connected to IKBKG stabilization and NF-κB activation, which are associated with immune tolerance and cellular survival pathways.

(**B**) Dot plot showing expression of MHC-I-associated genes (CHUK, SEC23A, UBA5, SIAH2, ANAPC10, LY96, FBXW5, IKBKG) in $HIV^+$ and $HIV^-$ microglia. Expression of these genes is observed in $HIV^+$ microglia, consistent with enrichment of MHC-I-associated transcripts.

(**C**) STRING-based protein–protein interaction network linking SPP1, CHUK, IKBKG, and LY96. This cluster connects ER stress and NF-κB signaling, consistent with a coordinated antiviral proteostatic response across the infected microglial network.

(**D**) Protein-protein interaction (PPI) network of MHC-I-associated genes identified in $HIV^+$ microglia, categorized by functional class: antigen processing (blue), immune receptor (pink), immune regulator (purple), membrane trafficking (green), and ubiquitin system components (orange).

**Supplementary Figure 10. HIV+ microglia show coordinated MHC-I pathway activation on ART.**

(**A-B**) Expression of MHC-I genes HLA-A and HLA-E induced during viremic state persisted during ART.

(**C**) HLA-A and HLA-E expression levels remained high in $HIV^+$ activated MG subset during ART suppression.

(D) Protein-protein interaction (PPI) network of MHC-I-associated genes (ERAP1, TAP2, HLA-A, HLA-E, SEC23A, UBA5, SNAP23, IKBKG, CHUK, SPP1, LY96, CD14, FCGR1A, MRC1) identified in $HIV^+$ microglia, categorized by functional response: inflammatory (red) and immune (purple). Immune-associated MHC-I genes (HLA-A, HLA-E) and inflammatory-associated genes (IKBKG, CHUK) are connected to the neuronal gene SNAP23, suggesting potential microglia-neuron communication pathways.

**Supplementary Figure 11. HIV-1 Nef RNA and protein detection by snRNA-seq and IHC and its functional interaction with HLA-A and HLA-E in the brain MG.**

(**A**) Genomic mapping of viral reads aligned to the HIV genome in the brain MG across ART-suppressed donors, compared with ART-interrupted PWH. Viral transcription was detected within the Nef region, including the C-terminal tail, confirming ongoing low-level HIV expression in infected MG.

(**B-C**) Immunohistochemical analysis and quantification show significant protein expression of Nef in viremic donors, which persists even on ART in frontal cortex brain tissues. Red arrow: Nef protein. Scale: 50 μm.

(**D–E**) Structural modeling of Nef–MHC-I complexes. Docking analysis revealed stable interactions between the Nef C-terminal loop (orange) and HLA-A or HLA-E (green), with hydrogen-bond distances ranging from 2.59–3.94 Å. These close contacts suggest that Nef physically engages both classical (HLA-A) and nonclassical (HLA-E) MHC-I molecules, promoting their endocytic internalization and degradation. Zoomed atomic views of Nef–MHC-I interfaces highlighting contact residues within the Nef tail region. The observed interatomic distances (2.6–3.9 Å) are consistent with stable, functional binding and support the model of Nef-mediated MHC-I downregulation through direct structural interference.

**Supplementary Figure 12. HIV+ microglia show synapse dysfunction signaling on ART.**

(**A-B**) Top 20 neuron-related GO-Terms enriched in HIV$^{+}$ (A) and HIV$^{-}$ (B), showing that microglia target glutamatergic neurons' synapses. HIV$^{+}$ MG acts on post-synapses in response to axon injury (C), and HIV$^{-}$ MG focuses on synapse assembly and immunological synapse (B).

(**C**) Functional GO network of neuron-related processes enriched in HIV$^{-}$ MG in ART-suppressed donors, highlighting synapse structure or activity and dendrite activity.

**Supplementary Figure 13. SPP1$^{+}$ microglia dysregulate immune-regulatory, biosynthetic, and damage checkpoint-related genes under ART.**

(**A**) Functional GO network of neuron-related processes enriched in HIV+ MG in ART-suppressed donors.

(**B**) Top DEGs (IL1B, C1QC, TREM2, APOE) confirmed that SPP1+ MG drives neuroinflammation and synaptic pruning in ART-suppressed.

(**C-J**) Comparative expression analysis of immune-regulatory and biosynthetic genes between SPP1$^{+}$ and SPP1$^{-}$ microglia across conditions. Bar plots display mean ± SEM expression values. Genes include MHC-I molecules (HLA-A, HLA-E (C)), MHC-I receptor (TAP2, CTSH (AC), synapse pruning (C1QC, TREM2(D)), amyloid precursor proteins (APP) (NCSTN, AGO2, APP(E)), interferon-response genes (IRF5 (F)), Coenzymes A genes (PPCDC, COASY (G)), Immune checkpoint gene CD274 (PDL1 (H)) and disease-associated microglia (DAM) genes (ITGAM, HEXB (I)) and inflammatory gene (IL1B (J))

**Supplementary Figure 14. Inflamed SPP1+ MG Activated TREM2/Complement Activation to Induce “Eat me” Signaling.**

(**A**) Comparison of SPP1$^{+}$ and SPP1$^{-}$ microglia reveals broad transcription (APOE, SPP1, IL1B, C1QA, TREM2) alteration, including upregulation of DNA repair, chromatin maintenance, and cytokine response genes in the SPP1$^{+}$ population.

**(B)** Complement gene expression showing SPP1$^{+}$ MG dominance in complement activation across all conditions, with C3 expression markedly elevated in ART-suppressed SPP1$^{+}$ MG (5.22 vs. 1.62 in SPP1$^{-}$).

**(C)** Differential expressions of IL1B, PTPRM, SKAP1, SPP1, and THEMIS in activated MG across ART-suppressed SPP1$^{+}$, ART-suppressed SPP1$^{-}$, viremic, and uninfected conditions (log10 scale).

**(D)** Flow of biological changes showing gained versus lost features in SPP1$^{+}$ MG, with inflammatory and immune-regulatory pathways gained and homeostatic functions lost.

**Supplementary Figure 15. DNA damage repair (DDR) gene expression changes across cell types in ART-suppressed and viremic donors, with selective downregulation in glutamatergic neurons.**

Plots showing expression of DNA damage repair genes in viremic and ART-suppressed donors compared to uninfected baseline (dashed line at 1.0).

Activated microglia show upregulation of DDR genes (ATR, BLM, CHEK2, PARP1, RAD51C, RAD51D) with fold changes ranging from 1.1 to 3.0 in both viremic and ART-suppressed donors, indicating enrichment of DNA repair transcripts in this cell type.

Glutamatergic neurons show downregulation of DDR genes with fold changes ranging from 0.20 to 0.50 in both viremic and ART-suppressed donors. GABAergic neurons show less severe downregulation (fold changes 0.20-0.80).

Activated microglia upregulate DDR gene expression under both viremic and ART-suppressed conditions, suggesting maintained or enhanced DNA repair capacity. In contrast, glutamatergic neurons show reduced DDR gene expression that persists in ART-suppressed donors and do not return to uninfected baseline levels. This pattern is consistent with impaired DNA repair capacity in glutamatergic neurons during viremia that is not restored by ART, while microglia maintain a robust DDR transcriptional response.

**Extended Figure 16. Proposed model of SPP1+ activated microglia-associated pathways in PWH on ART.**

This schematic illustrates a proposed model based on transcriptomic observations, computational predictions, and experimental validation from this study.

In ART-suppressed PWH, SPP1+ activated microglia are associated with upregulation of MHC-I-associated transcripts, enrichment of TREM2/C1Q signaling pathways, and activation of DNA damage repair (DDR) gene expression.

The model proposes that these microglial features are connected to synapse impairment and neuronal injury observed in ART-suppressed PWH, including reduced GAD67 expression in inhibitory neurons, downregulation of DNA repair genes in glutamatergic neurons, and persistent γ-H2AX DNA damage signal.

Conclusion: This working model summarizes the associations identified in this study. Causal relationships require experimental validation in future studies.

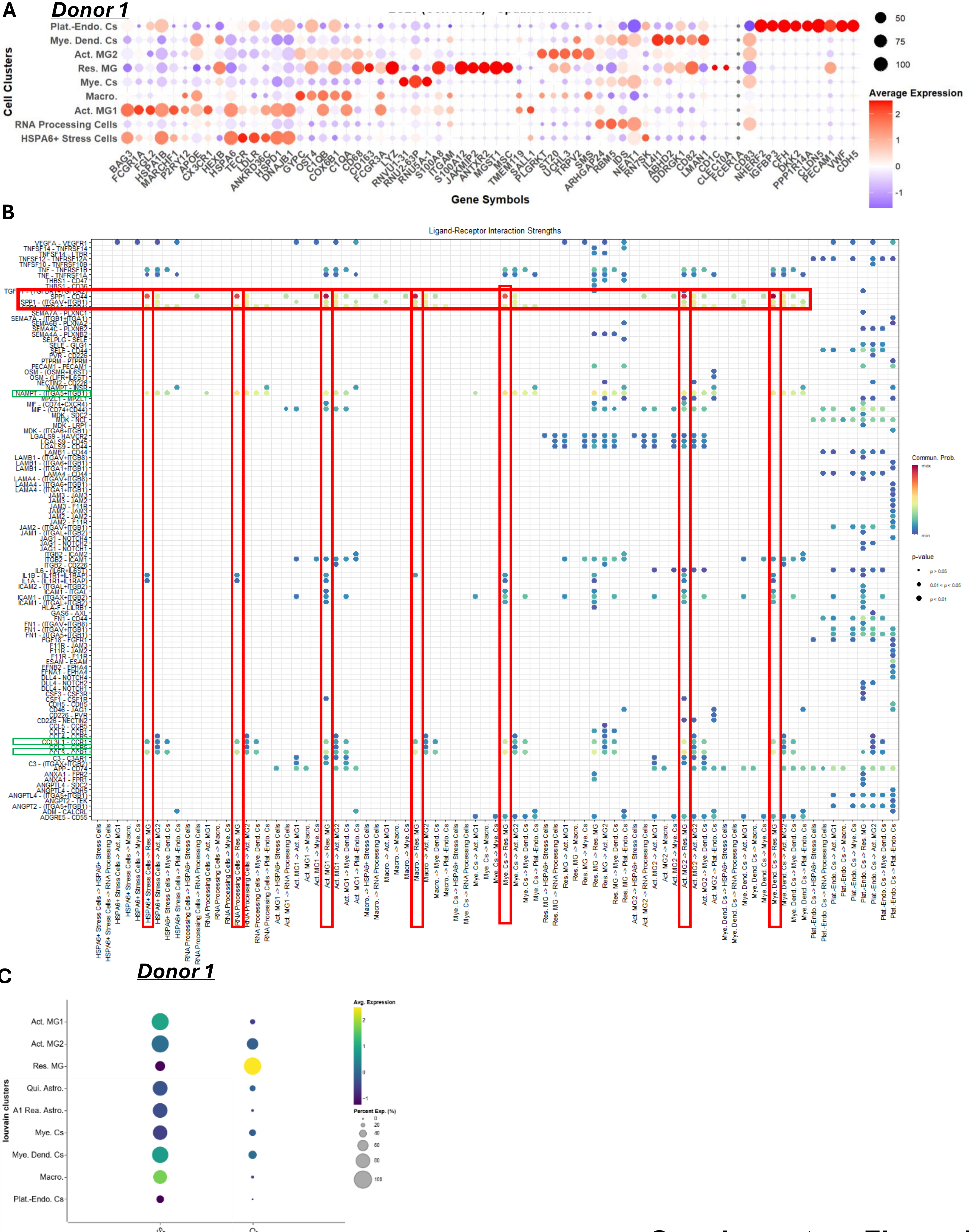


Supplementary Figure 1

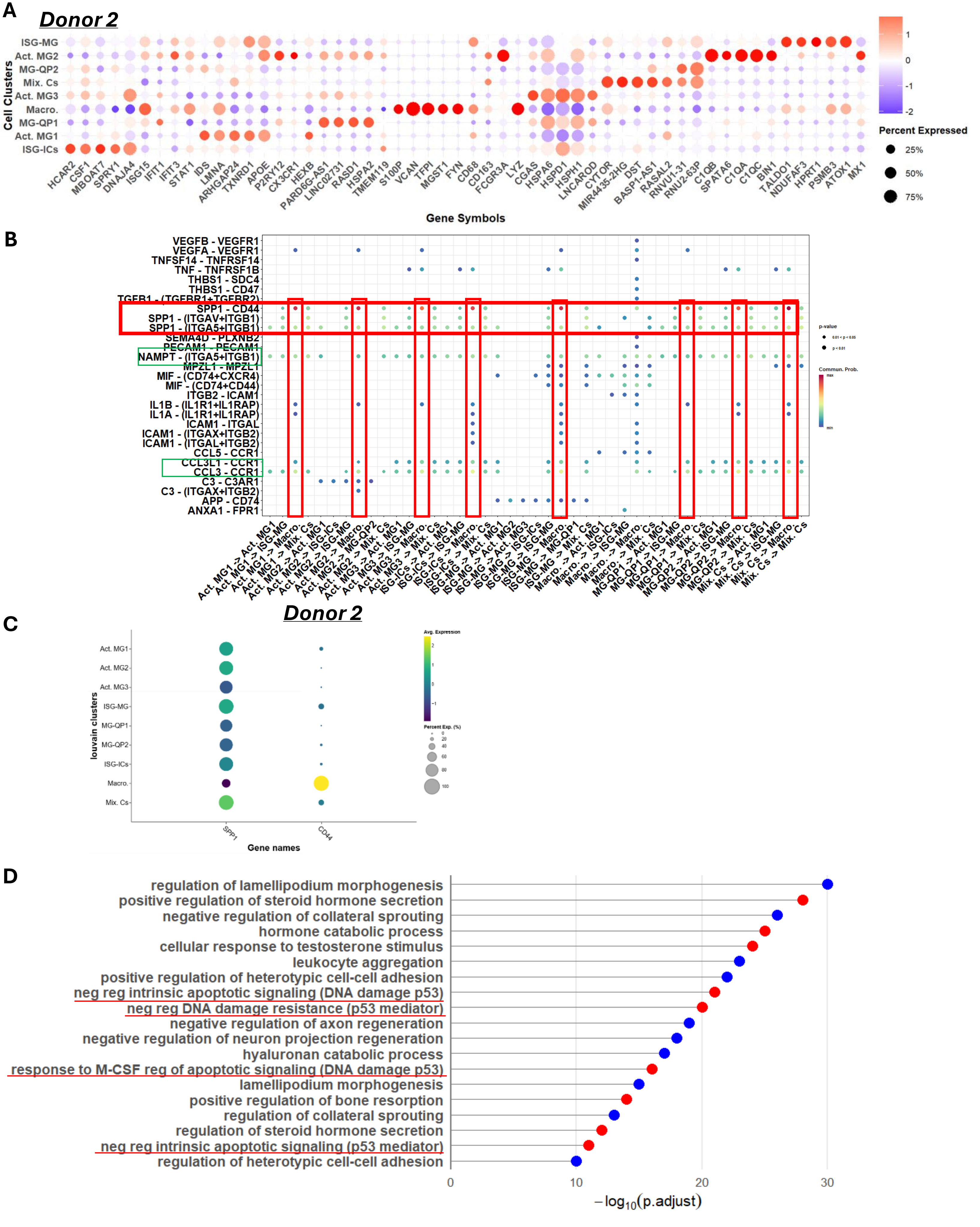


**Supplementary Figure 2**

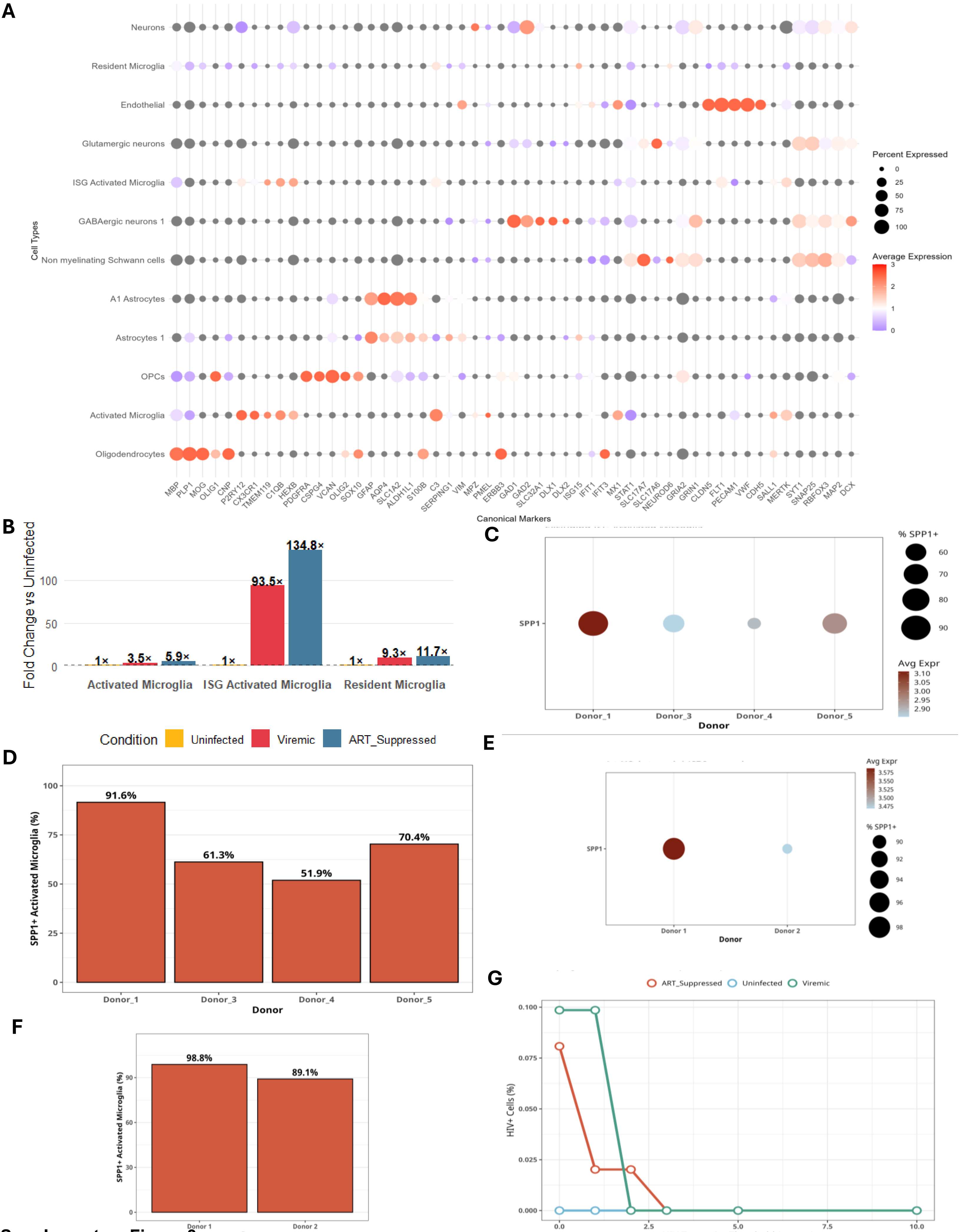


Supplementary Figure 3

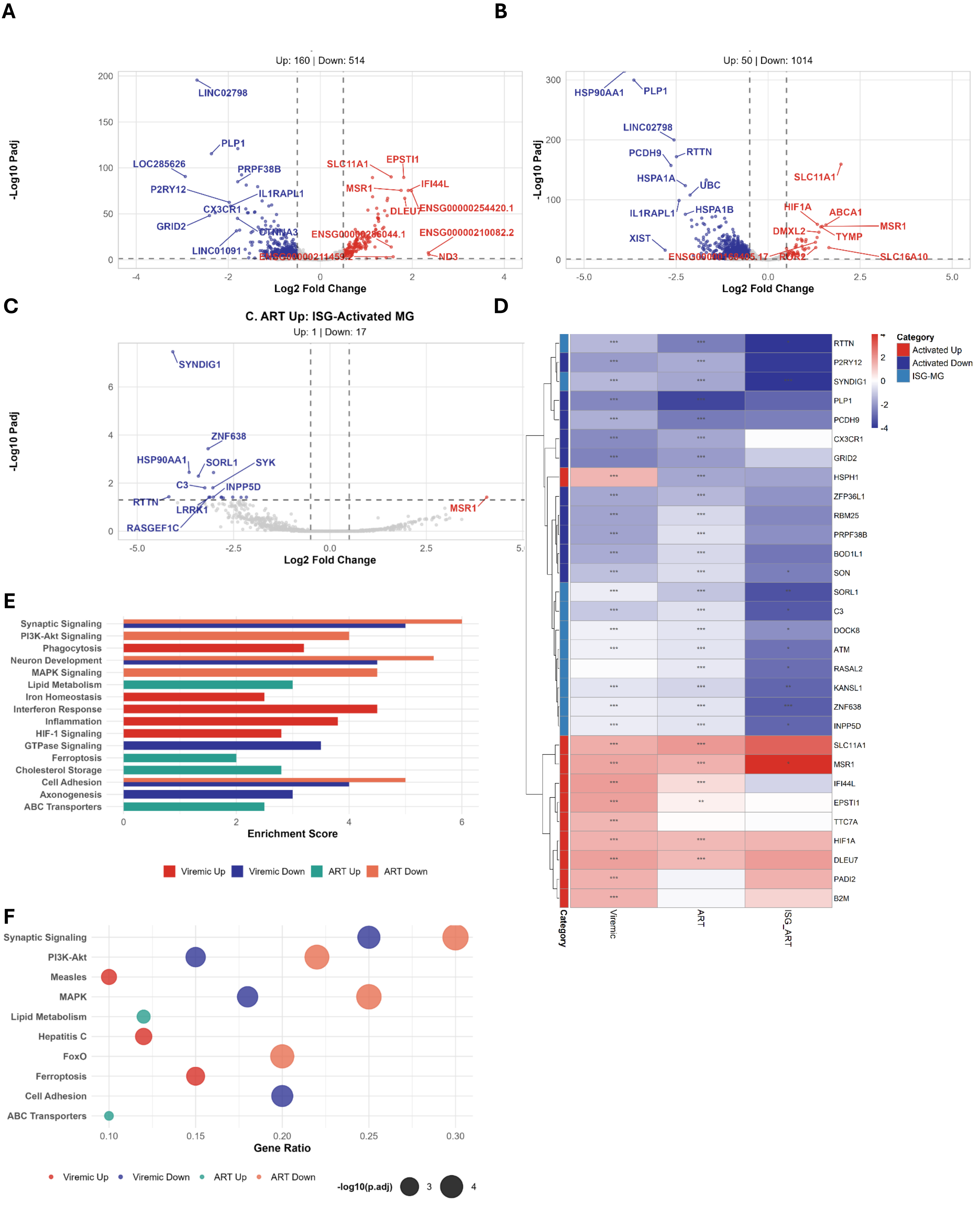


**Supplementary Figure 4**

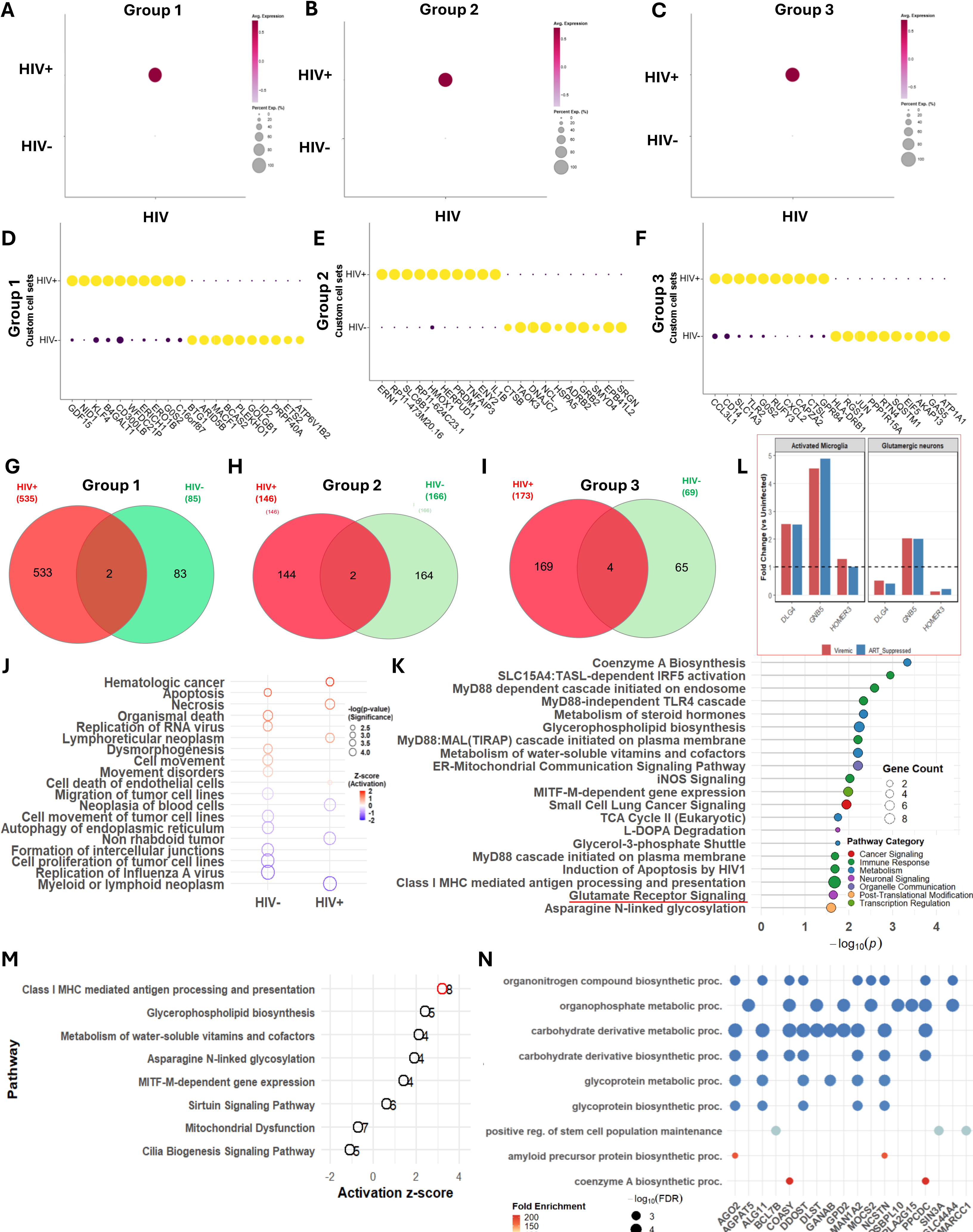


### Supplementary Figure 5

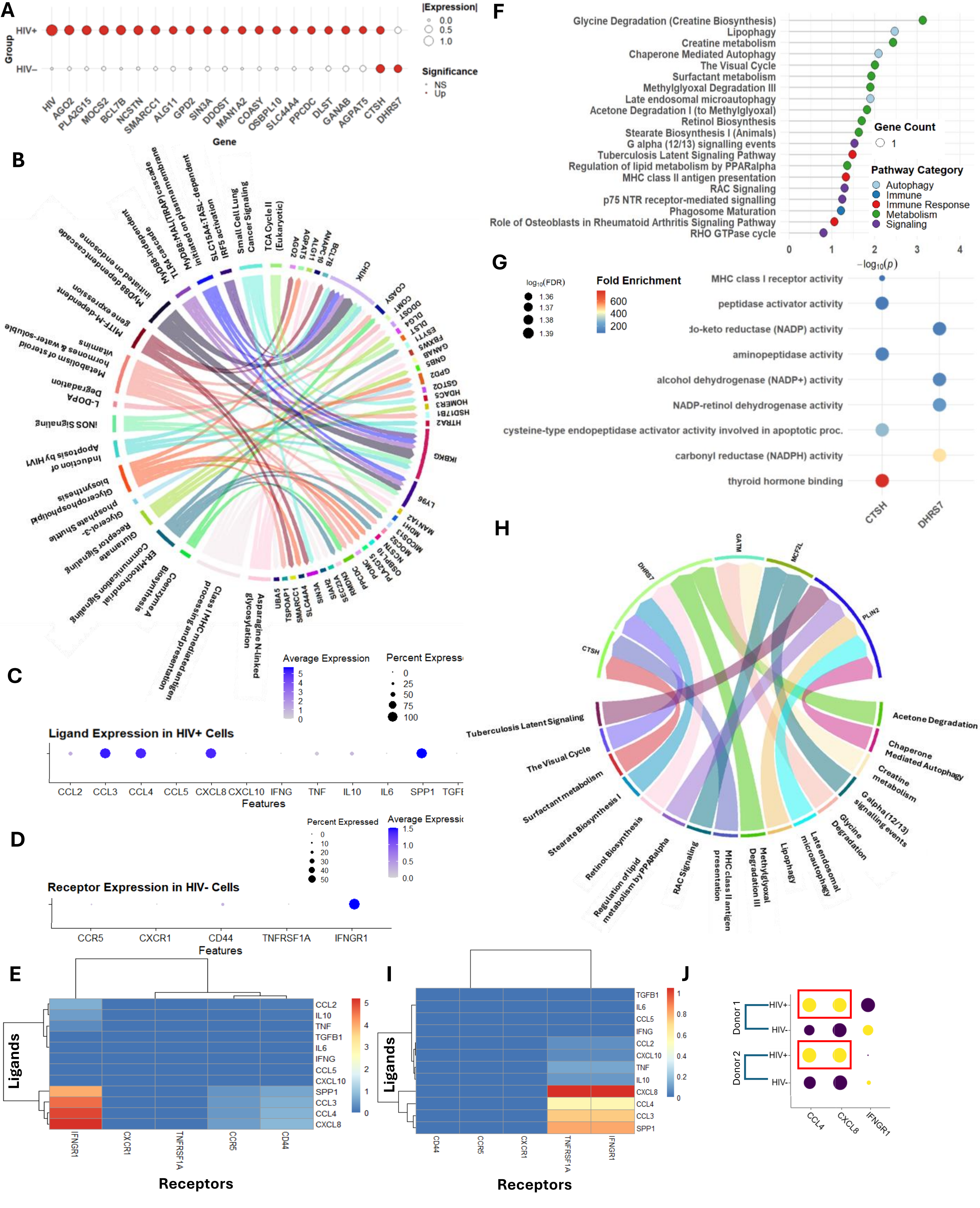

### Supplementary Figure 6

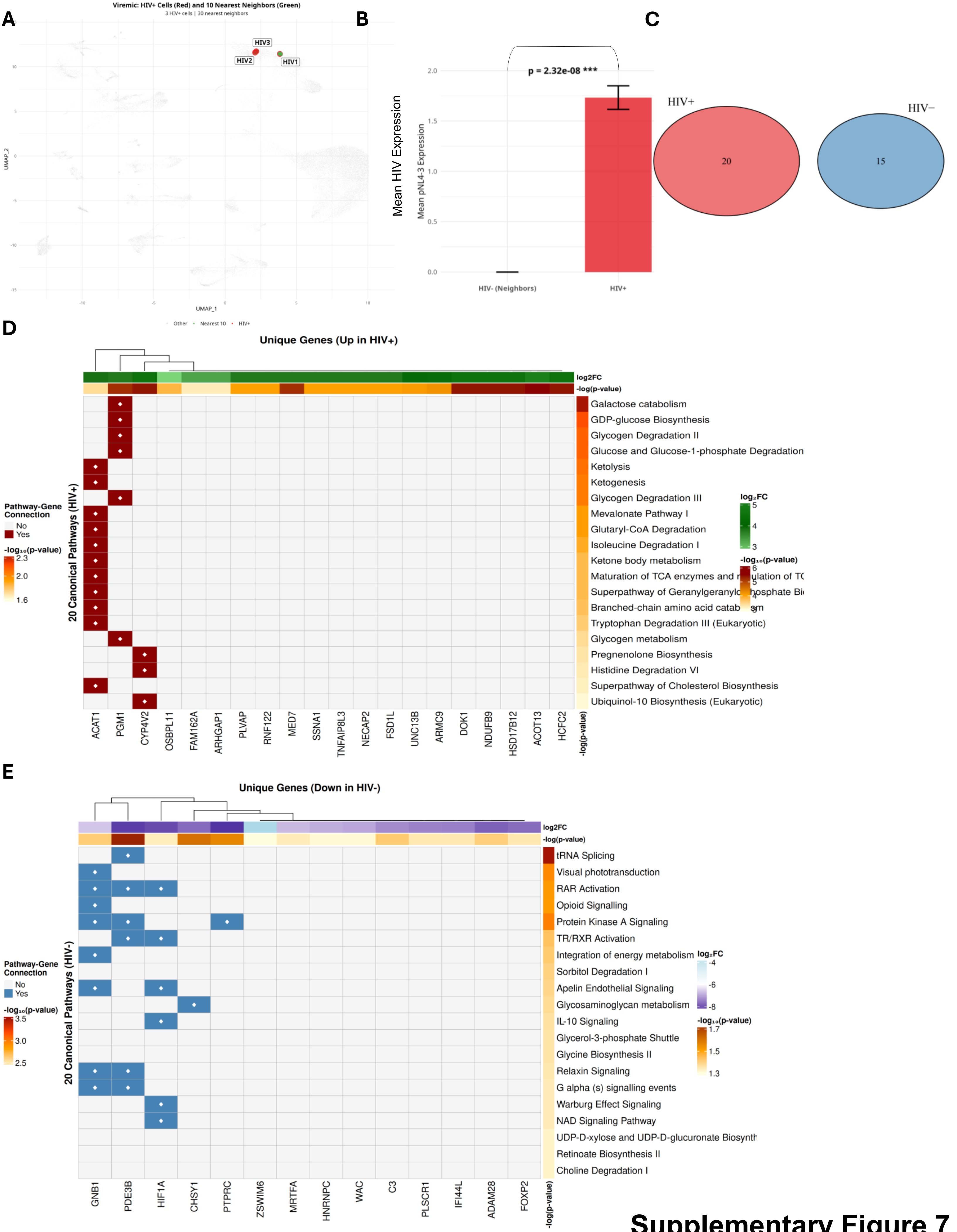


**Supplementary Figure 7**

**A**

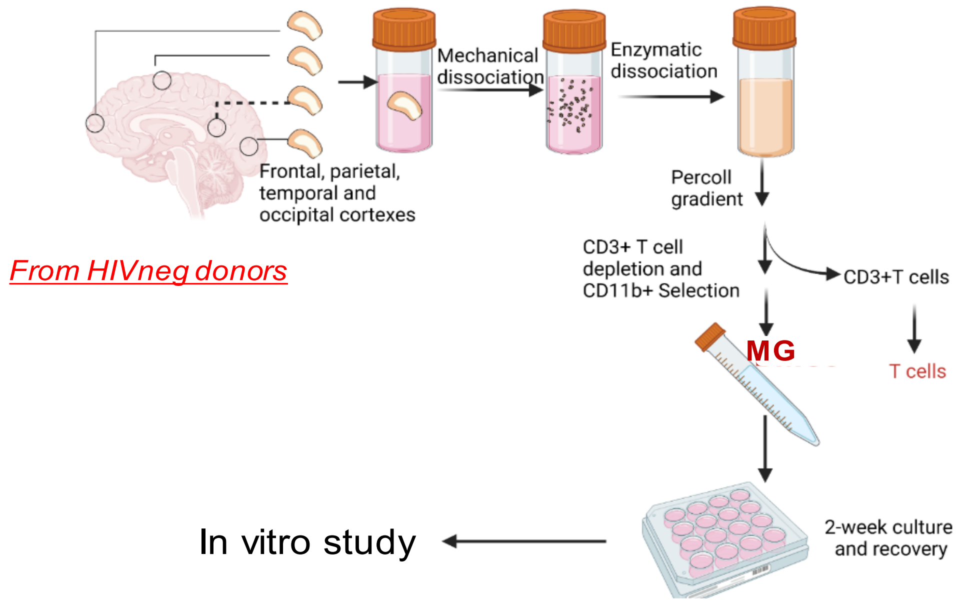


**B**

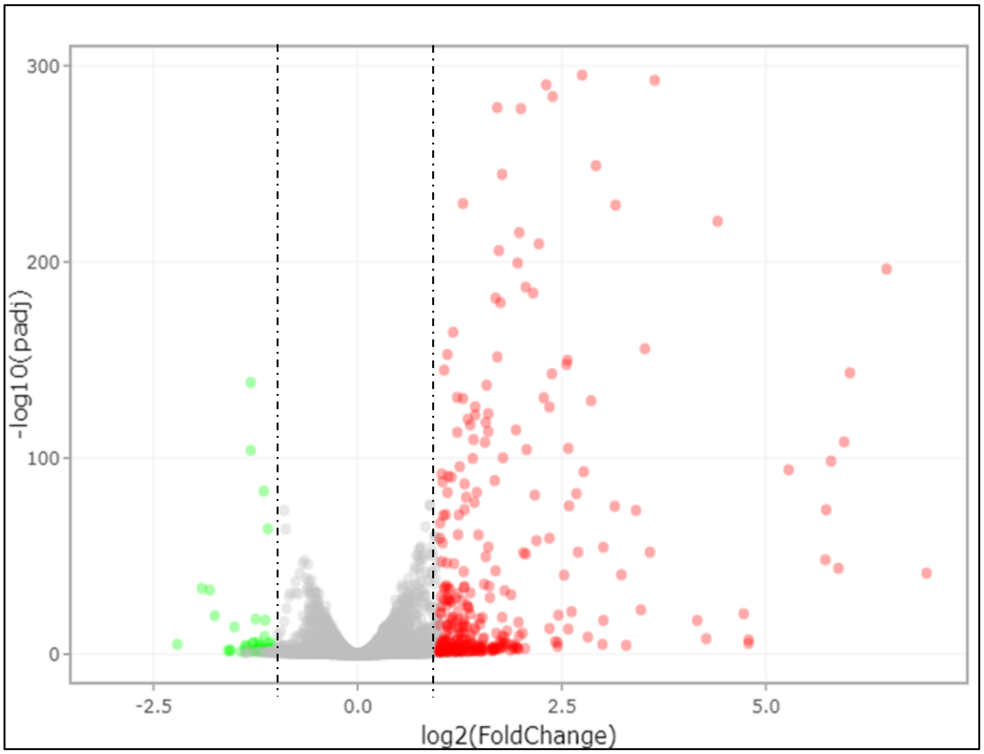


### Supplementary Figure 8

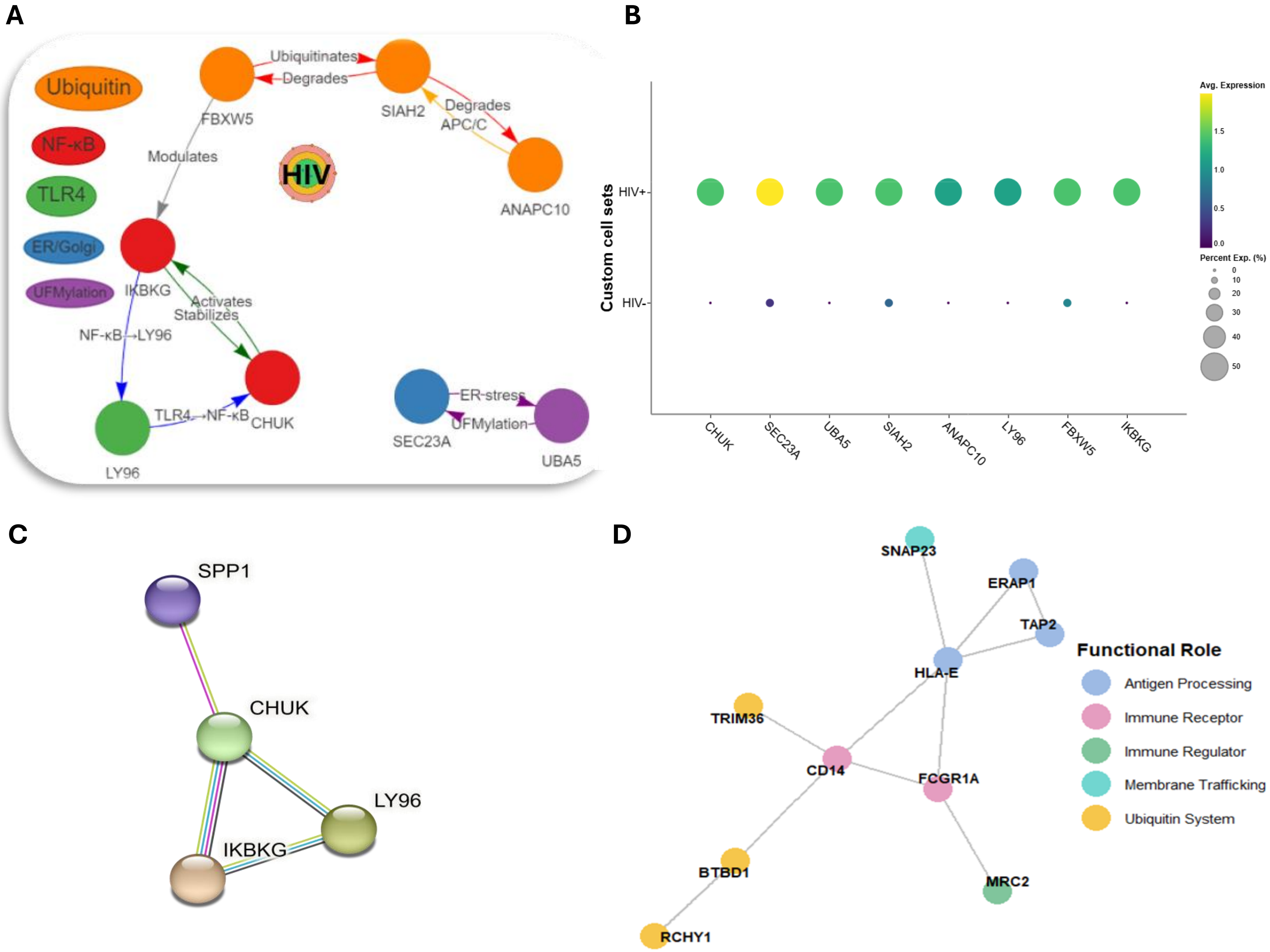


### Supplementary Figure 9

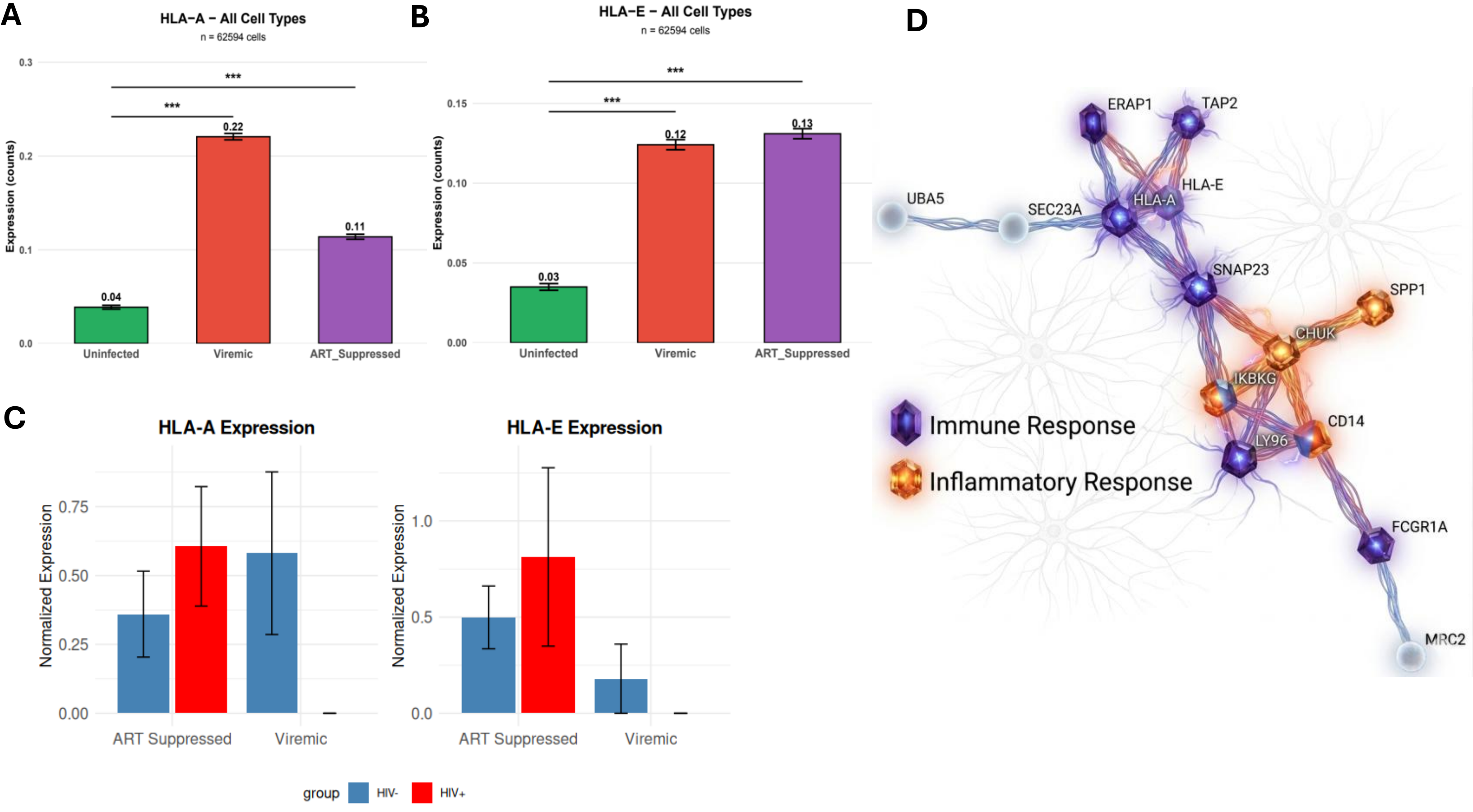


### Supplementary Figure 10

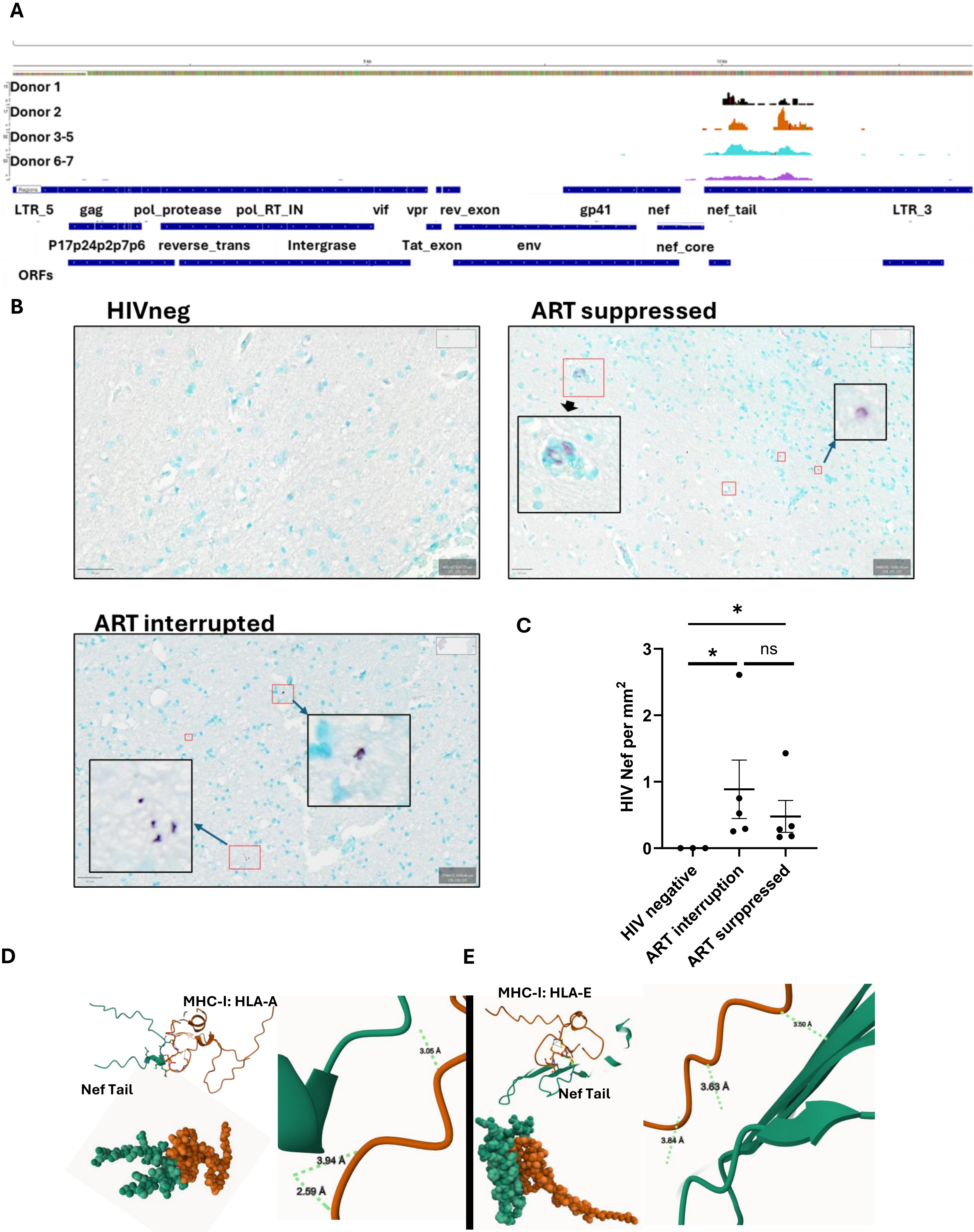


### Supplementary Figure 11

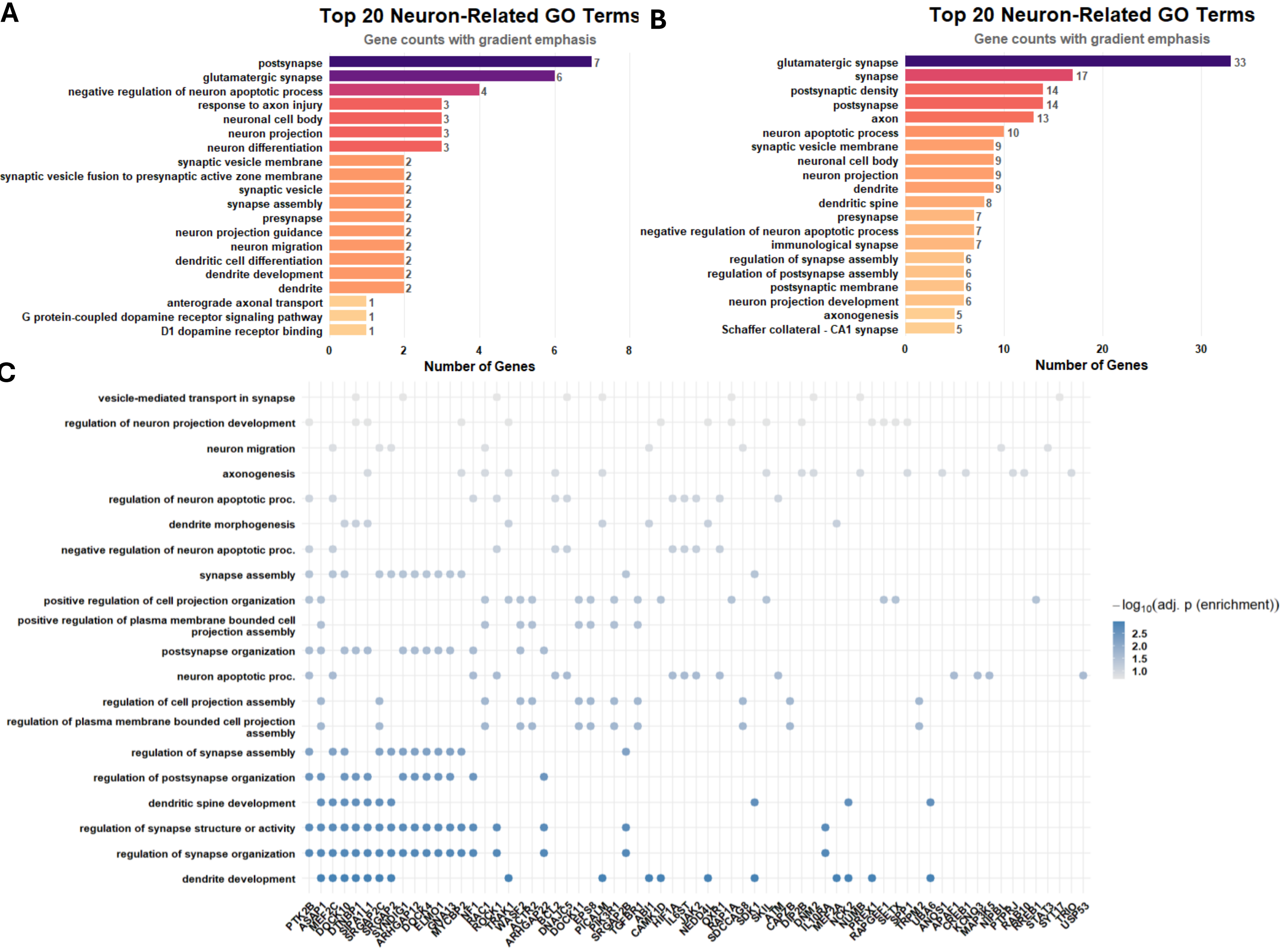


### Supplementary Figure 12

### Supplementary Figure 13

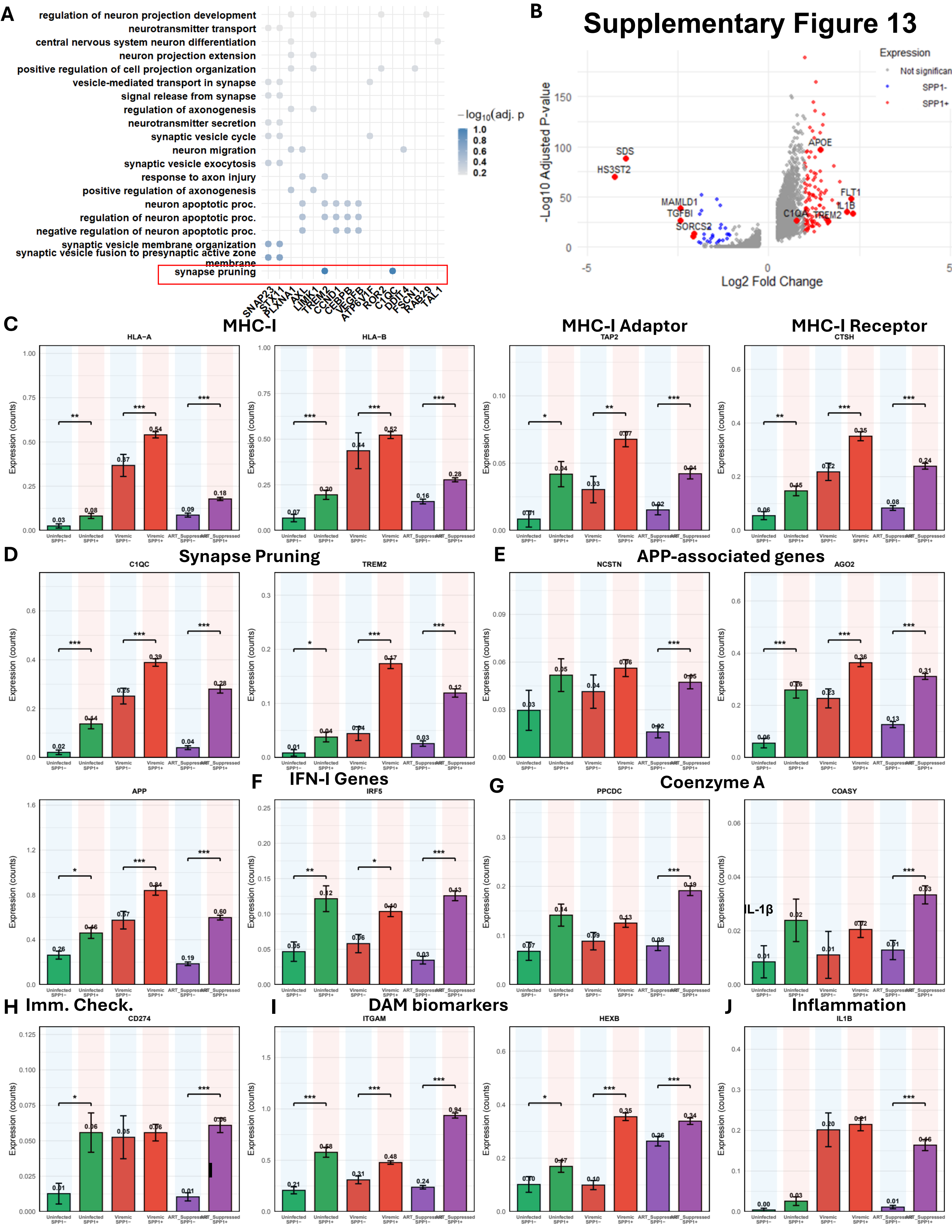

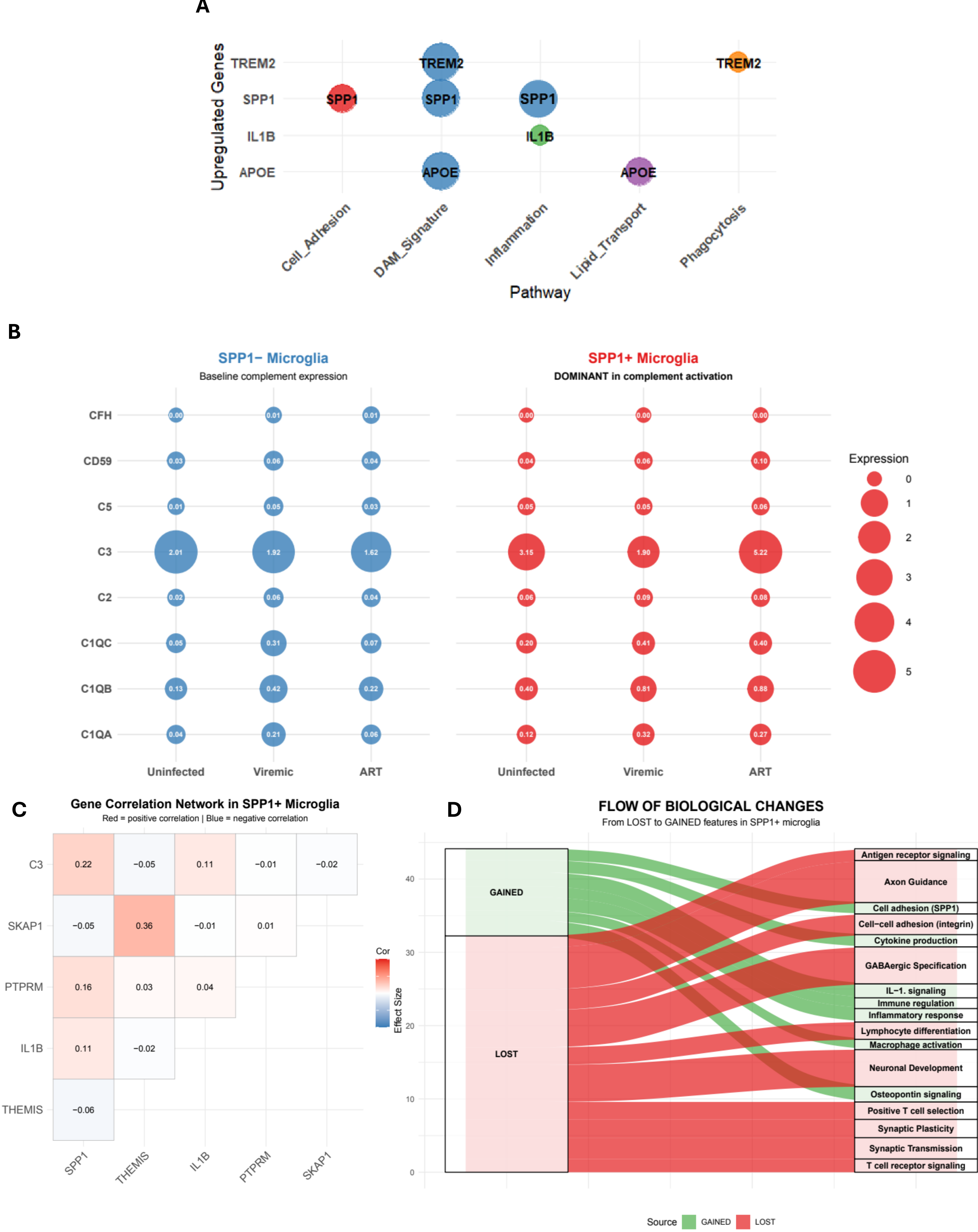


**Supplementary Figure 14**

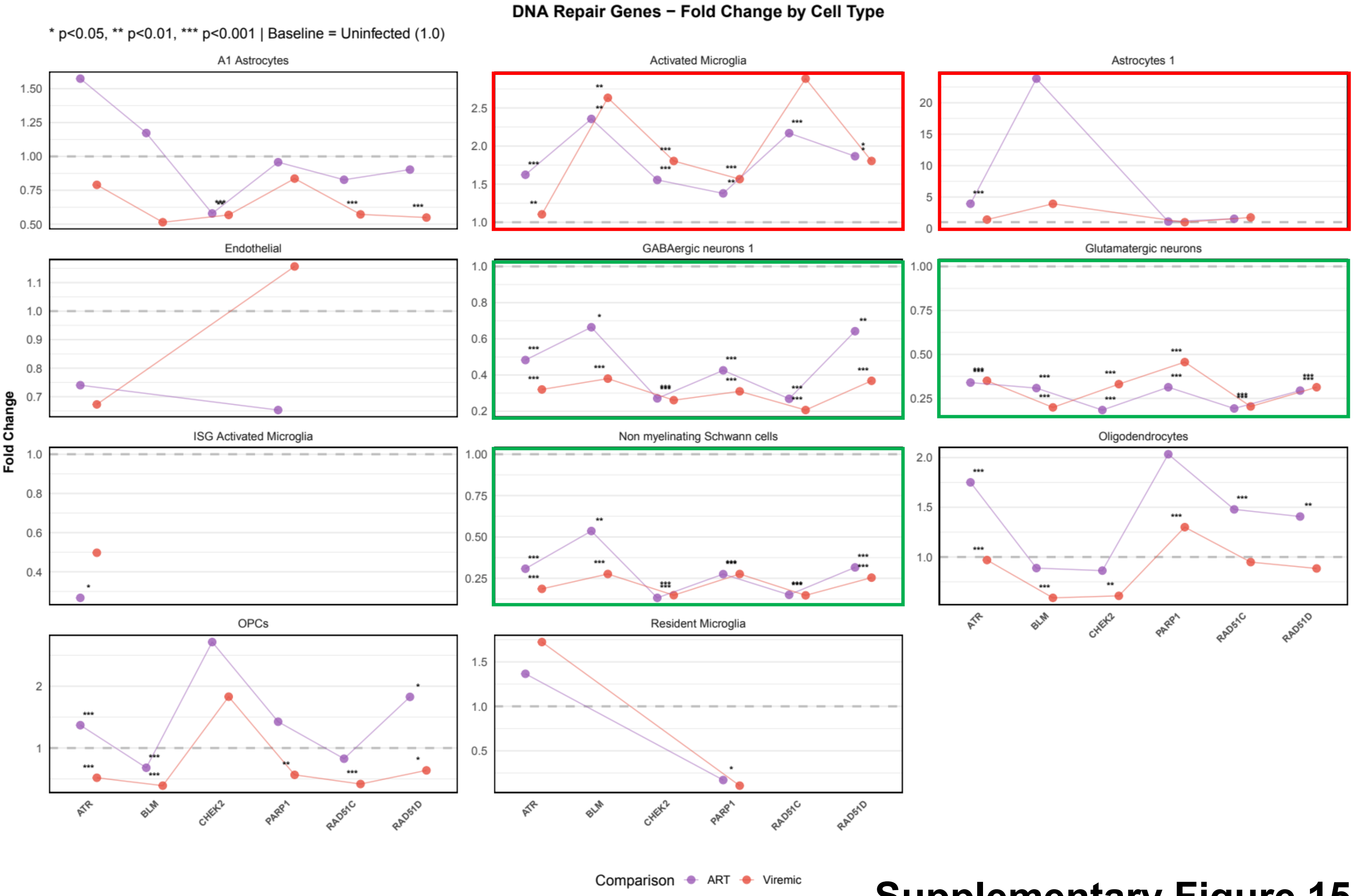


**Supplementary Figure 15**

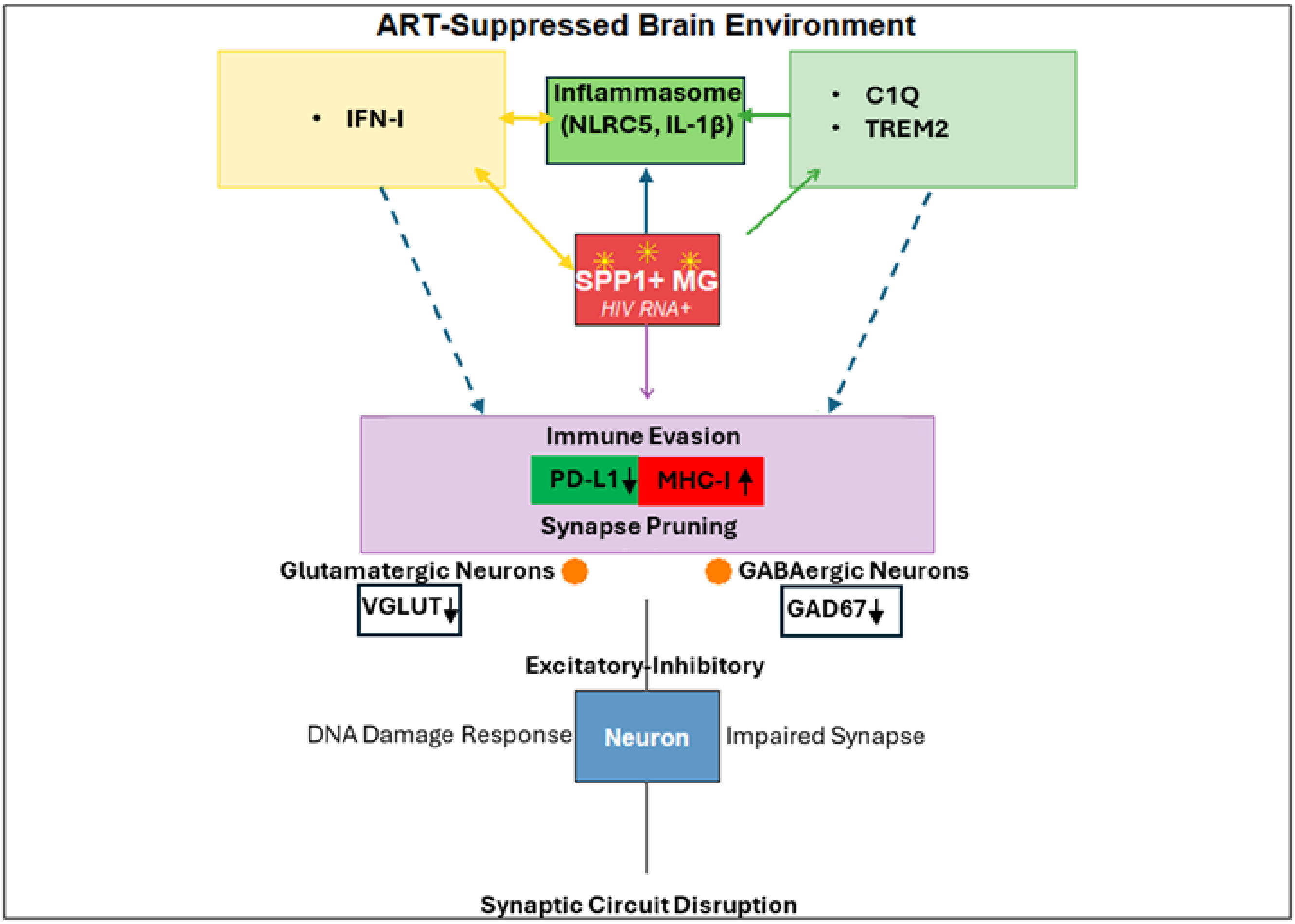


**Supplementary Figure 16**

#### Supplementary Table 1. Characterization of participants in this study

| PID | Age/Sex /Race | Years since HIV diagnosis | Terminal disease | Last CD4 before death (count/Date) | Last VL before death (Value/Date) | Last ART | Stopped ART before death? |
|---|---|---|---|---|---|---|---|
| **ART suppressed** | | | | | | | |
| Donor 1 | 48/M/W | 23 | metastatic anal cancer, Kaposi Sarcoma | 87/6 | <30/6 | FTC/TAF/BIC | No |
| Donor 2[a] | 65/F/W | - | Respiratory Arrest/Pulmonary Edema | -/- | -/-[b] | BIC/TAF/FTC | 7 days before death[c] |
| Donor 3 | 64/M/W | 27 | Esophageal and hepatic cancer | 162/131 | <30/30 | FTC/TAF+DTG | No |
| Donor 4 | 32/M/W | 10 | Disseminated multidrug resistant MAI, ESRD, cardiac arrest 2/2 hyperkalemia | 45/1 | <30/1 | FTC/TAF+DTG | No |
| Donor 2 | 39/-/- | N/A | | -/- | 40/0 | - | No |
| **ART interruption** | | | | | | | |
| Donor 6[d] | 47/M/W | 3 | MND | -/- | >7.5e5/>30 | 3TC/D4T/NVP/HU/KTA | 90 days before death |
| Donor 7 | 64/M/B | 36 | HCC, multiple comorbidities | 100/2 | 6.80E+04/1 | DRV/C/FTC/TAF | 111 days before death |
| Donor 8 | 44/F/W | 2 | Progressive multifocal leukoencephalopathy | 9/111 | 1.30E+06/25 | BIC/TAF/FTC | ≥215 days before death |
| **HIV negative** | | | | | | | |
| Control 1[d] | 56/F/W | N/A | - | N/A | N/A | N/A | N/A |
| Control 2[d] | 65/M/W | N/A | - | N/A | N/A | N/A | N/A |
| Control 3 | 70/F/W | N/A | Small cell lung CA, COPD, emphysema | N/A | N/A | N/A | N/A |

[a] Participant enrolled in NDRI. [b] The PWH was on ART throughout the study. While plasma HIV VL was lacking, the brain microglia HIV was latently infected, evidenced by a low gag RNA/DNA ratio in purified MG (***ref. 4***) and low Nef transcript detection by sc/snRNA-seq in activated MG (***Extended Fig.10A***). [C] Stopped ART 7 days before death with HIV latent infection in the brain MG. [d] Participants enrolled in NNTC. All the other participants were recruited through the "Last Gift" cohort. Drug abbreviations: FTC, Emtricitabine; TAF, Tenofovir Alafenamide; DTG, dolutegravir. N/A, data not available. Sex abbreviations: M, male; F, female.

**Supplementary Table 2: Summary of CellChat-Predicted Interactions Across Conditions**

| Condition | Total Interactions |
|---|---|
| Uninfected | 4,162 |
| Viremic | 4,574 |
| ART_Suppressed | 7,070 |

Total Interactions = sum of all predicted ligand-receptor interactions per condition. Mean Outgoing/Incoming = average signaling strength per cell type. ART-suppressed shows a 70% increase in total communication events compared to uninfected (7,070 vs 4,162).

#### Supplementary Table 3: Signaling Balance Across Cell Types and Conditions

| Condition | Cell Type | Outgoing | Incoming | Balance | Role |
|---|---|---|---|---|---|
| ART_Suppressed | Neurons | 586 | 564 | 22 | Net Sender |
| | Glutamatergic neurons | 447 | 550 | -103 | Net Receiver |
| | A1 Astrocytes | 489 | 500 | -11 | Net Receiver |
| | Non-myelinating Schwann cells | 465 | 500 | -35 | Net Receiver |
| | OPCs | 377 | 350 | 27 | Net Sender |
| | GABAergic neurons 1 | 327 | 375 | -48 | Net Receiver |
| | Oligodendrocytes | 251 | 202 | 49 | Net Sender |
| | Astrocytes 1 | 216 | 188 | 28 | Net Sender |
| | Endothelial | 239 | 161 | 78 | Net Sender |
| | Activated Microglia | 70 | 110 | -40 | Net Receiver |
| | ISG Activated Microglia | 45 | 33 | 12 | Net Sender |
| | Resident Microglia | 23 | 2 | 21 | Net Sender |
| Uninfected | Non-myelinating Schwann cells | 374 | 422 | -48 | Net Receiver |
| | Glutamatergic neurons | 375 | 386 | -11 | Net Receiver |
| | A1 Astrocytes | 312 | 332 | -20 | Net Receiver |
| | Endothelial | 320 | 239 | 81 | Net Sender |
| | GABAergic neurons 1 | 212 | 241 | -29 | Net Receiver |
| | OPCs | 221 | 186 | 35 | Net Sender |
| | Astrocytes 1 | 144 | 122 | 22 | Net Sender |
| | Oligodendrocytes | 76 | 83 | -7 | Net Receiver |
| | Activated Microglia | 47 | 70 | -23 | Net Receiver |
| | ISG Activated Microglia | — | — | — | No interactions |
| | Resident Microglia | — | — | — | No interactions |
| Viremic | Glutamatergic neurons | 538 | 483 | 55 | Net Sender |
| | Non-myelinating Schwann cells | 366 | 375 | -9 | Net Receiver |
| | A1 Astrocytes | 311 | 329 | -18 | Net Receiver |
| | GABAergic neurons 1 | 205 | 269 | -64 | Net Receiver |
| | Endothelial | 254 | 175 | 79 | Net Sender |
| | OPCs | 191 | 231 | -40 | Net Receiver |
| | Oligodendrocytes | 190 | 159 | 31 | Net Sender |
| | Astrocytes 1 | 82 | 115 | -33 | Net Receiver |
| | Activated Microglia | 77 | 113 | -36 | Net Receiver |
| | ISG Activated Microglia | 53 | 37 | 16 | Net Sender |
| | Resident Microglia | 20 | 1 | 19 | Net Sender |

Outgoing = signaling sent by cell type; Incoming = signaling received by cell type; Balance = Outgoing - Incoming. Positive balance indicates net sender; negative balance indicates net receiver. Activated microglia are consistently net receivers across all conditions (balance: -23 to -40), while ISG-activated microglia shift to net senders under inflammatory conditions (+16 in viremic, +12 in ART-suppressed). The "--" indicates no predicted interactions.

#### Supplementary Table 4: Key Pathway Communication Probabilities Across Conditions

| Pathway | Uninfected | Viremic | ART_Suppressed | Fold Change (ART vs Uninfected) |
|---|---|---|---|---|
| SPP1 | 0.46 | 4.18 | 2.91 | 6.3× |
| NCAM | 14.69 | 7.22 | 27.12 | 1.85× |
| NRXN | 16.57 | 15.52 | 27.08 | 1.63× |
| NRG | 14.44 | 15.92 | 24.23 | 1.68× |
| LAMININ | 3.42 | 3.47 | 5.91 | 1.73× |
| CADM | 5.99 | 4.32 | 7.33 | 1.23× |
| NEGR | 5.00 | 2.48 | 5.58 | 1.12× |
| EPHA | 4.65 | 1.98 | 3.81 | 0.82× |
| CDH | 3.02 | 2.07 | 3.11 | 1.03× |
| SEMA6 | 1.24 | 1.78 | 2.64 | 2.13× |
| NGL | 1.23 | 1.78 | 2.59 | 2.10× |
| COLLAGEN | 0.66 | 2.45 | 1.77 | 2.68× |
| PTN | 0.32 | 0 | 1.51 | 4.75× |
| BMP | 0.65 | 0 | 1.38 | 2.14× |
| VISFATIN | 0.29 | 1.33 | 1.02 | 3.51× |
| TENASCIN | 0.35 | 1.14 | 1.21 | 3.45× |
| JAM | 0.00 | 0.65 | 1.11 | — |
| FGF | 0.35 | 0.89 | 0.85 | 2.39× |

Values represent pathway-level communication probabilities from CellChat analysis. Higher scores indicate stronger predicted signaling. Bold = highest value per pathway across conditions. SPP1 shows a 9-fold increase during viremia (4.18 vs 0.46) and 6-fold increase under ART (2.91 vs 0.46) relative to uninfected. Neuronal pathways (NCAM, NRXN, NRG) are enhanced 1.6-1.9× under ART compared to uninfected.

**Supplementary Table 5: SPP1 Signaling Sources and Receivers in ART-Suppressed Brains**

| Role | Cell Type | Predicted Interactions |
|---|---|---|
| Senders | Activated Microglia | 53 |
| | Oligodendrocytes | 52 |
| | ISG Activated Microglia | 42 |
| | Resident Microglia | 35 |
| | Astrocytes | 11 |
| Receivers | Glutamatergic neurons | 37 |
| | A1 Astrocytes | 36 |
| | GABAergic neurons 1 | 24 |
| | OPCs | 22 |
| | Non-myelinating Schwann cells | 20 |
| | Endothelial | 17 |
| | Astrocytes 1 | 12 |
| | Neurons | 11 |
| | Oligodendrocytes | 8 |

SPP1 source cell analysis reveals that activated microglia are the dominant cellular source of SPP1 signaling in ART-suppressed brains (53 predicted interactions), followed by oligodendrocytes (52 interactions). The primary receivers of SPP1 signals are glutamatergic neurons (37 interactions) and A1 astrocytes (36 interactions). This positions activated microglia as key drivers of SPP1-mediated signaling to neurons and astrocytes in the ART-suppressed brain.

**Supplementary Table 6: All Diseases & Functions by Association Score of HIV$^+$ and HIV$^-$ Cells**

| HIV Status | Score | Focus Molecules | Top Diseases and Functions |
|---|---|---|---|
| **HIV+** | **68** | **31** | **Immunological Disease, Inflammatory Disease, Inflammatory Response** |
| **HIV+** | **62** | **29** | **Developmental Disorder, Hereditary Disorder, Neurological Disease** |
| **HIV+** | **59** | **28** | **Connective Tissue Disorders, Developmental Disorder, Hereditary Disorder** |
| HIV+ | 45 | 23 | Molecular Transport, Developmental Disorder, Organismal Injury and Abnormalities |
| HIV+ | 40 | 21 | Developmental Disorder, Hereditary Disorder, Metabolic Disease |
| **HIV-** | **32** | **10** | **Cell-To-Cell Signaling and Interaction, Cell Cycle, Cancer** |
| HIV+ | 26 | 15 | Cardiovascular Disease, Organismal Injury and Abnormalities, Renal and Urological Disease |
| HIV+ | 21 | 13 | Cell Morphology, Connective Tissue Development and Function, Embryonic Development |
| HIV+ | 19 | 12 | Cell Death and Survival, Organismal Injury and Abnormalities, Lipid Metabolism |
| HIV+ | 19 | 12 | Cellular Function and Maintenance, Gene Expression, Protein Synthesis |
| HIV+ | 19 | 12 | Connective Tissue Development and Function, Organ Morphology, Reproductive System Development and Function |
| HIV+ | 19 | 12 | Developmental Disorder, Hereditary Disorder, Organismal Injury and Abnormalities |
| HIV+ | 15 | 10 | Cell Death and Survival, Cellular Development, Cellular Growth and Proliferation |
| HIV+ | 13 | 9 | Digestive System Development and Function, Nutritional Disease, Organismal Injury and Abnormalities |
| HIV+ | 6 | 5 | Embryonic Development, Organ Development, Organismal Development |
| HIV+ | 2 | 1 | Cell Cycle, Cellular Development, Cellular Growth and Proliferation |

**Supplementary Table 7. Significantly upregulated genes, including MHC-I pathway genes, antigen-processing components, and immune regulators, in IFN-β-treated MG, compared with IFNβ-untreated control MG**

| Symbol | Log FC | Symbol | Log FC | Symbol | Log FC | Symbol | Log FC | Symbol | Log FC |
|---|---|---|---|---|---|---|---|---|---|
| ABCA13 | 1.05 | DDX60 | 1.98 | KRT33B | 1.54 | PSMB8-AS1 | 1.3 | VIT | 1.73 |
| ABCA8 | 1.35 | DDX60L | 2.15 | KRTAP1-5 | 1.16 | PSMB9 | 1.56 | VMO1 | 1.36 |
| AC0099502 | 2.46 | DHX58 | 2.57 | LAG3 | 1.65 | PTCSC2 | 1.45 | XAF1 | 3.64 |
| ACP5 | 1.12 | DTX3L | 2.05 | LAMA3 | 1.55 | PTN | 2.58 | XXbac-BPG248L24.12 | 1.08 |
| ACSL5 | 2.82 | DUSP2 | 1.3 | LAP3 | 1.44 | PTPRB | 1.66 | ZNFX1 | 1.01 |
| ACSM5 | 1.65 | EBI3 | 1.28 | LCP2 | 1.44 | PTPRQ | 2.42 | ZSCAN23 | 1.22 |
| ACTN2 | 2.45 | ECM2 | 1.09 | LGALS3BP | 1.86 | RAB17 | 1.17 | CTD_2095E45 | 1.32 |
| ADAR | 1.1 | EHF | 1.66 | LGALS9 | 3.15 | RAPGEF5 | 1.94 | CTSS | 2.35 |
| ADGRB1 | 1.7 | EIF2AK2 | 1.71 | LGI4 | 1.02 | RASGRP3 | 1.06 | CXCL10 | 1.87 |
| AF1312171 | 1.28 | EPGN | 1.05 | LGR5 | 1.41 | REC8 | 1.33 | CXCL11 | 2.45 |
| AGMO | 1.44 | EPSTI1 | 3.5 | LINC00968 | 1.01 | RET | 1.01 | CYP1A1 | 1.01 |
| ANGPTL1 | 2.19 | ETV7 | 1.69 | LINC01085 | 1.31 | RGS16 | 1.04 | CYTIP | 1.02 |
| ANKRD34B | 1.52 | ETV7-AS1 | 1.01 | LINC01091 | 1.5 | RIGI | 2.06 | DARS1-AS1 | 1.02 |
| ANO3 | 1.39 | F3 | 1.34 | LINC01444 | 1.15 | RMDN2-AS1 | 1.95 | DDO | 1.38 |
| ANO7L1 | 1.23 | FAM167A | 1.66 | LINC02015 | 1.03 | RNF213 | 1.78 | IRF9 | 1.21 |
| ANXA3 | 1.19 | FBXO39 | 4.79 | LINC02112 | 1.17 | RP11-468E2.4 | 1.07 | ISG15 | 3.18 |
| APLN | 1.8 | FBXO6 | 1.34 | LINC02154 | 2.28 | RP11_134L101 | 1.83 | KCNK2 | 1.2 |
| APOBEC3D | 1.36 | FGL2 | 1.15 | LINC02574 | 4.79 | RP11_284F2110 | 1.23 | KCNT2 | 1.3 |
| APOBEC3G | 1.1 | FIBCD1 | 1.27 | LINC02941 | 1.39 | RP11_288L94 | 4.73 | KCTD4 | 1.15 |
| APOL1 | 1.97 | FNDC1 | 1.13 | LINC03013 | 1.35 | RP11_532F63 | 1.75 | KIAA1755 | 1.51 |
| APOL3 | 1.68 | FRMPD4 | 1.09 | LIPC | 1.74 | RP11_849N153 | 1.12 | KIF6 | 1.33 |
| APOL6 | 1.6 | GALNT15 | 1.76 | LMO2 | 1.46 | RP4_730K33 | 1.29 | KMO | 1.22 |
| APOLD1 | 1.08 | GALR1 | 1.04 | LOC100128770 | 1.27 | RSAD2 | 5.8 | PLSCR1 | 2.92 |
| ARC | 1.27 | GBP1 | 1.73 | LOC105374235 | 1.01 | RTP4 | 3.23 | PML | 1.03 |
| ASPA | 1.01 | GBP1P1 | 4.16 | LY6E | 3.06 | SAMD9 | 2.7 | PNPT1 | 1.09 |
| ATP8B4 | 1.02 | GBP4 | 2.7 | LY6E-DT | 1.97 | SAMD9L | 2 | PPFIA4 | 1.19 |
| ATP8B5P | 1.05 | GHDC | 1.11 | MAPK8IP2 | 1.53 | SAMHD1 | 2.21 | PRAL | 4.27 |
| B2M | 1.29 | GIMAP2 | 1.53 | MDK | 2.58 | SECTM1 | 1.58 | PSG1 | 1.36 |
| B3GNT3 | 1.62 | GKN1 | 1.23 | MEI1 | 1.01 | SEMA3A | 1.2 | PSG2 | 1.48 |
| BATF2 | 3.41 | GMPR | 1.03 | MIR133A1HG | 1 | SEMA6D | 1.09 | PSMB8 | 1.01 |
| BCAN-AS1 | 1.89 | GPR65 | 1.05 | MKX | 1.11 | SERPINB2 | 1.3 | TRIM69 | 2.22 |
| BEX1 | 1.18 | GRIA2 | 1.38 | MLKL | 1.13 | SERPING1 | 2.68 | TSPAN8 | 1.09 |
| BICDL2 | 1.67 | GRIP2 | 3.58 | MOV10 | 1.1 | SFRP1 | 1.2 | TXNIP | 1.31 |
| BISPR | 3.29 | HAS2 | 1.5 | MS4A4A | 1.25 | SHFL | 1.44 | TYMP | 2.86 |
| BRCA2 | 1.05 | HCG4P11 | 1.96 | MS4A7 | 1.38 | SLC15A3 | 2.31 | UBA7 | 1.78 |
| BRIP1 | 1.08 | HCG4P5 | 1.57 | MSTN | 1.34 | SLC16A4 | 1.23 | UBE2L6 | 1.96 |
| BST2 | 5.96 | HCP5 | 2.35 | MX1 | 5.74 | SLC16A6 | 1.12 | UBQLNL | 1.52 |
| BTC | 1.31 | HELZ2 | 2.77 | MX2 | 5.89 | SLC18A3 | 1.46 | UNC93B1 | 1.38 |
| BTN3A1 | 1.33 | HERC5 | 2.56 | MYCT1 | 1.15 | SLC6A1 | 1.27 | USP18 | 3.16 |
| BTN3A2 | 1.31 | HERC6 | 3.36 | NBEAP1 | 1.01 | SLC7A2 | 1.06 | CMPK2 | 5.73 |
| BTN3A3 | 1.43 | HLA-A | 1.68 | NCKAP5 | 1.62 | SLC8A2 | 1.43 | CRYBG1 | 1.13 |
| C15orf48 | 1.03 | HLA-B | 2.1 | NFE2L3 | 1.24 | SMTNL1 | 3 | CT69 | 1.24 |
| C1R | 2.59 | HLA-C | 1.79 | NLRC5 | 1.25 | SOCS1 | 1.66 | CTA_212A21 | 1.91 |
| C1S | 2.26 | HLA-E | 1.17 | NLRP10 | 1.56 | SP110 | 2.07 | CTA_384D836 | 1.11 |
| C2 | 1.28 | HLA-F | 2.22 | NMI | 1.1 | SPEF1 | 1.2 | IGFBP1 | 1.7 |
| C6orf62 | 1.22 | HLA-G | 1.8 | NOD2 | 2.05 | SRPX2 | 1.07 | IL12RB1 | 1.61 |
| CABP1 | 1.07 | HLA-H | 1.57 | NPTX1 | 1.22 | SSPOP | 1.2 | IL18BP | 1.29 |
| CAMK2B | 1.76 | HLA-J | 2.02 | NTRK3 | 1.1 | STAT1 | 1.06 | IL23A | 1.52 |
| CARD16 | 1.48 | HLA-L | 1.11 | NXNL2 | 1 | STAT2 | 1.35 | IRF7 | 2.39 |
| CARINH | 1.17 | HLF | 1.01 | OAS1 | 6.48 | STKLD1 | 1 | PKD1P3 | 1.15 |
| CASP1 | 1.55 | HTR2B | 1.23 | OAS2 | 4.28 | SUCNR1 | 1.24 | PLAAT4 | 1.49 |
| CASP10 | 1.77 | IFI16 | 1.05 | OAS3 | 4.41 | SUGCT-AS1 | 1.34 | PLEKHA4 | 1.23 |
| CCL2 | 1.23 | IFI27 | 3.47 | OASL | 2.35 | SULT1A1 | 1.19 | PLEKHG4 | 1.12 |
| CCL5 | 1.6 | IFI30 | 1.98 | OGFR-AS1 | 1.47 | TAP1 | 1.04 | PLPPR4 | 1.23 |
| CD274/PD-L1 | 1.23 | IFI35 | 2.75 | OLFML1 | 1.48 | TAS2R1 | 1.3 | TRANK1 | 2.38 |
| CD36 | 2.17 | IFI44 | 3.24 | PAMR1 | 1.28 | TEK | 1.38 | TRIM14 | 2.94 |
| CD68 | 1.43 | IFI44L | 6.97 | PARP10 | 1.29 | TEKT2 | 1.41 | TRIM21 | 1.31 |
| CD7 | 1.77 | IFI6 | 5.42 | PARP12 | 1.71 | TENT5A | 1.42 | TRIM25 | 1.77 |
| CELSR3 | 1.11 | IFIH1 | 2.85 | PARP14 | 1.69 | THEMIS2 | 2.06 | TRIM34 | 1.02 |
| CFAP54 | 1.42 | IFIT1 | 5.28 | PARP9 | 2.73 | THSD7B | 1.1 | CLEC3B | 3.01 |
| CFAP57 | 1.1 | IFIT2 | 2.53 | PATL2 | 1.8 | TLDC2 | 1.93 | IFITM3P2 | 1.8 |
| CFAP95 | 1.7 | IFIT3 | 3.52 | PDE2A | 1.14 | TLR3 | 1.6 | PKD1P1 | 1.07 |
| CFB | 2.03 | IFIT5 | 1.11 | PDE4C | 1.02 | TMEM140 | 1.75 | TNFSF18 | 1.92 |
| CHAT | 1.11 | IFITM1 | 6.03 | PDGFB | 1.54 | TMEM156 | 1.44 | PIWIL4 | 1.39 |
| CHRM2 | 1.14 | IFITM2 | 1.15 | PDZD2 | 1.83 | TNFSF13B | 2.62 | TNFSF15 | 1.88 |
| CIMAP1B | 3.01 | IFITM3 | 1.95 | | | | | | |
